# Scale-mixture Langevin sampling in the subspaces of recurrent cortical circuit dynamics

**DOI:** 10.64898/2026.09.28.754844

**Authors:** Zimei Chen, Yi Ren, Wen-Hao Zhang

## Abstract

Recurrent cortical circuits are hypothesized to perform Bayesian posterior sampling through their stochastic dynamics. Yet identifying *which* sampling algorithm a circuit intrinsically implements, as opposed to mapping manually designing algorithms onto circuits, requires rigorous theoretical analysis of the biological circuit dynamics, and remains a fundamental open challenge because of the analytical intractability of nonlinear recurrent dynamics. Here, we tackle this challenge through comprehensive theoretical analysis of analytically tractable nonlinear recurrent circuit dynamics used in neuroscience research— the continuous attractor network (CAN). Surprisingly, we find that the circuit, without any non-Gaussian components in its connectivity or internal variability, intrinsically emerges heavy-tailed subspaces dynamics described as scale-mixture Langevin posterior sampling. The mechanism emerges from the circuit’s own nonlinear dynamics: divisive normalization renders the overall population activation Gamma-distributed which multiplicatively modulates the time constant of the Langevin sampling in the stimulus feature subspace. Eventually, this scale-mixture structure produces a heavy-tailed (Student-*t*) sampling step size distribution, significantly accelerating the sampling speed. Moreover, the scale-mixture Langevin sampling approaches a Lévy-like regime in the weak input limit. Our results analytically identify an intrinsic circuit algorithm embedded in subspaces of nonlinear recurrent dynamics, and establish a minimal circuit mechanism for endogenous heavy-tail statistics without any explicit non-Gaussian components.

## 1 Introduction

A wide spectrum of perceptual and cognitive processes have been mathematically formulated as Bayesian inference [1], including visual perception [2], multisensory integration [3], sensorimotor control [4], and decision-making [5], establishing the “Bayesian brain” framework. A fundamental question is how neural circuits mechanistically implement this inference [6–8]. Among the proposed mechanisms, sampling-based Bayesian inference has attracted particular attention [7, 9–12]: the prominent variability of cortical responses is hypothesized to be a functional signature of stochastic posterior sampling [10–16]. Progress has been made in identifying specific sampling algorithms in recurrent circuits, including Gibbs sampling [12], Langevin sampling [17–19], Hamiltonian sampling [10, 11, 16, 20], and natural-gradient sampling [21, 22], etc.

Yet a conspicuous mismatch exists between these identified algorithms and empirical observations of circuit properties. Most theoretically identified circuit sampling algorithms assume Gaussian dynamics to gain analytical tractability (e.g., [9, 10, 12, 13, 16, 17]), whereas a growing body of evidence reveals that neural circuit properties are systematically *heavy-tailed*, including synaptic weights [23, 24], single neurons’ firing rate [25–27], population response geometry [28], and hippocampal replay sequences during sharp-wave ripples [29, 30], etc. These suggest that heavy-tailed structure is not an incidental feature of neural circuits, but probably has functional significance.

A growing body of theoretical work has investigated heavy-tailed circuit dynamics and their roles in Bayesian computation. To generate heavy-tailed neuronal responses, the predominant strategy *directly* imports non-Gaussian components into the circuit dynamics—using heavy-tailed (log-normal) recurrent synaptic weights [31–33] or explicitly injecting Lévy-stable noise into the dynamics [34, 35]. In such models, heavy-tailed circuit responses are largely *inherited* from explicitly non-Gaussian connectivity and/or variability, and it is thus not surprising that these networks produce heavy-tailed responses. A more fundamental question remains unaddressed: <u>can heavy-tailed structure emerge from a network without explicit non-Gaussian components, arising purely from the circuit’s nonlinear recurrent dynamics? Furthermore, how does such emergent heavy-tailed structure benefit Bayesian computations?</u> Answering this requires rigorous analysis of the nonlinear stochastic circuit dynamics, while it remains generally challenging due to the analytical intractability.

The present study answers this question affirmatively through rigorous theoretical analysis of a *minimal*, analytically tractable recurrent circuit model–a continuous attractor network (CAN) widely used in neuroscience [36–39]. Surprisingly, our analysis reveals that the CAN *without* any heavy-tailed components collectively generates heavy-tailed structure in its subspaces satisfying the **scale-mixture Langevin posterior sampling**, an advanced sampling algorithm developed in ML with heavy-tailed step-size structure [40]. Notably, the heavy-tailed structure resides in the *sampling dynamics* with the step-size distribution being heavy-tailed (Student-*t*), while the *equilibrium distribution* of the sampling still converges to Gaussian posteriors. The circuit mechanism emerges in four steps:

- The bump-profile circuit dynamics has two dominant subspaces: the bump **position** (representing the stimulus feature) and the bump **height** (proportional to overall population activation) (Fig. 2B).
- Due to the nonlinear recurrent dynamics, the two subspaces are **coupled**: the bump height **multiplicatively** modulates the Langevin time constant of the position subspace (Fig. 2C).
- The position subspace executes Langevin sampling of the Gaussian stimulus posterior; the bump height satisfies a **Gamma** distribution, arising from divisive normalization (Fig. 2E), 3A).
- The Gamma-distributed height acts as the scale “mixer” for the position step size, yielding a heavy-tailed Student-*t* step-size distribution—implementing scale-mixture Langevin sampling.

The above emergent structure, including Gamma scale mixer, Gaussian conditional, Student-*t* marginal, is identical to the *Gaussian scale mixture* (GSM) model [41–45] widely used in heavy-tailed natural image statistics [46–49], suggesting a deep structural alignment between the circuit’s intrinsic dynamics and the natural statistics. This emergent scale-mixture structure significantly accelerates sampling speed relative to the sampling with the same mean step size (Fig. 2G), and even admits a Lévy-like regime in the weak input limit (Fig. 3B). Simulations confirm our theory, and reveal the robustness of the scale-mixture structure across a wide range of circuit parameters.

### Significance and contribution

**1)** A **minimal** circuit mechanism for the endogenous emergence of heavy-tailed sampling without any explicit non-Gaussian components in the circuit dynamics. **2)** The first mechanistic, analytical understanding of how the GSM structure emerges from nonlinear recurrent circuit dynamics. **3)** The first study analytically reveals the scale-mixture Langevin sampling in the circuit’s subspace, deepening our understanding of advanced circuit algorithms.

## 2 The canonical recurrent circuit model

We consider a canonical recurrent circuit model based on continuous attractor networks [16, 36–38] or supralinear stabilization network [11, 50], which are widely used in neuroscience research and can be regarded as a cortical hypercolumn. Briefly, the circuit consists of excitatory (E) neurons and a pool of global inhibitory (I) neurons (Fig. 1). The E neurons tune to stimulus and are recurrently connected, while I neurons are not tuned for simplicity (details in Appendix B).

**Figure 1:**
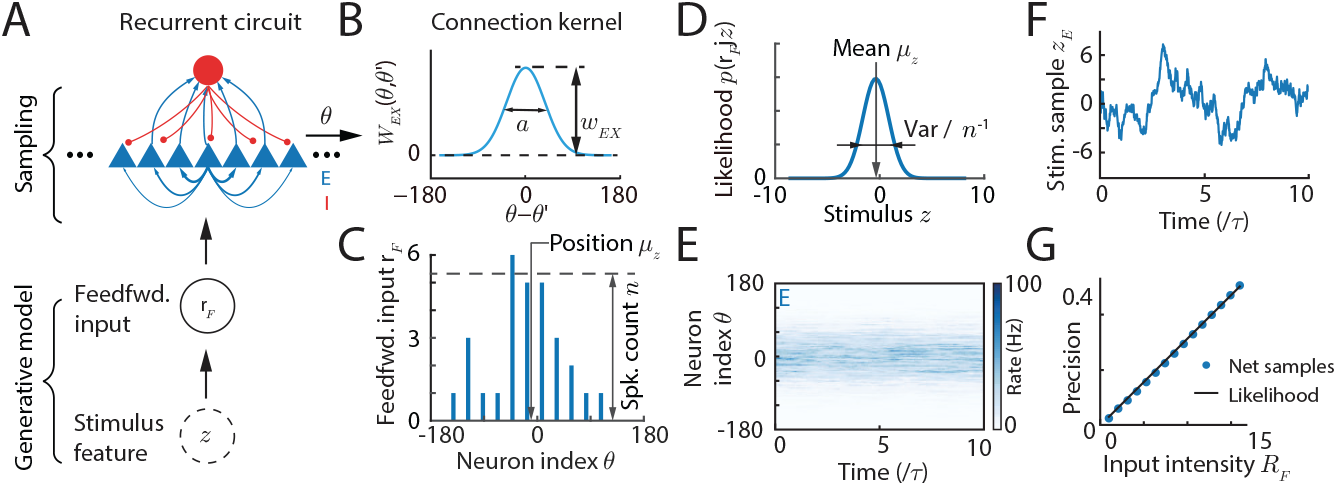
(A) The recurrent circuit of E neurons and inhibitory neurons and sampling-based Bayesian inference. (B) The recurrent and feedforward connection kernel between E neurons. (C) Feedforward (spiking) input with Gaussian tuning across the stimulus feature. In simulation, it is approximated as Gaussian random variables with multiplicative noise. (D) The stimulus feature likelihood given a feedforward input. (E) E population responses. (F) Stimulus feature samples from E neurons. (G) The circuit with fixed weights flexibly samples posteriors with different uncertainties in the equilibrium.

**Figure 2:**
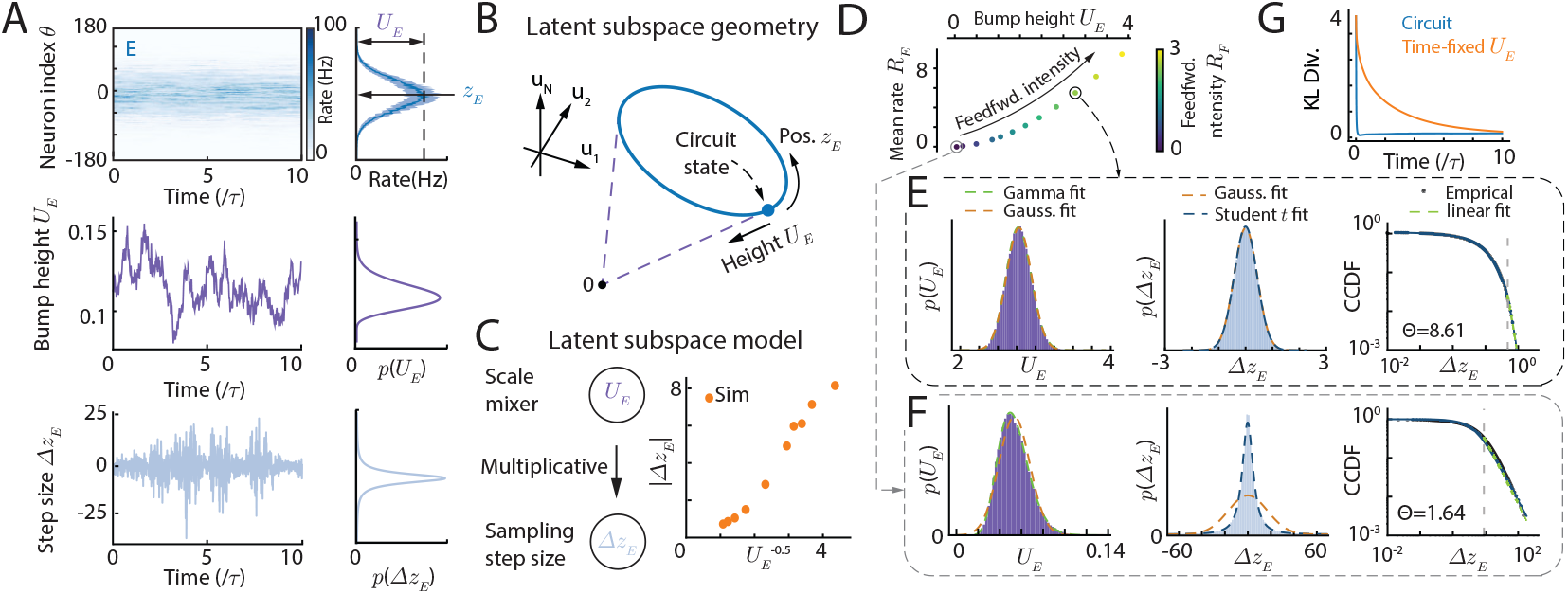
Scale-mixture Langevin sampling dynamics in circuit subspaces. (A) E population responses and the trajectories of bump height *U*_*E*_ and bump position step size Δ*z*_*E*_ linearly read out from instant E responses. (B-C) The latent subspace geometry and model from theoretical analysis. The position and height subspaces are intertwined, where the height *U*_*E*_ multiplicatively modulates the position step size Δ*z*_*E*_. (D-F) The heavy-tailed sampling step size emerges at the limit of low circuit activation. (D): The population synaptic input height and population firing rate height with changing feedforward input intensity. (E-F): The distributions of bump height (left) and the sampling step size (middle), and the complementary cumulative distribution function of the step size (right). (G) The heavy-tailed step sizes accelerate sampling speed compared with the sampling with time-fixed *U*_*E*_.

**Figure 3:**
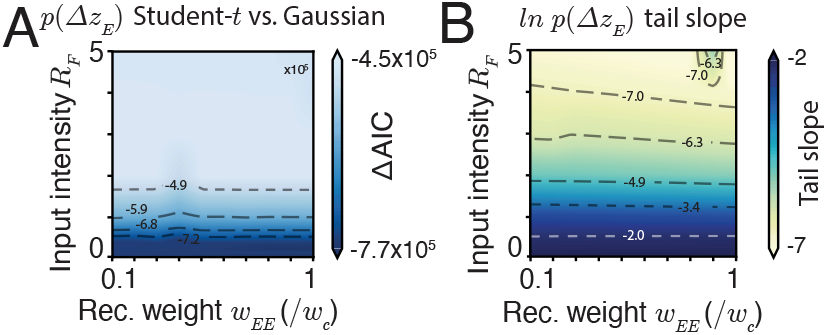
(A) The comparison of student-*t* vs Gaussian fit the sampling step size distribution *p*(Δ*z*_*E*_). A negative Δ*AIC* means student-*t* provides a better fit. (B) The tail slope of ln *p*(Δ*z*), corresponding to the DoF *v* in Eq. (15).

### E neurons’ dynamics

The E neurons in the circuit are selective for a 1D stimulus feature *z*, e.g., spatial location. Denote *θ*_*j*_ as the preferred stimulus feature of the *j*-th E neuron, and the preferred feature of all *N*_*E*_ E neurons, 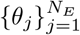 uniformly cover the entire stimulus feature space *z*, which is widely considered in neuroscience research. Mathematically, in the continuum limit (*θ*_*j*_ → *θ*) corresponding to an infinite number of neurons, the dynamics of E neurons is,

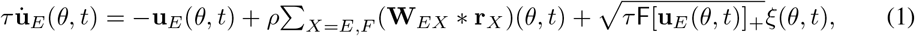

where **u**_*E*_(*θ, t*) and **r**_*E*_(*θ, t*) represent, respectively, the synaptic inputs and firing rates of neurons preferring *z* = *θ. X* = *E, F* denotes E neurons and feedforward sensory. *τ* is the time constant, *ρ* = *N/*2*π* is the neuronal density covering the stimulus feature space, and [*x*]_+_ = max(*x*, 0). F is the Fano factor characterizing the internal Poisson variability mimicking spike generation.

### Recurrent connection kernel

**W**_*Y X*_ (*θ*) is the recurrent connection kernel from neurons with type *X* to those with type *Y*, which are modeled as Gaussian functions in the model (Fig. 1B),

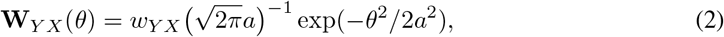

where the scalar *w*_*Y X*_ is the peak recurrent weight strength and is the only circuit parameter to be adjusted for realizing Bayesian sampling. and *a* the connection width across the stimulus feature space. The symbol ∗ denotes the convolution, i.e., **W**(*θ*) ∗ **r**(*θ*) = **W**(*θ* − *θ*^*′*^)**r**(*θ*^*′*^)*dθ*^*′*^.

### Divisive normalization from inhibitory neurons

I neurons are driven by E neurons and then provide global, unstructured inhibitory feedback. To facilitate the math analysis to obtain the circuit algorithm, we don’t explicitly model I neurons’ dynamics but absorb their effect into the divisive normalization that acts as E neurons’ activation function, a canonical cortical operation [51–54].

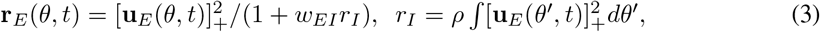

where *r*_*I*_ (scalar) corresponds to the mean firing rate of the I neurons. *w*_*EI*_ (a positive scalar) characterizes the global inhibition strength.

### Stochastic sensory feedforward input

The feedforward input **r**_*F*_ (*θ, t*) (Eq. 1) is stochastically evoked by an external stimulus *z*, and is modeled as conditionally independent Poisson spikes with Gaussian tuning given a stimulus *z* (Fig. 1C), consistent with neural coding models (e.g., [6, 12]).

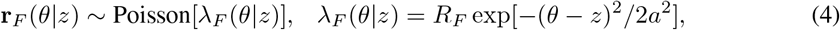

where *λ*_*F*_ (*θ*|*z*) is the mean firing rate. In simulating the rate-based circuit dynamics, **r**_*F*_ is approximated as a continuous Gaussian random variable with multiplicative noise.

### 2.1 The generative model and posteriors to be computed by the circuit

The stage from the external stimulus feature *z* to the feedforward input **r**_*F*_ is regarded as the generative process of the circuit’s observation (Fig. 1A). To focus on the mechanistic understanding of the scale-mixture structure in the nonlinear recurrent dynamics, we consider a uniform prior over the stimulus feature *z*, as previous studies also suggested the recurrent circuits with translation-invariant connectivity also store an internal uniform prior [12]. Later we extend the circuit to the case of a non-uniform prior and show the scale-mixture structure still holds (Appendix G.1 and Fig. A2).

With the uniform prior, the stimulus posterior *π*(*z*) is proportional to the likelihood ℒ (*z*),

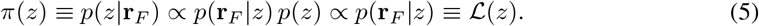

From Eq. 4, the stimulus likelihood is derived as a Gaussian distribution (Appendix B.4),

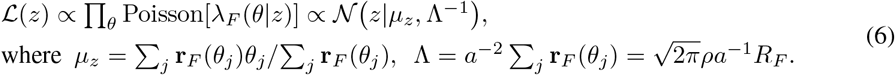

where 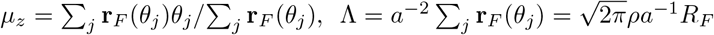.

Note that the Gaussian stimulus likelihood is not directly assigned, but resulted from the Gaussian profile of feedforward input tuning *λ*_*F*_ (*θ*| *z*) (Eq. 4) [6, 8]. Its mean *µ*_*z*_ and the precision Λ are regarded as **r**_*F*_ ‘s location in the stimulus feature space and height (total input spike count) (Fig. 1C-D), and can be read out from **r**_*F*_ via a linear decoder called population vector [55, 56],

## 3 Naive Langevin sampling in the circuits

### 3.1 Theoretical analysis of the circuit dynamics to understand circuit algorithm

Mechanistic understanding of the circuit’s algorithm requires rigorous theoretical analysis of the nonlinear recurrent dynamics, which is a major challenge in the field. The recurrent circuit model in the present study exists **analytical** solutions [38, 57], including attractor states, full eigenspectrum of the perturbation dynamics. This enables us to project the high-dimensional circuit dynamics onto the eigenmodes (subspaces), and then the subspace dynamics will be compared and linked to Bayesian sampling algorithms. Below, we briefly introduce the key steps and results (details in Appendix B).

### Attractors

E neurons in canonical circuit dynamics have the following attractor states with a bump profile over the stimulus feature space (Fig. 1E; Appendix B),

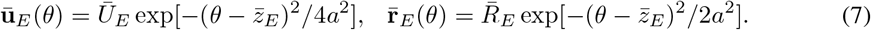

In contrast, I neurons don’t have a spatial bump profile since they are not tuned to stimulus (Eq. 3).

### Sampling dynamics on the stimulus feature subspace

The first dominant eigenmode of the circuit dynamics corresponds to the change of bump position *z*_*E*_ (Appendix B, [38, 57]). *z*_*E*_ is regarded as the circuit’s internal representation of stimulus feature. Projecting the E dynamics (Eq. 1) onto the position eigenvector (calculating the inner product of the circuit dynamics and the eigenvectors) yields the dynamics of *z*_*E*_ (details in Appendix B.3),

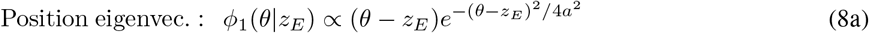

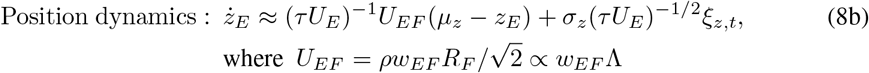

where *U*_*E*_ is the height of the bump attractor (Eq. 7) and determines the time constant of the *z*_*E*_ dynamics that will affect the average step size Δ*z*_*E*_ over time (Fig. 2C). *U*_*EF*_ characterizes the magnitude (height) of the received feedforward input, and is proportional to the likelihood precision Λ (Eq. 6). This is a key property to link the position subspace dynamics to the Langevin sampling that will be shown below. Moreover, 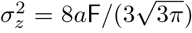 is a constant determining the strength of internal variability projected onto the position subspace, and doesn’t change with network activities.

### 3.2 Naive Langevin sampling in the circuits

We explore how the position dynamics (Eq. 8b) can sample the posterior *π*(*z*), i.e., its equilibrium distribution should converge to *π*(*z*). By utilizing the proportionality between *U*_*EF*_ and Λ (Eq. 8b), the position dynamics can be converted in conventional Langevin posterior dynamics [58, 59],

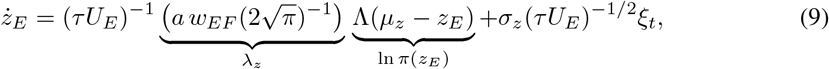

To match the equilibrium distribution of *z*_*E*_ with the posterior *π*(*z*), we only need to adjust the ratio between *λ*_*z*_ and 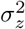 by setting the feedforward weight *w*_*EF*_ (Appendix Sec.B.5)

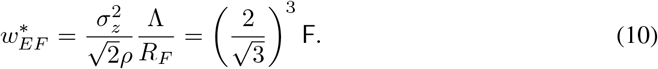

In addition, once the *w*_*EF*_ is set to the optimal value, the recurrent circuit with fixed synaptic weights can sample posteriors with different means and precisions (Fig. 1G), exhibiting significant flexibility.

## 4 The scale-mixture Langevin sampling in the circuit

The above analyses of the Langevin sampling (Eq. 8b) implicitly treats the bump height *U*_*E*_ as a constant, free parameter. In practice, *U*_*E*_ is also a stochastic variable fluctuating over time due to the internal variability (Fig. 2A). Since the *U*_*E*_ governs the sampling time constant, its fluctuations will directly modulate the sampling step size and may generate complex sampling behaviors.

### 4.1 The bump height dynamics and its equilibrium distribution

We next analyze the dynamics of the bump height *U*_*E*_ and then find its fluctuation statistics. Previous studies identified the change of bump height *U*_*E*_ as the second dominant eigenmode of the circuit dynamics (Appendix B, [38, 57]), and its dynamics is derived by projecting the circuit dynamics onto the height eigenvector (details in Appendix B.3),

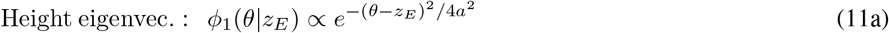

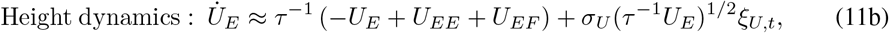

where 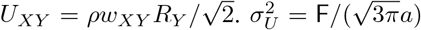 is a constant determining the strength of internal variability projected onto the height subspace, and doesn’t change with network activities.

### Dependency between position and height subspaces

Combining Eqs. (8b and 11b), it is worth noting that the *U*_*E*_ dynamics governs the step size of *z*_*E*_ dynamics (Fig. 2C), and hence the two dynamics are not independent, even if the eigenvectors of bump position and height are *locally* orthogonal. In the nonlinear attractor dynamics, both position and height eigenvectors depend on the instantaneous *z*_*E*_ (Eqs. 8a, 11a). The dependency over subspaces provides an important backbone supporting the scale-mixture structure in the sampling dynamics, as will be shown below.

### The equilibrium bump height distribution

We use the Fokker-Planck approach to determine the equilibrium *p*(*U*_*E*_) that can be directly formulated based on the Langevin equation of *U*_*E*_ (Eq. 11b):

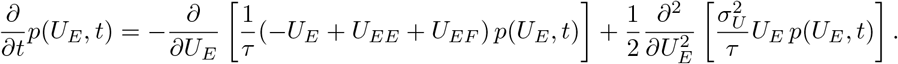

Setting *∂p/∂t* = 0, the equilibrium distribution can be analytically solved (Appendix C.1),

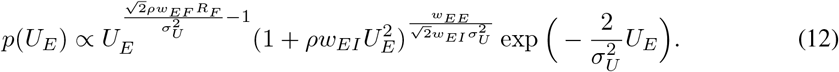

In the circuit model, the global inhibition strength *w*_*EI*_ is typically a small positive value (e.g., *w*_*EI*_ = 5 × 10^*−*4^) to enable the circuit produces physiologically realistic firing rates. This leads to the 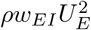 within the parenthesis way smaller than 1 in the main support of the distribution. Ignoring it allows us to obtain a convenient approximation of *p*(*U*_*E*_) that satisfies a Gamma distribution, i.e.,

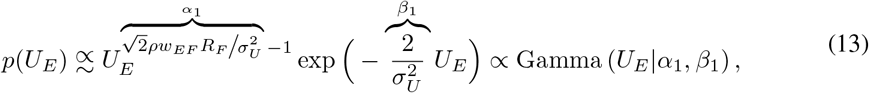

Moreover, to confirm *p*(*U*_*E*_) satisfies the Gamma distribution, we compare its goodness of fit to a Gaussian fit (Fig. A1B), which reveals that the Gamma distribution fits better than the Gaussian for most of the parameter regime especially the low-input regime. Lower input *R*_*F*_ leads to a smaller shape parameter *α*_1_, resulting in a highly skewed distribution. Conversely, at higher input regimes, the increased *α*_1_ causes the distribution to naturally converge toward a Gaussian-like bell curve.

### 4.2 Scale-mixture Langevin sampling in the position subspaces

#### Gaussian-conditional position step-size distribution

We next analyze how the *U*_*E*_ fluctuations affect the position *z*_*E*_ dynamics. **First**, the *U*_*E*_ fluctuations will leave the sampling distribution *p*(*z*_*E*_) **invariant** in the equilibrium, i.e., the equilibrium distribution of *z*_*E*_ will still match the posterior *π*(*z*). **Second**, the *U*_*E*_ fluctuations will significantly reshape the **sampling trajectories** of *z*_*E*_, especially its increment or step size, Δ*z*_*E*_(*t*) = *z*_*E*_(*t* + Δ*t*) − *z*_*E*_(*t*). To see this effect, we discretize the position *z*_*E*_ dynamics using the Euler-Maruyama method with a time step Δ*t*:

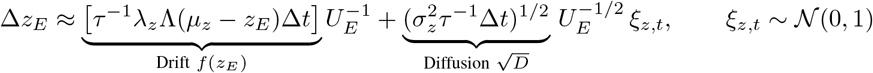

where we define the drift term *f* (*z*_*E*_) = *τ*^*−*1^*λ*_*z*_Λ(*µ*_*z*_ − *z*_*E*_)Δ*t* and the diffusion variance parameter 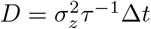. Then the distribution of Δ*z*_*E*_ conditioned on *U*_*E*_ is a Gaussian:

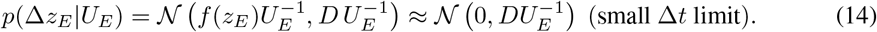

It clearly shows that Δ*z*_*E*_ depends on the bump height *U*_*E*_ in a **multiplicative** way (Fig. 2C), which is the key mechanism generating the scale-mixture structure in the sampling dynamics. In the limiting case when Δ*t* is sufficiently small, the drift term *f* (*z*_*E*_) scales linearly with Δ*t*, while the diffusion standard deviation 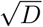 scales with 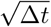. Thus, the drift term is of higher order than the diffusion term and becomes negligible. Eventually, the conditional distribution converges to a zero-mean Gaussian distribution with variance inversely scaling with *U*_*E*_ (Eq. 14). Fig. 2A confirms our theoretical derivation, showing that *p*(Δ*z*_*E*_) can largely be approximated as a zero-mean Gaussian distribution with variance inversely scaling with *U*_*E*_.

### Marginal position step-size Δ*z*_*E*_ distribution

The marginal step-size distribution is obtained by integrating over the Gamma height distribution *p*(*U*_*E*_) (Eq. 13). This integration reveals a **Gaussian scale mixture** structure, where the bump height *U*_*E*_ serves as the scale “mixer” variable [41, 42, 44]:

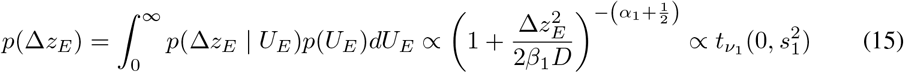

Note that this calculation implicitly utilizes the independent fluctuations of *U*_*E*_ and *z*_*E*_, i.e., *ξ*_*z,t*_ and *ξ*_*U,t*_ are independent. Although both sources of variability originate from the same internal noise in the neural dynamics (Eq. 1), they act independently because they are projected onto two locally orthogonal subspaces (the position and height subspaces, Eqs. 8a and 11a).

The resulting marginal *p*(Δ*z*_*E*_) is a Student-*t* distribution defined by two parameters: degrees of freedom (DoF) *v*_1_ = 2*α*_1_ and a scale parameter 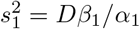 (details in Appendix C.3).

- The DoF *v*_1_ uniquely controls the sharpness of the central peak and the heaviness of the tail. When *v*_1_ is small (near 1), it produces a much sharper central peak than a Gaussian, combined with very thick, heavy tails where the probability density decays polynomially rather than exponentially. This regime accommodates robust sampling and extreme outliers.
- The scale parameter *s*_1_ controls the statistical dispersion, acting as a horizontal dilation or compression factor. Increasing *s*_1_ stretches the distribution outwards from the center and simultaneously lowers the peak density to ensure the total area under the curve remains exactly 1.

To empirically validate the scale-mixture model, we analyze how feedforward input intensity *R*_*F*_ shapes the sampling geometry (Fig. 2D). Under weak input intensity *R*_*F*_, the network operates in a superlinear regime where bump-height distribution (*p*(*U*_*E*_)) is more Gamma like. This generates highly sparse, heavy-tailed marginal step-size distributions *p*(Δ*z*_*E*_) that strongly favor a Student-*t* fit over a Gaussian baseline (Fig. 2F). Conversely, as *R*_*F*_ increases, the network exits this superlinear phase, and the step-size fluctuations become increasingly Gaussian-like (Fig. 2E).

Network simulations confirm the Student-*t* distribution of the position increments Δ*z*_*E*_ across a wide range of parameters (Fig. 3A). This is demonstrated by comparing the difference in the Akaike Information Criterion (ΔAIC) between the Student-*t* and Gaussian models for the step-size segments. The ΔAIC is consistently and heavily negative across all evaluated feedforward input intensities *R*_*F*_ and recurrent weights *w*_*EE*_, strongly penalizing the Gaussian assumption. This shows that the heavy-tailed Student-*t* distribution is a significantly better fit, particularly in weakly activated regimes where the disparity reaches its maximum magnitude.

### 4.3 Heavy-tailed dynamics accelerate sampling convergence

The heavy-tailed step-size Δ*z*_*E*_ distribution enables the circuit sampling to execute occasional large jumps (Fig. 2F), facilitating rapid exploration of the stimulus subspace. As shown in Fig. 2G, the circuit sampling converges much faster to the posteriors compared to a Langevin sampler whose the bump height *U*_*E*_ is fixed at the temporal average. This suggests the bump height fluctuations can benefit the Bayesian sampling, rather than merely noise that degrades the sampling performance.

Furthermore, systematic evaluation across the circuit parameter space reveals a distinct Lévy-like exploration phase (Fig. 3). This behavior is typically approximated using an *α*-stable distribution, where a stability parameter Θ between 0 and 2 defines the Lévy-like regime. To **robustly** quantify this heavy-tailed behavior, we estimate the CCDF of the step sizes, 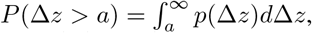, which is a parameter-free method, unlike the estimation of empirical probability density function (PDF) that needs a bin size. Importantly, a power-law tail in the CCDF Δ*z*^*−*Θ^ corresponds to a power-law probability density tail, *p*(Δ*z*) ∝ Δ*z*^*−*(Θ+1)^ (Fig. 2F) [60]. Therefore we will estimate the probability density tail through estimating the tail of CCDF.

As shown in Fig. 3B, this Lévy-like behavior emerges in the weak input limit (low feedforward input *R*_*F*_ or weak recurrent weight *w*_*EE*_). In this regime, the low baseline bump height allows multiplicative internal noise to induce massive transient fluctuations, propelling the network state across vast distances in the stimulus feature space.

## 5 Scale-mixture sampling in two-dimensional recurrent circuits

To demonstrate the generality of the identified scale-mixture sampling mechanism in recurrent circuits, we extend our analysis from the 1D recurrent circuit (the preferred stimulus *θ* is 1D, Eq. 1) to 2D recurrent circuits whose preferred stimulus ***θ*** = (*θ*_1_, *θ*_2_)^*⊤*^ ∈ (−*π, π*)^2^ is 2D. Similarly, we consider ***θ*** are uniformly distributed across the whole 2D stimulus feature space and we consider the periodic boundary of either *θ*_*i*_ to avoid boundary effect. The whole circuit dynamics is basically the same as the 1D case, as we only need to replace the 1D preferred stimulus *θ* and the external stimulus feature *z* by the 2D preferred stimulus ***θ*** and **z** = (*z*_1_, *z*_2_)^*⊤*^ respectively from Eqs. (1-4) (details in Appendix D). The precision and mean can be flexibly sampled by the network (Fig. 4B-C).

**Figure 4:**
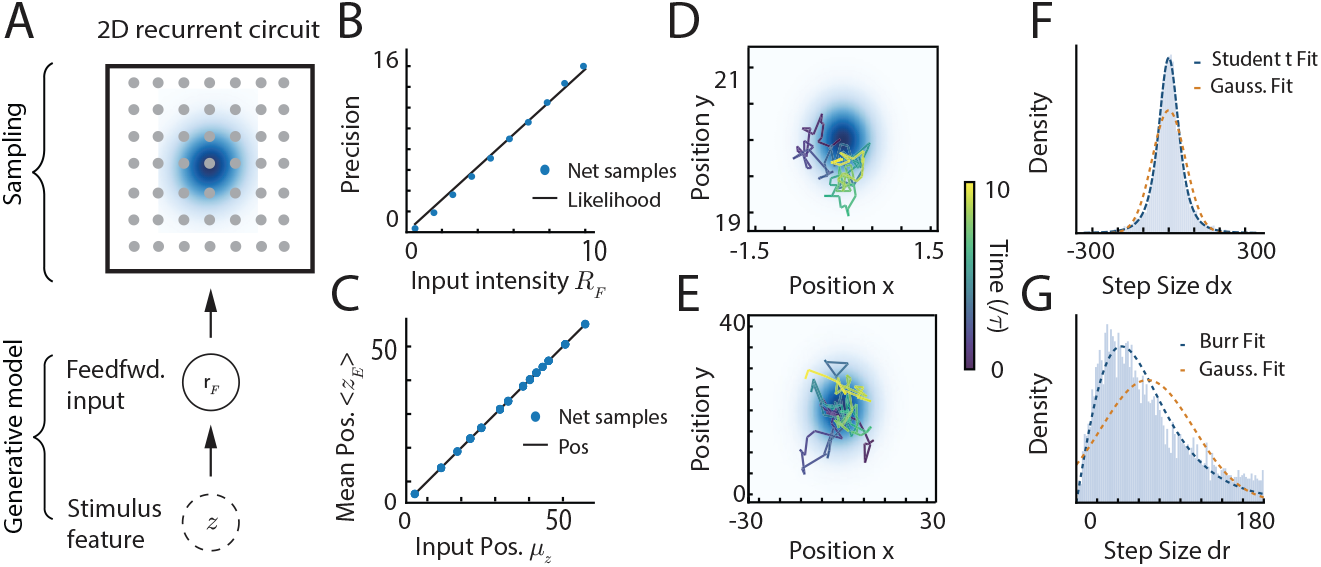
(A) The 2D recurrent circuit. (B) Precision of the network samples compared to the theoretical likelihood across different input intensities *R*_*F*_. (C) Mean position of the network samples compared to the actual input position. (D-E) The decoded 2D trajectories of sampled positions over time. The network can sample different posteriors with varying precision, showing a more concentrated trajectory in with *R*_*F*_ = 2.5 in (D) and a wider spread with *R*_*F*_ = 0.01 in (E). (F) Probability density of step size on each dimension (*dx*), overlaid with Student-*t* and Gaussian fits. (G) The distribution of radial step size (*dr*), overlaid with custom Burr fit (Eq. 19) and Gaussian fit.

### 5.1 Dimensionality reduction for the joint bump position and height dynamics

Following the same theoretical approach in the 1D case, the dynamics of the 2D bump position **z**_*E*_ and height *U*_*E*_ (still 1D) can be similarly derived (details in Appendix B.1),

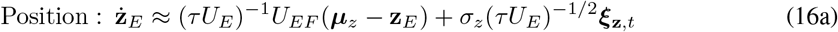

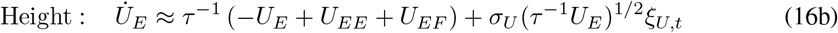

Several key parameters governing the amplitude dynamics remain one-dimensional (scalar) quantities, preserving the core structure of the 1D circuit: *U*_*E*_ represents the peak amplitude (height) of the 2D Gaussian bump. Regardless of the spatial dimensionality the bump spans, its maximum firing rate is a single scalar value. *R*_*F*_ is the maximum intensity (peak firing rate) of the external feedforward stimulus, which acts as a scalar scaling factor. *U*_*EF*_ represents the effective feedforward input drive. Because it is obtained by integrating the projection of the external input onto the bump’s spatial profile over the 2D space, the resulting magnitude is inherently 1D.

Similarly, the bump height *U*_*E*_ multiplicatively modulates the time constant of the position dynamics, while leaving the equilibrium distribution of **z**_*E*_ invariant. Performing similar analyses as the 1D case, the equilibrium distribution of the bump height can be approximated as a Gamma distribution:

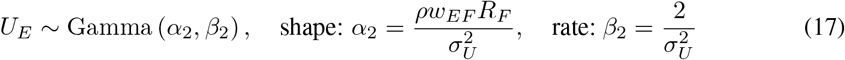

Empirical fits from network simulations confirm that *U*_*E*_ exhibits this expected long-tailed behavior rather than a symmetric Gaussian profile (Fig. A3A).

### 5.2 Gaussian scale mixture induces 2D heavy-tailed spatial exploration

The functional consequence of these bump height fluctuations is a direct modulation of the local 2D diffusivity. Over a short time interval Δ*t*, conditioned on a fixed bump height *U*_*E*_, the positional increment is an isotropic bivariate Gaussian (Appendix D). To determine the sampling step size statistics of the network, we integrate this conditional Gaussian over the Gamma distribution of *U*_*E*_, resulting in an isotropic bivariate Student-*t* distribution (Fig. 4F):

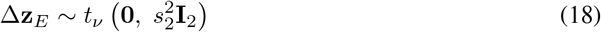

with degrees of freedom *v* = 2*α*_2_ and scale parameter 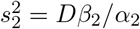.

A defining feature of the 2D sampling dynamics is the jump magnitude (radial displacement) *r* = ∥Δ**z**_*E*_∥ is a nonlinear function of the step size along each dimension. This transforms the bivariate Student-*t* distribution into another heavy-tailed distribution called the Burr distribution (Fig. 4G):

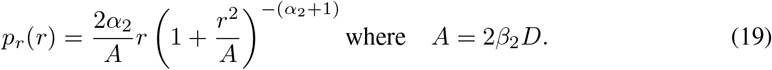

These marginal distributions demonstrate that the scale-mixture mechanism generalizes robustly to higher dimensions (Fig. 4F-G). Crucially, the heavy-tailed radial jump length *r* ensures that the network is not constrained to slow, purely diffusive random walks. Instead, it allows the 2D network to routinely execute large, multi-dimensional exploratory leaps, facilitating the rapid traversal of complex, high-dimensional joint posterior landscapes.

## 6 Conclusion and Discussion

The present study rigorously reveals the nonlinear CAN dynamics without explicit non-Gaussian components can collectively emerge the scale-mixture Langevin dynamics to sample the stimulus posterior in the circuit’s subspaces. The scale-mixture structure significantly speeds up the sampling convergence in the circuit, and it can enter the Lévy-like regime under extremely low circuit activation. Our study provides a minimal circuit mechanism for heavy-tail statistics generation, suggesting a deep structural alignment between the circuit’s intrinsic dynamics and the GSM used in describing natural statistics. To the best of our knowledge, this is the first theoretical work to analytically identify the Gaussian scale mixture structure in nonlinear recurrent circuit dynamics without non-Gaussian components.

### Comparison with previous studies

**First**, nearly all previous studies of long-tailed circuit Bayesian sampling explicitly incorporate non-Gaussian components in the circuit dynamics, e.g., the heavy-tailed synaptic weights [31], and/or the non-Gaussian variability [34, 35]. In contrast, our work emphasizes that the nonlinear recurrent circuit without explicit non-Gaussian components can internally generate the heavy-tailed sampling dynamics. Although a previous study also showed the heavy-tailed structure can emerge in recurrent networks without non-Gaussian components [61], it didn’t realize the posterior sampling that requires another constraint of the equilibrium distribution. And its circuit model is more complicated than ours by having neuronal adaptation. **Second**, although our circuit model is similar to previous studies of circuit sampling [16, 22], none of them explored the heavy-tailed sampling dynamics. **Third**, although heavy-tailed sampling algorithms were proposed in ML studies with top-down design [40], these algorithms have never been embedded into biologically plausible recurrent circuit dynamics. Our work provides a minimal, biological circuit mechanism for emerging heavy-tailed sampling.

### Limitations, generalizations, and future directions

**First**, <u>on the circuit model level</u>, the proposed circuit model doesn’t include diverse types of inhibitory neurons in canonical cortical circuits, including PV, SOM, and VIP neurons. And the inhibitory neuron pool that provides unstructured, divisive inhibition in our circuit model is regarded as the PV neurons [52, 54]. We believe adding the SOM neurons that provide structured, subtractive inhibition into the circuit model could enrich the dynamical repertoire of the circuit dynamics and algorithms, with a likely potential of enlarging the Lévy-like regime in the circuit parameter space. This was suggested by a previous study [61] whose neural adaptation is similar to the inhibition from SOM neurons. Moreover, the VIP neurons exclusively inhibit SOM neurons and can act as a “knob” to switch the circuit algorithms. **Second**, <u>on the types of posterior distributions</u>, as a proof-of-concept example, we only demonstrate the 1D and 2D Gaussian posterior sampling in our circuit model, and emphasize how the circuit internally generates heavy-tailed sampling dynamics to sample Gaussian posteriors. Based on the neural coding theory, the circuit can be easily extended to utilize scale-mixture Langevin sampling to sample other types of posteriors, as long as we change the spatial profile of recurrent kernel (Eq. 2) and feedforward input tuning (Eq. 4). Here, the spatial profile effectively acts as the natural parameter of the sampling distribution [6, 12]. **Finally**, our circuit model theory produces testable experimental predictions. We show the scale-mixture sampling will have heavier tails as the circuit activation decreases (Fig. 2E). This is generally supported by spatial trajectories of hippocampal place cells: the replay sequences decoded during sharp-wave ripples, during which the mean firing rate is lower, exhibit heavier tails of step size than the ones of theta sequences when the mean firing rate is higher [29, 30, 62].

## Supporting information

Supplementary Codes

## Acknowledgments and Disclosure of Funding

W.H.Z. is supported by the UT Southwestern Endowed Scholars program. The authors thank K.Y. Michael Wong and Ying Nian Wu on early formation of this project.

## Appendix

### A Appendix Figures

**Figure A1:**
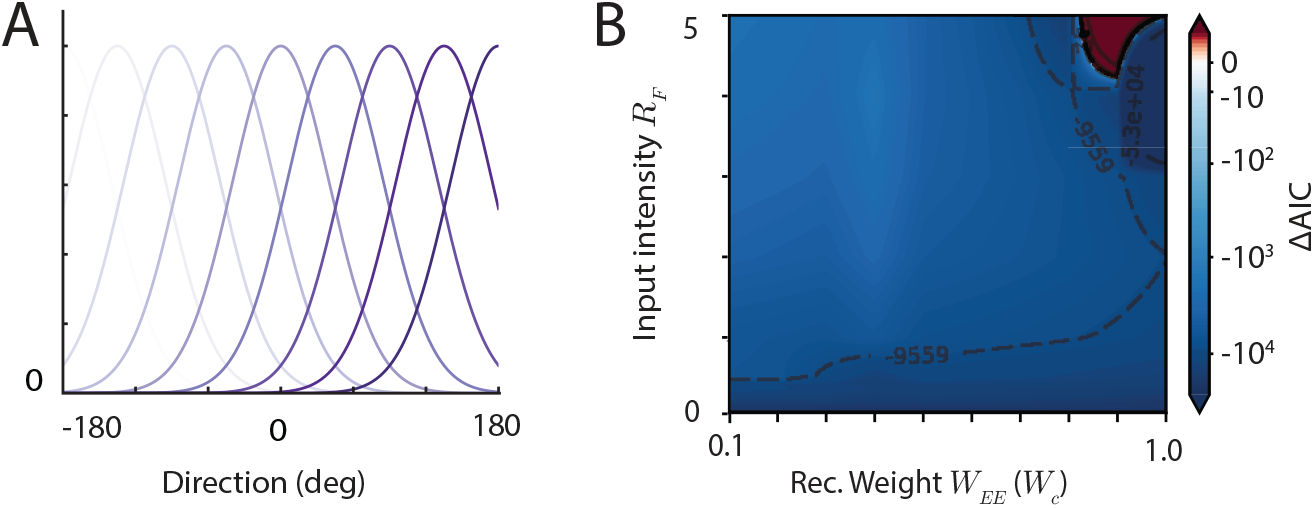
(A) Tuning curves of the neural population across the network space, mapping the preferred directions from− 180^*°*^ to −180^*°*^. (B) Heatmap of ΔAIC = AIC(Gamma) AIC(Gaussian) evaluating the goodness of fit for the bump height distribution across varying input intensities *R*_*F*_ and recurrent weights *W*_*EE*_.

**Figure A2:**
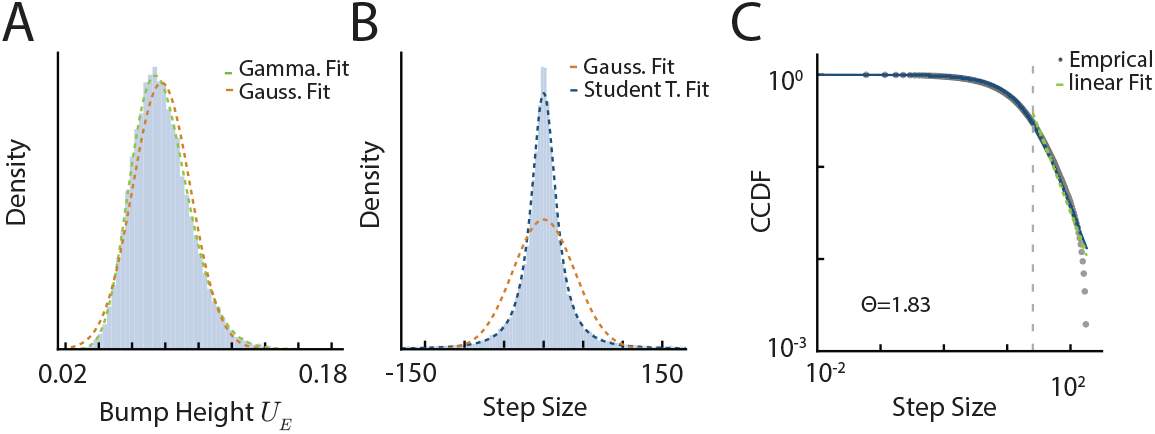
Sampling dynamics with a spatially shifted prior input. The prior input is provided at the same intensity as the likelihood input, but its position is shifted 5^*°*^ away from the likelihood. This configuration still successfully induces long-tail sampling behavior (see details in Sec. G.1). (A) Probability density of the Bump Height (*U*_*E*_), overlaid with Gamma and Gaussian fits. (B) Probability density of the step size, overlaid with Gaussian and Student-*t* fits. The Student-*t* fit better captures the heavy-tailed nature of the distribution. (C) Complementary Cumulative Distribution Function (CCDF) of the step size plotted on a log-log scale. The empirical data is overlaid with a linear fit in the tail region, revealing a tail exponent of Θ = 1.83.

**Figure A3:**
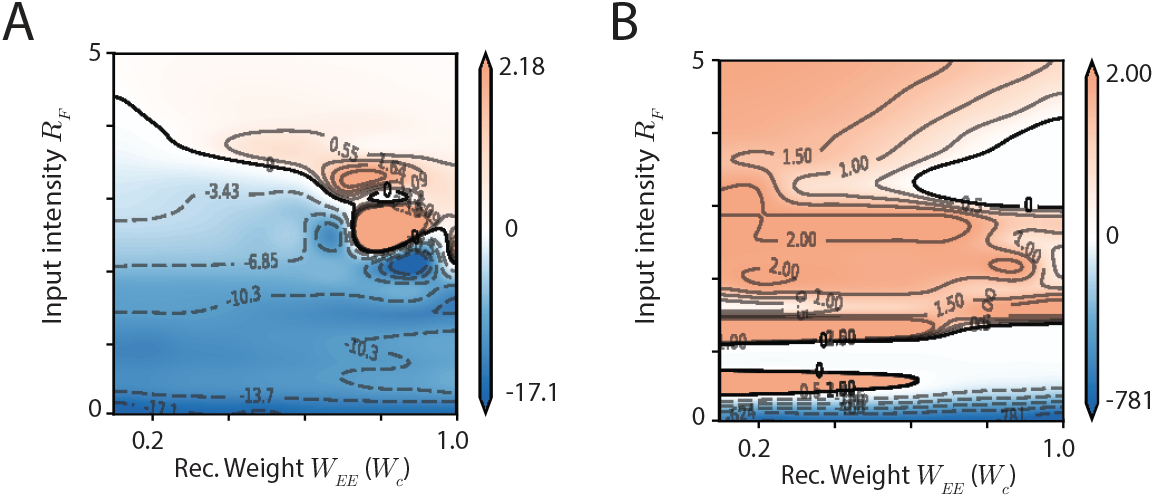
Goodness of fit analysis in the 2D network model. (A) Heatmap of ΔAIC = AIC(Gamma) −AIC(Gaussian) evaluating the goodness of fit for the bump height (*U*) distribution across varying input intensities *R*_*F*_ and recurrent weights *W*_*EE*_. (B) Heatmap of ΔAIC =AIC(Student-*t*) −AIC(Gaussian) evaluating the goodness of fit for the 1D step size (*dx*) distribution across varying input intensities *R*_*F*_ and recurrent weights *W*_*EE*_.

**Figure A4:**
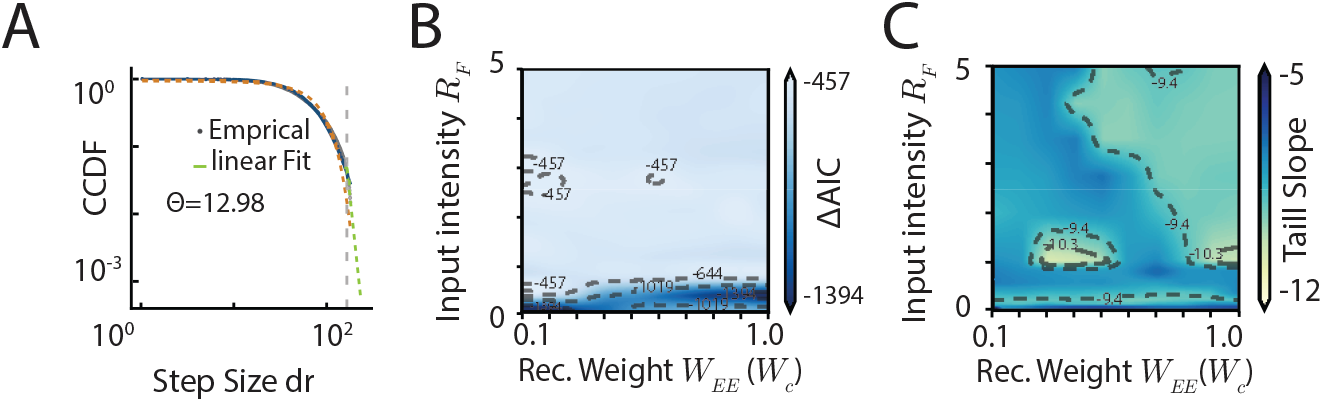
(A) Log-log plot of the Complementary Cumulative Distribution Function (CCDF) for the radial step size (*dr*), showing an empirical linear fit with a tasil exponent of Θ = 12.98. (B) Heatmap of ΔAIC comparing the goodness of fit for step size distributions across different input intensities *R*_*F*_ and recurrent weights *W*_*EE*_. (C) Tail slope of the step size distribution across different input intensities *R*_*F*_ and recurrent weights *W*_*EE*_.

### B A single canonical circuit and 1D natural gradient sampling: theory

In this section, we detail the mathematical foundation and theoretical properties of our simplified recurrent circuit model, which comprises excitatory (E) and inhibitory (I) neurons driven by continuous attractor network (CAN) dynamics.

#### B.1 Continuous attractor network dynamics

For clarity, we restate the governing equations of the network(Eq. 1). The temporal evolution of the excitatory population is described by:

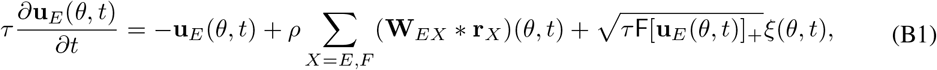

and the divisive normalization provided by I neurons (Eq. 3),

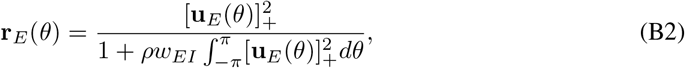

and the recurrent connection kernel **W**_*EX*_ (Eq. 2)

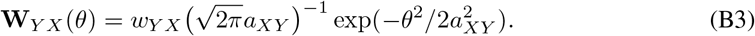

#### B.2 Network’s attractor states

We verify the proposed Gaussian ansatz of the attractor states of E neurons (Eq. 7),

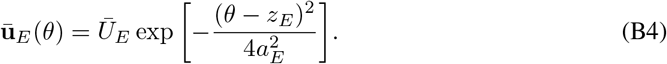

First, we substitute it into the divisive normalization (Eq. B2), yielding the following expression for the firing rate of E neurons,

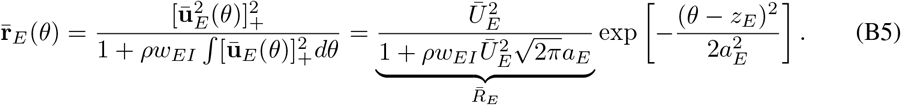

Then we use the above E firing rate (Eq. B5) to calculate the recurrent input from the neuronal population of type *Y* to the one with type *X* in the circuit model,

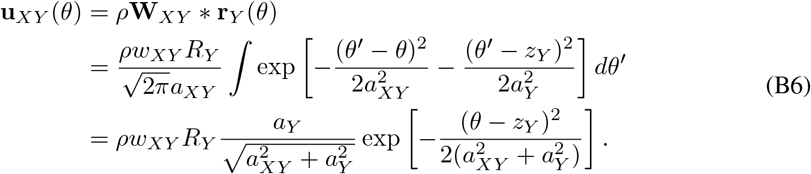

Specifically, based on Eq. (B6), the recurrent E population input is

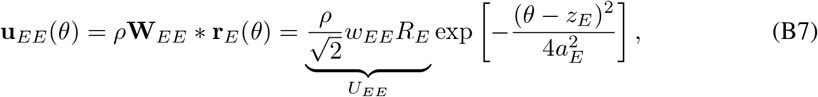

and the feedforward population input is

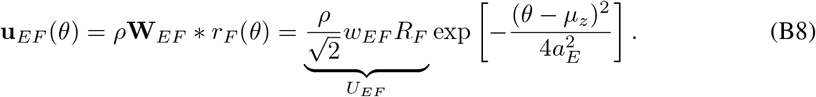

It can be checked the proposed Gaussian ansatz (Eq. B4) is indeed the sum of the recurrent input (Eq. B7) and the feedforward input (Eq. B8), i.e.,

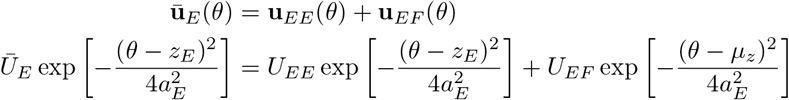

and implies

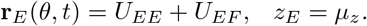

This completes the recurrent loop of the dynamics, and verify the validity of the Gaussian ansatz (Eq. 7).

#### B.3 Dimensionality reduction by projecting on dominant modes

We substitute Eqs. (B4-B8) into the Eq. (B1),

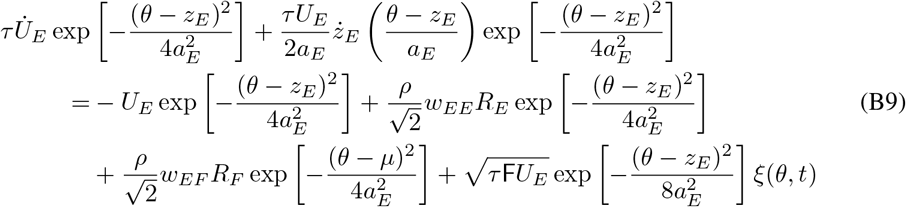

Previous studies analytically calculated the first two dominant eigenvectors (modes) [38, 57], corresponding to the change of the position and height of the Gaussian ansatz respectively,

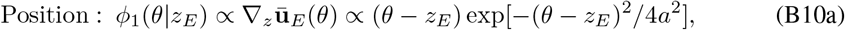

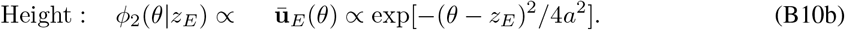

Projecting the dynamics Eq. (B9) into these 2 motion modes (Eq. B10), which means calculate the inner product ∫ *f* (*θ*)*ϕ*(*θ*|*z*_*E*_)*dθ* with *f* (*θ*) a term in Eq. (B9),

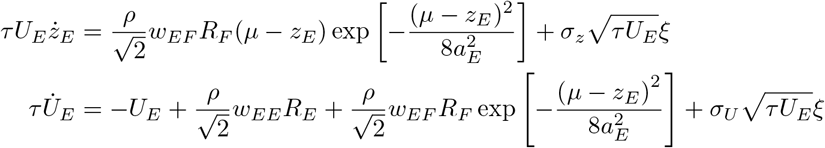

where

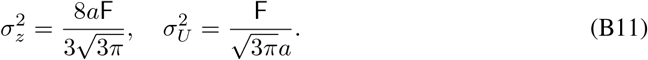

When the bump position *z*_*E*_ is near the input position, i.e., *µ* − *z*_*E*_ ≪ *a*_*E*_, which is usually the case in the circuit model, the exponential term exp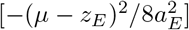 is close to one and can be safely ignored,

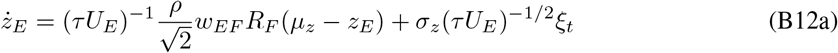

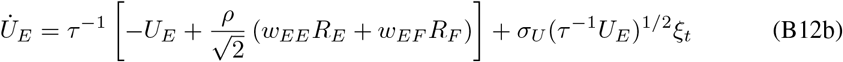

Furthermore, by using the notation

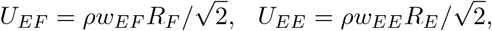

we arrive at Eqs. (8b and 11b) in the main text.

#### B.4 The probabilistic generative model embedded in the circuit model

##### B.4.1 The stimulus likelihood

We study how feedforward input defines the latent stimulus likelihood, i.e., ℒ (*z*) ∝ *p*(**r**_*F*_ |*z*). From the Eq. (4), the feedforward input **r**_*F*_ is modeled as a set of independent Poisson spike trains, where each neuron’s firing rate is Gaussian-tuned to the stimulus [6]:

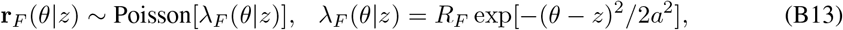

where *λ*_*F*_ (*θ*|*z*) is the mean firing rate of the neuron with stimulus preference *θ*. **r**_*F*_ denotes the peak input rate, and *a* specifies the tuning width. Explicitly writing the Poisson distribution of feedforward input spikes (we discretize the continuous *θ* into equally spaced *θ*_*j*_),

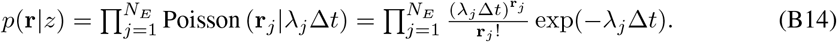

Taking the logarithm,

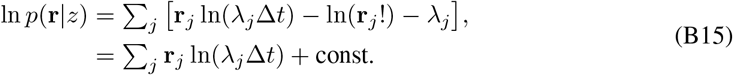

The const. in the above equation is under the assumption that the sum of population firing rate ∑_*j*_ *λ*_*j*_ is a constant irrelevant to latent stimulus *z*, which is true in a homogeneous population with a large number of neurons. Substituting the expression of the Gaussian tuning,

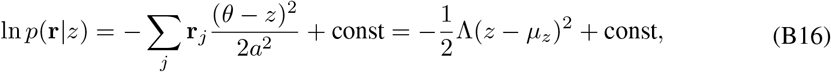

where

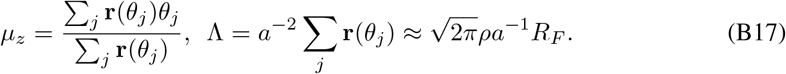

This implies the latent stimulus likelihood for the latent stimulus feature *z* given an observed feedforward input **r**_*F*_ is derived as a Gaussian distribution,

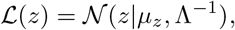

which is the Eq. (6) in the main text. Notably, the Gaussian distribution comes from the profile of the Gaussian tuning (Eq. B13) [6].

##### B.4.2 Uniform stimulus prior in the circuit

To convert the E bump position dynamics (Eq. B12a) into the Langevin sampling dynamics, it immediately suggests that the circuit stores a uniform (uninformative) stimulus prior, i.e., *p*(*z*) is uniform. This is because the gradient of the log-likelihood (∇ ℒ(*z*), Eq. 6) has the same form as the drift term in the E position dynamics

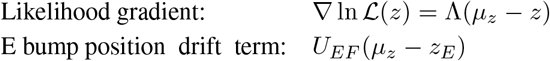

suggesting the gradient of the prior is zero, i.e., ∇ ln *p*(*z*) = 0. This uniform prior arises from the circuit’s homogeneous neurons (uniformly distributed in feature space) and its translation-invariant connection profile. Consequently, for the circuit to store a non-uniform prior, it must break this inherent symmetry in its neural organization and connectivity.

#### B.5 Conditions for realizing Langevin sampling in the circuit

The circuit sampling of the likelihood means the equilibrium distribution of the bump position (Eq. B12a) should match with the likelihood (Eq. 6). We copy the circuit bump position dynamics and the likelihood Langevin sampling dynamics in below for comparison,

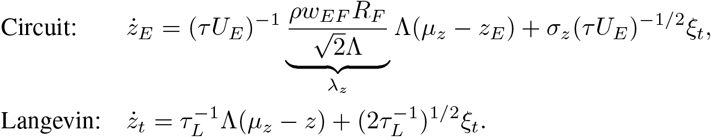

The *σ*_*z*_ is a constant that doesn’t change with neuronal activities. Therefore, the likelihood Langevin sampling in the circuit can be realized by setting the feedforward weight *w*_*EF*_ appropriately to make the ratio of the drift and diffusion coefficients the same as the Langevin sampling dynamics. The optimal feedforward weight can be found as (by using Eq. B17)

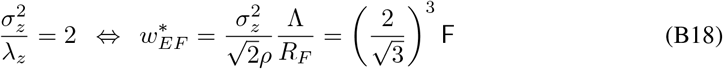

Furthermore, the time constant of the *z*_*E*_ dynamics is

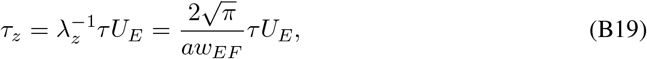

which is proportional to the E bump height *U*_*E*_. Finally, the equation of bump position (Eq. B12a) can be converted into the same form as a standard Langevin sampling,

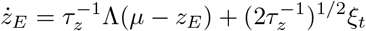

### C How to derive the distributions of bump height and position increments

#### C.1 Fokker-Planck equation for the bump-height dynamics

To derive the stationary distribution of the bump height, we start from the reduced stochastic dynamics of *U*_*E*_,

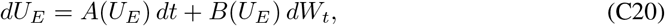

with drift and diffusion terms

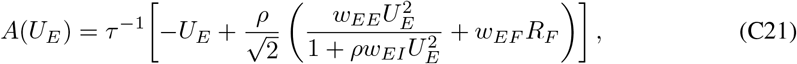

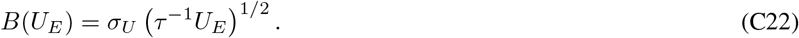

For a one-dimensional Langevin equation of the form *dX* = *a*(*X*) *dt* + *b*(*X*) *dW*_*t*_, the corresponding Fokker–Planck equation for the probability density *p*(*x, t*) is

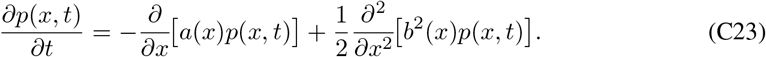

Applying Eq. (C23) to Eq. (C20) gives

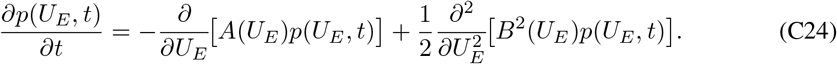

At steady state, *∂*_*t*_*p* = 0, so the stationary density *p*(*U*_*E*_) satisfies

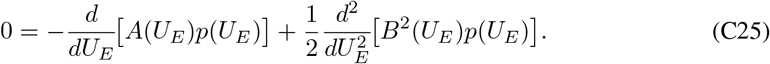

Assuming zero probability flux at the boundaries, we integrate once to obtain

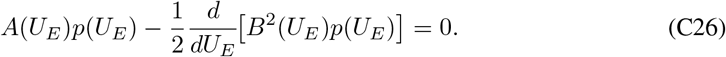

Rearranging gives

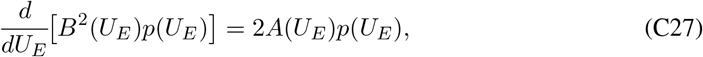

and therefore

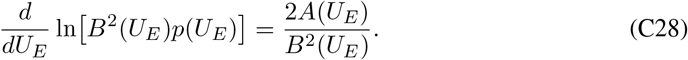

Integrating both sides yields the standard stationary solution

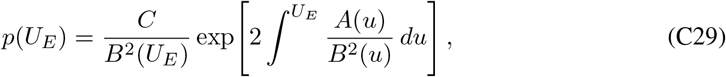

where *C* is a normalization constant.

For the present dynamics,

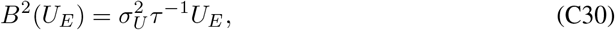

so that

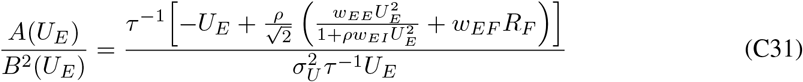

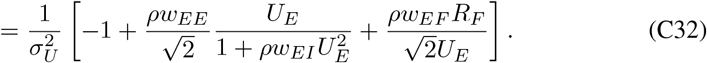

The integral in Eq. (C29) can then be evaluated term by term:

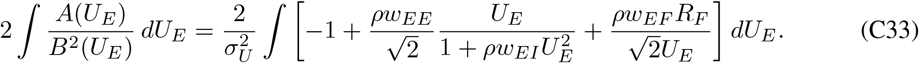

Using

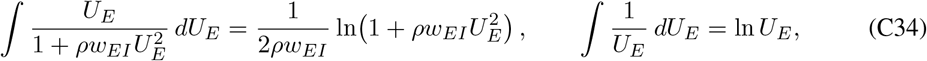

we obtain

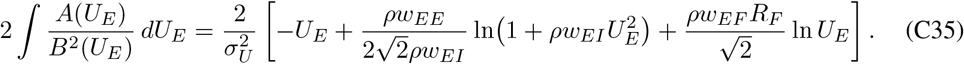

Substituting this result back into Eq. (C29) and absorbing all constants into the normalization factor gives

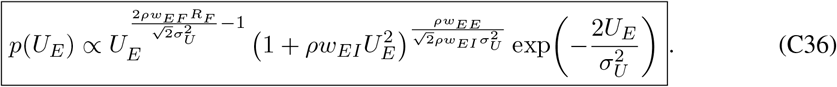

It is convenient to rewrite this stationary law as

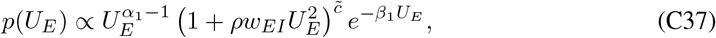

with

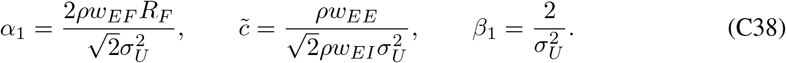

#### C.2 Approximation of the bump-height distribution

This distribution can be viewed as a Gamma-like kernel multiplied by an additional correction term that arises from recurrent excitation under divisive normalization.

To make this correction explicit, we rewrite

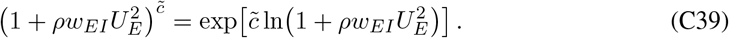

When 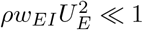, we use the Taylor expansion

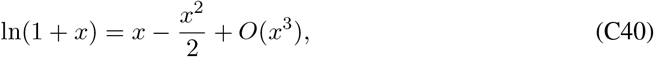

with 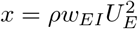, to obtain

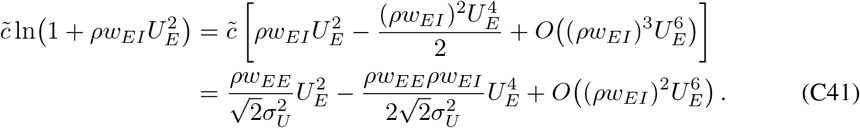

Therefore,

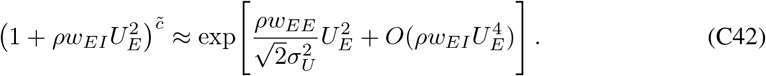

Substituting this into Eq. (C37) gives the local approximation

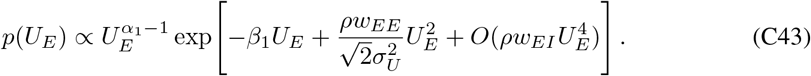

Equation (C43) shows that the exact stationary density is a Gamma-like distribution with a quadratic exponential correction. In the regime relevant for the leading-order analysis, we further assume that this correction is weak over the dominant support of the distribution, so that

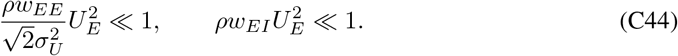

Then

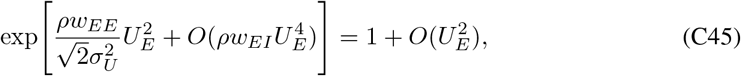

and to leading order we may neglect this factor entirely. The stationary law then reduces to

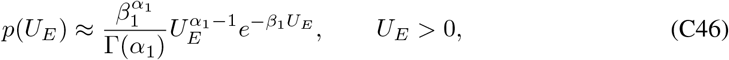

With

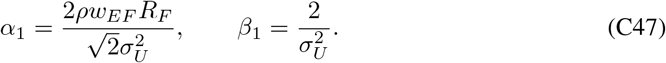

Thus, the Gamma approximation should be understood as the leading-order truncation of the more general stationary density in Eq. (C36). The retained exponential correction exp 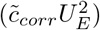, with 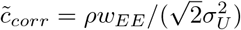, may be viewed as the first higher-order deviation from the Gamma form. Keeping this correction gives a refined approximation, whereas dropping it yields the simple Gamma law used in the main text and in the Gaussian scale mixture argument for the position increments.

#### C.3 Gaussian scaled mixture model

We begin with the reduced neural dynamics for the bump position:

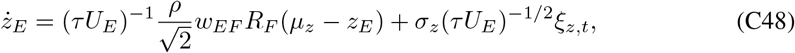

For a small time interval Δ*t*, the Euler–Maruyama approximation gives

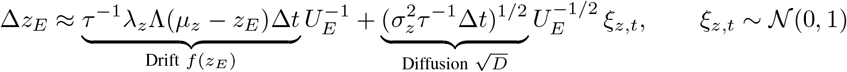

Hence, conditioned on *U*_*E*_, the displacement follows a Gaussian distribution:

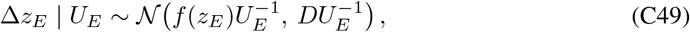

where 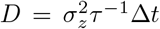. In the short-time or diffusion-dominated regime (as Δ*t*→ 0), the drift contribution *f* (*z*_*E*_) ∝ Δ*t* is subleading compared to the diffusion term 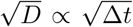. Thus, we approximate:

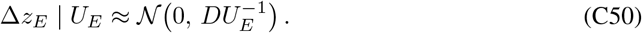

As the bump height is approximated as Gamma distributed:

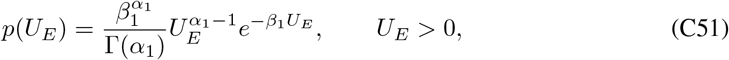

the marginal displacement density is obtained by integrating out *U*_*E*_:

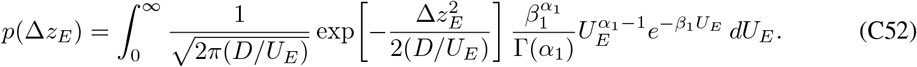

Using 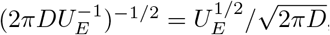, we obtain

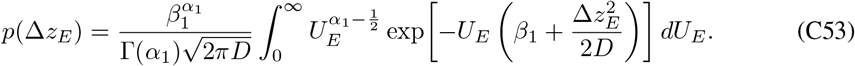

Rewriting the power as 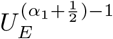 shows that this is a standard Gamma integral:

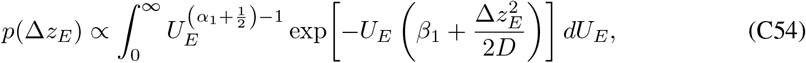

so that

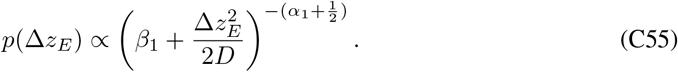

Factoring out *β*_1_ gives

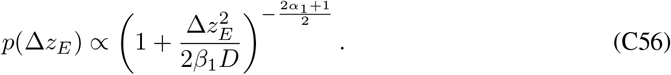

Comparing this with the standard Student-*t* distribution form:

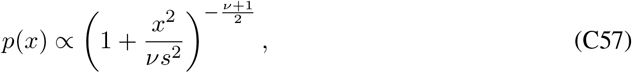

we can identify the degrees of freedom:

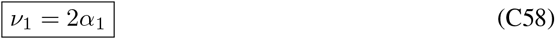

and the scale parameter:

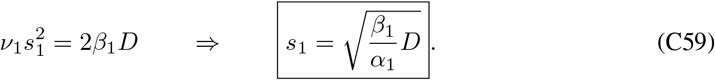

Therefore, the marginal step size follows a Student-*t* distribution:

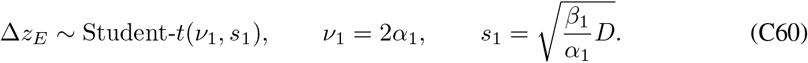

This rigorous derivation shows that the heavy tail arises directly from a Gaussian scale mixture: conditioned on a fixed bump height, the displacement is locally Gaussian, but integrating over the stochastic network fluctuations in *U*_*E*_ produces a heavy-tailed Student-*t* marginal law.

### D Dimensionality reduction in a 2D CANN

#### D.1 2D continuous attractor network dynamics

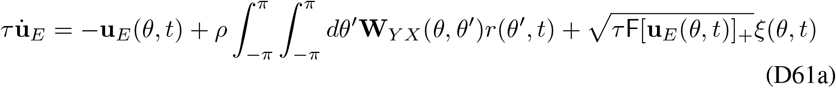

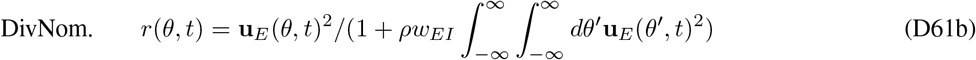

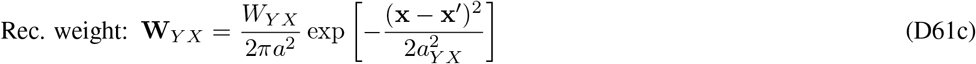

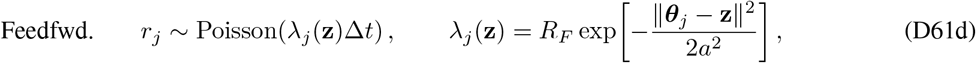

#### D.2 Network’s attractor state

To derive a low-dimensional description of the 2D network, we assume that the excitatory activity remains close to a Gaussian bump whose amplitude and center evolve slowly in time. We approximate the bump by

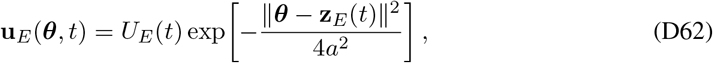

where **z**_*E*_(*t*) = (*z*_*E*,1_(*t*), *z*_*E*,2_(*t*)) denotes the bump center. Under divisive normalization, the corresponding firing-rate profile is

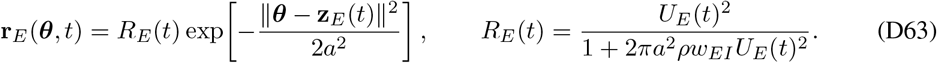

For a Gaussian kernel

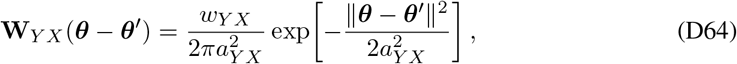

and a Gaussian firing profile centered at **z**_*X*_ with width *a*_*X*_, direct convolution yields

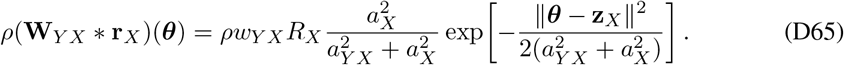

In the matched-width case *a*_*XY*_ = *a*_*X*_ = *a*, this reduces to

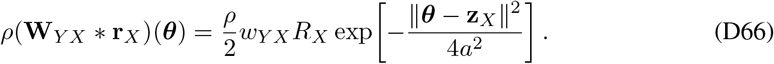

#### D.3 Dimensionality reduction by projecting on dominant modes

The dominant slow modes are the height mode

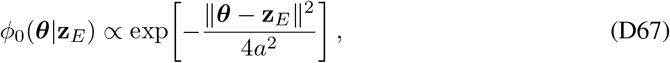

and the two translational modes

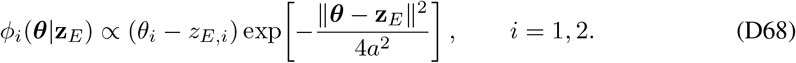

Thus the slow manifold is three-dimensional, parametrized by (*U*_*E*_, *z*_*E*,1_, *z*_*E*,2_).

Substituting Eq. (D62) into the network dynamics and projecting onto the height mode and the two translational modes yields the reduced stochastic dynamics

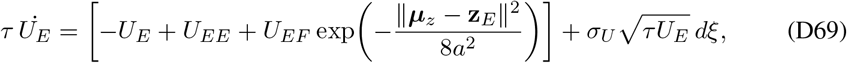

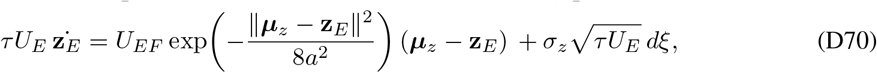

where *W*_*U*_ is a scalar Wiener process and **W**_*z*_ = (*W*_*z*,1_, *W*_*z*,2_)^*⊤*^ is a two-dimensional Wiener process with independent components. For the Gaussian bump ansatz, the projected noise strengths are

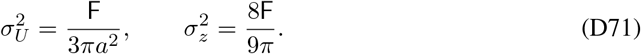

Here

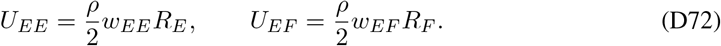

When ∥***µ***_*z*_ − **z**_*E*_∥ ≪ *a*, the equations simplify to

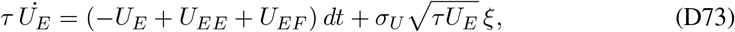

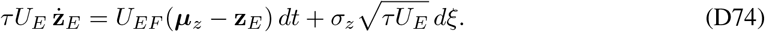

#### D.4 The probabilistic generative model embedded in the circuit model

##### D.4.1 The 2D stimulus likelihood with noise

We extend the feedforward population-code likelihood to a two-dimensional latent stimulus **z** = (*z*_1_, *z*_2_)^*⊤*^. Let each feedforward neuron have preferred stimulus ***θ***_*j*_ = (*θ*_*j*1_, *θ*_*j*2_)^*⊤*^, and let its spike count over a small time bin Δ*t* be conditionally independent with Gaussian tuning,

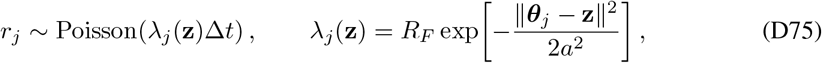

where ∥***θ***_*j*_ **z**∥ ^2^ = (*θ*_*j*1_ − *z*_1_)^2^ + (*θ*_*j*2_ − *z*_2_)^2^, *R*_*F*_ is the peak input rate, and *a* is the shared tuning width.

The likelihood of the population response 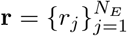 is

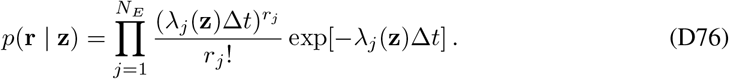

Taking the logarithm gives

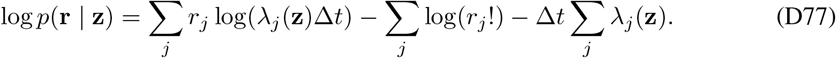

For a homogeneous dense population, ∑ _*j*_ *λ*_*j*_(**z**) is approximately independent of **z**, so the last two terms can be absorbed into the constant. Substituting the Gaussian tuning curve then yields

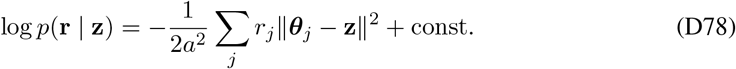

Expanding the quadratic form in **z**,

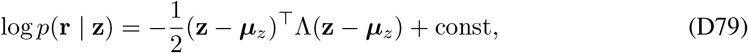

Where

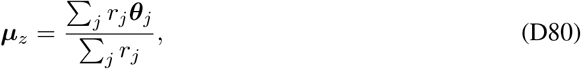

and

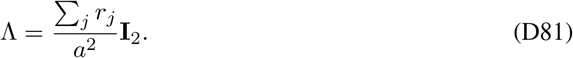

Therefore, conditioned on the observed noisy feedforward response, the latent stimulus likelihood is

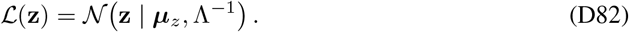

To make the noise explicit, let the true latent stimulus be **z**_*t*_, and define the total feedforward count

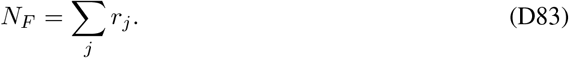

In the continuum limit for a homogeneous population of density *ρ*,

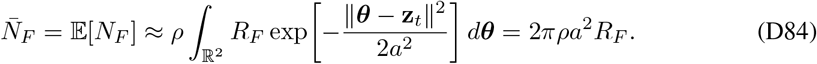

Allowing a Fano factor F for the feedforward spike-count variability,

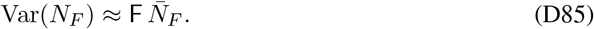

Hence the precision fluctuates around its mean as

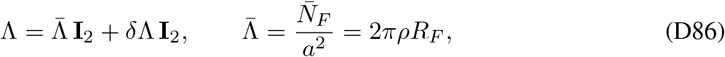

with

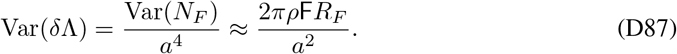

Similarly, a first-order expansion of the population vector gives

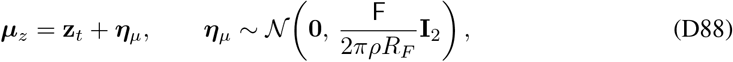

so that the decoded stimulus mean fluctuates isotropically around the true stimulus.

Thus, in the minimal 2D extension, the noisy feedforward population encodes an isotropic Gaussian likelihood whose center and precision both fluctuate from trial to trial:

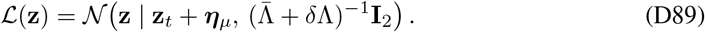

##### D.4.2 Uniform stimulus prior in the 2D circuit

For the minimal 2D extension, the feedforward population encodes an isotropic Gaussian likelihood over the latent stimulus **z** ∈ ℝ^2^,

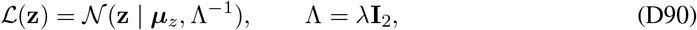

where ***µ***_*z*_ is the decoded 2D population vector and, in the continuum limit,

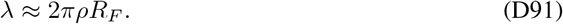

Therefore,

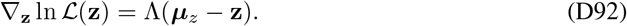

Comparing this likelihood gradient with the drift term in the reduced E-bump position dynamics shows that the circuit again stores a uniform (uninformative) prior over the 2D stimulus space, i.e., *p*(**z**) is constant. This is because the circuit is homogeneous and translation invariant on the 2D manifold, so there is no preferred location in stimulus space. Thus,

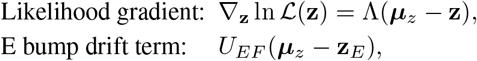

which implies

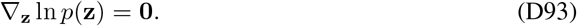

Hence, in the minimal 2D circuit, the posterior is determined entirely by the likelihood. As in the 1D case, a non-uniform prior would require breaking the translational symmetry of the network through non-homogeneous neural density, non-uniform external bias, or structured recurrent connectivity.

#### D.5 Conditions for realizing 2D Langevin sampling in the circuit

To realize likelihood sampling in the minimal 2D circuit, the equilibrium distribution of the bump position should match the likelihood in Eq. (D90). Near the stimulus, the linearized reduced bump-position dynamics can be written as

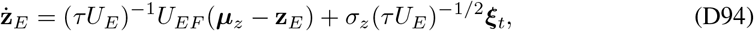

where ***ξ***_*t*_ is a two-dimensional white-noise process with independent components,

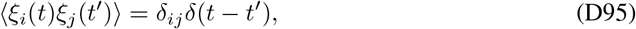

and

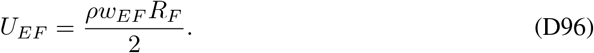

Using the 2D likelihood gradient,

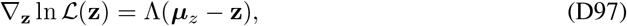

Eq. (D94) can be rewritten as

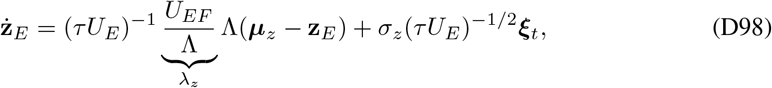

where, in the isotropic minimal extension,

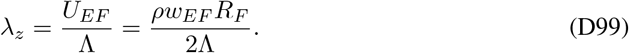

For comparison, the standard overdamped Langevin dynamics that samples the posterior *p*(**z** | **r**_*F*_) ∝ L(**z**)*p*(**z**) is

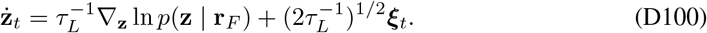

Because the prior is uniform, this reduces to

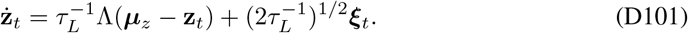

Therefore, the 2D circuit realizes Langevin sampling when its drift and diffusion coefficients satisfy the same ratio as in Eq. (D101). Matching the diffusion terms gives

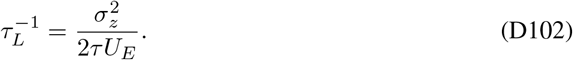

Matching the drift terms then requires

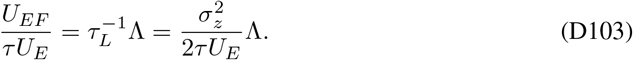

Equivalently,

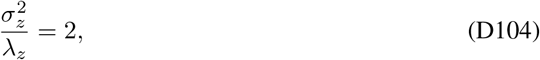

which yields the optimal feedforward weight

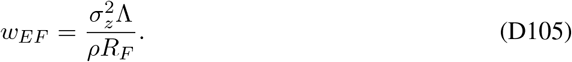

Using the isotropic 2D likelihood precision

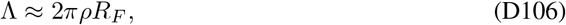

we obtain

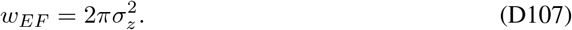

If we further substitute the projected 2D bump-position noise strength

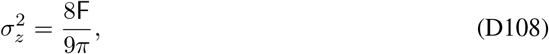

then the optimal feedforward weight becomes

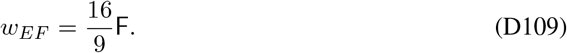

The corresponding sampling time constant is

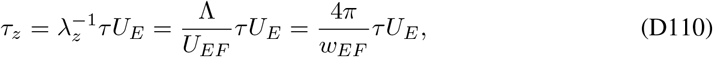

which is again proportional to the E bump height *U*_*E*_. Thus the bump-position dynamics can be written in the standard 2D Langevin form

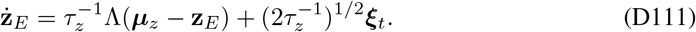

### E The distribution of step size in 2d case

#### E.1 2D Fokker–Planck equation for the bump-height dynamics

For the reduced 2D CANN, the bump height satisfies

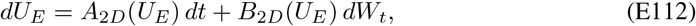

with

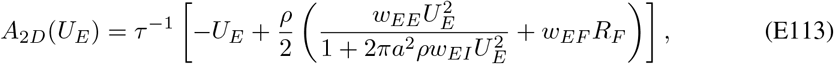

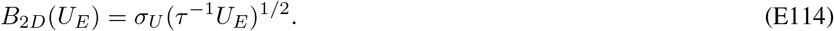

The corresponding Fokker–Planck equation is

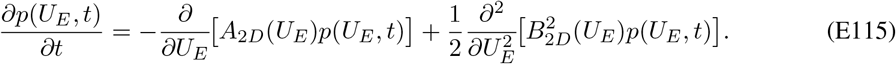

At stationarity, imposing zero probability flux gives

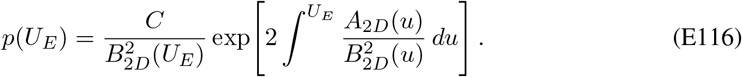

Since

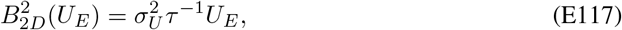

we obtain

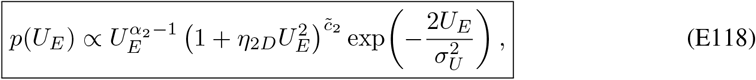

where

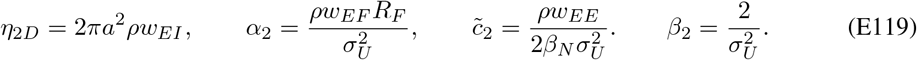

#### E.2 Gamma approximation of the 2D bump-height distribution

When 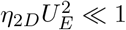, we write

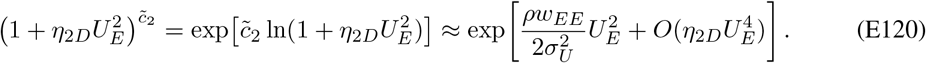

If this quadratic correction is weak over the dominant support, then to leading order

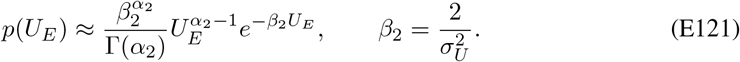

#### E.3 2D Gaussian scale mixture model for bump-position increments

The reduced bump-position dynamics in 2D is:

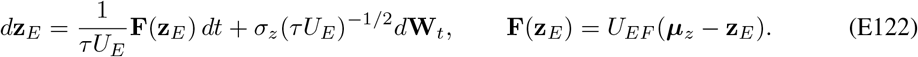

Over a short interval Δ*t*, the increment conditioned on *U*_*E*_ is:

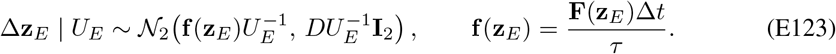

In the diffusion-dominated regime (small Δ*t*), we neglect the subleading drift term **f** (**z**_*E*_) and use:

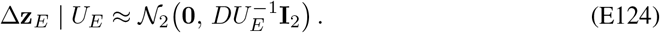

Mixing over the Gamma law of *U*_*E*_ yields:

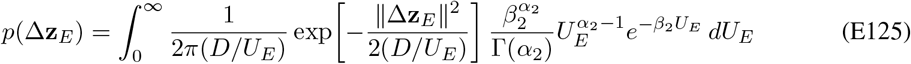

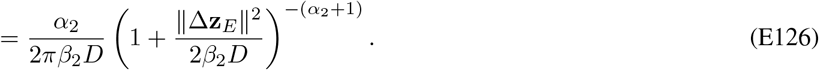

This is an isotropic bivariate Student-*t* distribution:

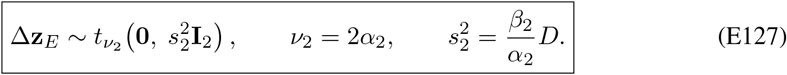

Each coordinate marginal therefore obeys:

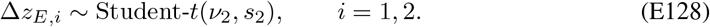

#### E.4 Distribution of the displacement magnitude

Define the jump length (radial step size):

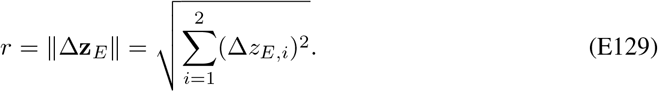

Under the short-time, diffusion-dominated approximation,

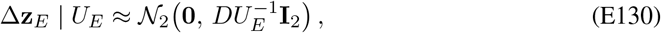

Equivalently,

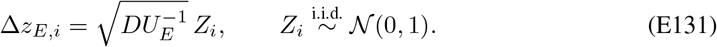

Hence

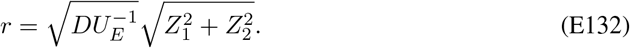

Since 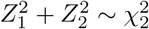, it follows that

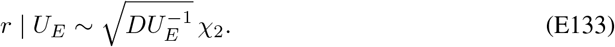

Equivalently, *r* | *U*_*E*_ follows a Rayleigh distribution with density:

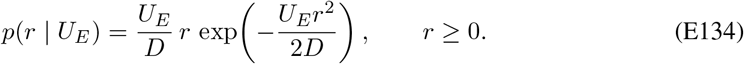

Using the Gamma approximation for the bump-height distribution,

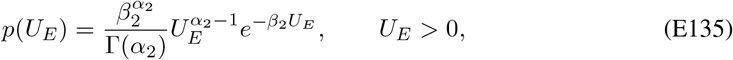

the marginal density becomes

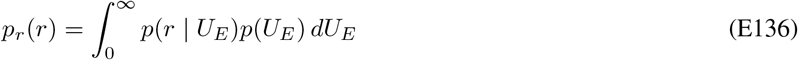

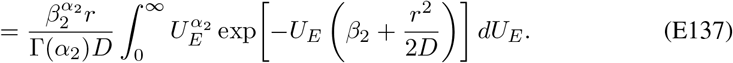

Applying the Gamma integral yields:

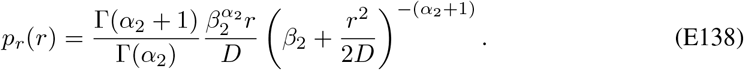

Using Γ(*α*_2_ + 1) = *α*_2_Γ(*α*_2_) and defining

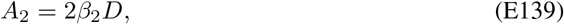

we obtain

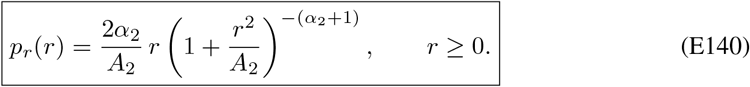

### F Circuit simulation parameters and details

#### F.1 Critical weight

To scale the connection strengths in our network model, we use a critical recurrent connection strength as a reference point. This critical strength is defined as the smallest value that allows the network to maintain persistent activity even when there is no feedforward input.

##### F.1.1 1D case

In the absence of feedforward input (*R*_*F*_ = 0), the stationary state of the 1D circuit’s excitatory bump height is determined solely by the recurrent input:

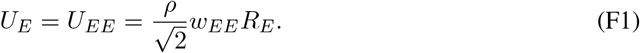

Furthermore, the firing rate of the E population, *R*_*E*_, is related to its input *U*_*E*_ by the divisive normalization activation function. Since the spatial integral of the squared 1D Gaussian bump is 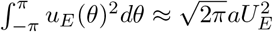, the firing rate is:

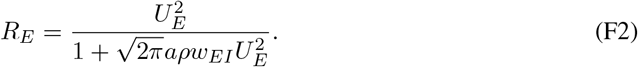

Substituting Eq. (F2) into Eq. (F1) allows us to write an equation solely in terms of *U*_*E*_:

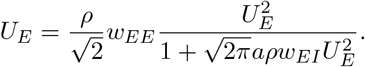

Assuming *U*_*E*_ ≠ 0 (for persistent activity), we can divide by *U*_*E*_ and rearrange the equation into a quadratic form for *U*_*E*_:

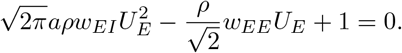

This quadratic equation for *U*_*E*_ has real solutions if and only if its discriminant is non-negative (Δ ≥ 0):

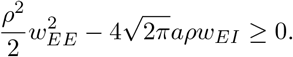

The critical weight, *w*_*c*_, is the smallest value of *w*_*EE*_ that permits non-zero persistent activity, which occurs when the discriminant is exactly zero:

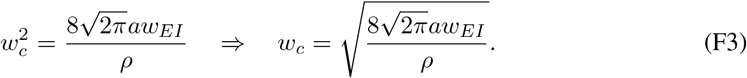

##### F.1.2 2D case

In the 2D network extension, the stationary state of the bump height without feedforward input is similarly driven by the 2D recurrent input projection:

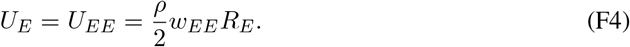

For the 2D divisive normalization, the spatial integral of the squared 2D Gaussian bump over the feature space is 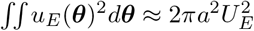. Thus, the peak firing rate becomes:

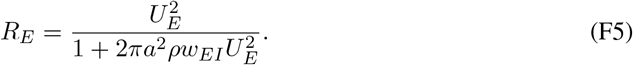

Substituting Eq. (F5) into Eq. (F4) yields:

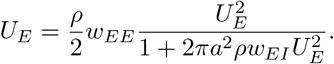

Assuming *U*_*E*_≠ 0, we divide by *U*_*E*_ and rearrange to form the 2D quadratic equation:

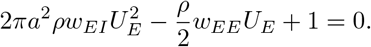

For persistent 2D activity to exist, the discriminant of this quadratic must be non-negative:

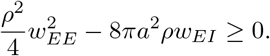

Setting the discriminant to zero gives the 2D critical weight *w*_*c*_:

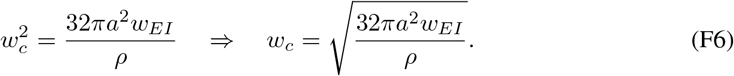

#### F.2 Simulation and Statistical Fitting Procedures

##### F.2.1 Network Simulation Protocol

To systematically investigate the network dynamics across different dynamical regimes, simulations were conducted over a comprehensive grid of parameters, specifically varying the feedforward input intensity (*R*_*f*_), the recurrent excitatory synaptic weight (*w*_*ee*_), and the global normalization constant (*k*).

For each parameter combination, the network dynamics were numerically integrated over a fixed simulation duration. To ensure the analysis accurately reflected the steady-state equilibrium properties of the network, an initial burn-in period (*t*_*steady*_) was discarded from the time-series data to eliminate transient initialization artifacts. From this raw steady-state time series, we extracted the macroscopic variables governing the network’s continuous attractor dynamics. The bump height (*U*) was calculated as the spatial integral of the excitatory synaptic input across the network, normalized by the theoretical bump width and neuron density. The center of mass (bump position, *z* for 1D, or **r** = (*x, y*) for 2D) was decoded at each time step, from which the raw coordinate step sizes between successive time points were extracted for statistical analysis.

##### F.2.2 Statistical Distribution Fitting

To characterize the stochastic nature of the network’s continuous attractor, we evaluated the empirical distributions of the macroscopic variables against several theoretical probability density functions. Parameters for all theoretical distributions were estimated using Maximum Likelihood Estimation (MLE). Model selection was performed by computing the Akaike Information Criterion (AIC) for each fit, where a lower (or negative relative) AIC indicates a superior model fit.

###### Bump Height (*U*)

The steady-state distribution of the integrated bump height *U* exhibits strictly positive properties. We compared fits between a symmetric Gaussian distribution and a Gamma distribution. The superiority of the Gamma distribution was determined via ΔAIC = AIC_Gamma_ − AIC_Gaussian_.

###### Coordinate Step Increments (Δ*z*, Δ*x*, Δ*y*)

For 1D spatial dynamics, the coordinate step size Δ*z* was extracted. For 2D spatial dynamics, the step vector was decomposed into its orthogonal components, Δ*x* and Δ*y*. To capture the anomalous diffusion and heavy-tailed jumping behaviors observed in specific parameter regimes, the empirical distributions of these signed increments were fitted to a Student’s t-distribution (characterized by degrees of freedom *v*, location *µ*, and scale *σ*) and compared against a standard Gaussian baseline.

###### 2D Step Magnitude (Δ*r*)

In the 2D network, the absolute step magnitude 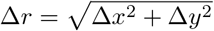 is strictly non-negative and requires a specialized radial distribution to accurately capture heavy-tailed radial jumps. We modeled this using a custom Burr distribution, defined by the probability density function:

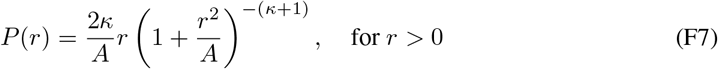

where *κ* dictates the shape (tail heaviness) and *A* governs the scale. The parameters were derived via numerical optimization of the negative log-likelihood. The AIC of this custom burr distribution was then compared against the AIC of a baseline Gaussian radial fit.

##### F.2.3 Heavy Tail Quantification

To systematically quantify the presence of heavy tails in the step-size distributions independently of global MLE fits, we analyzed the CCDF of the absolute step magnitudes.

The empirical CCDF, *P* (*X > x*), was computed and evaluated in log-log space. Rather than relying on a fixed heuristic threshold, the onset of the heavy tail was identified algorithmically. We iteratively performed linear regression on the log-log empirical CCDF across a sweeping range of candidate upper-percentile thresholds. The optimal tail threshold was selected as the point that maximized the Pearson correlation coefficient (*R*^2^), indicating the most linear tail regime in log-log space. The heavy-tail index Θ was subsequently extracted from the negative slope of this optimal linear fit.

#### A.4 Simulation Parameters

The core network simulations were carried out using the parameters listed in Table 1. The parameter sets for 1D and 2D spatial environments are largely identical, with specific scalar adjustments made to the global normalization constant, total simulation time, and the analytical feedforward scale to maintain consistent bump dynamics across dimensions.

**Table 1:** Core Network and Simulation Parameters

| Parameter | Variable | Value |
| --- | --- | --- |
| *Integration & Timing* |  |  |
| Time Constant | $\tau$ | 1 |
| Integration time step | $dt$ | $0.1 \tau$ |
| Total simulation time (1D) | $T$ | $500 \tau$ |
| Total simulation time (2D) | $T$ | $500 \tau$ |
| Burn-in period | $t_{steady}$ | $50 \tau$ |
| *Network Architecture* |  |  |
| Excitatory time constant | $\tau_E$ | 1.0 |
| Number of neurons | $N$ | 180 |
| Spatial domain | | $[-180^\circ, 180^\circ)$ |
| Excitatory connection width | $a_E$ | $40^\circ$ |
| *Dynamics & Synaptic Weights* |  |  |
| Fano factor (noise intensity) | $F$ | 0.5 |
| Baseline recurrent weight | $w_{ee}$ | $0.4w_c$ |
| Normalization constant (1D) | $k$ | $5 \times 10^{-4}$ |
| Normalization constant (2D) | $k$ | $5 \times 10^{-5}$ |
| Feedforward scale (1D) | $w^{EF}$ | 0.716 |
| Feedforward scale (2D) | $w^{EF}$ | 0.605 |

### G Generalization

#### G.1 Encoding a Non-Uniform Prior

To successfully encode a non-uniform marginal prior, the network must break the functional symmetry of its excitatory (E) neurons. Unlike standard models where tuning curves are homogeneous and translation-invariant across the stimulus space, storing a complex prior requires introducing heterogeneity. This can involve varying the tuning widths, amplitudes, and spatial distributions of the E neurons across the stimulus subspace.

Biologically, this required tuning heterogeneity can emerge through two primary circuit mechanisms:

- **Internal Network Structure:** Embedding the prior directly into the local circuit by superimposing a heterogeneous weight structure over the standard translation-invariant recurrent weight matrix.
- **External Top-Down Input:** Supplying a sustained prior signal from higher cortical regions.

Focusing on the external input mechanism, we can demonstrate that an applied prior signal successfully preserves the critical relationship between the bump amplitude and the Fisher Information (FI). For analytical simplicity, we model this external prior as a time-invariant Gaussian profile added to the excitatory network dynamics:

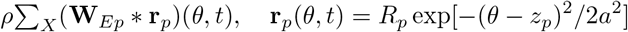

Incorporating this into the primary circuit dynamics yields:

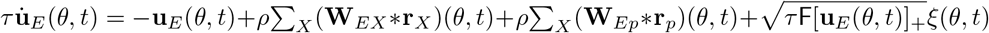

Because the external prior input functions mathematically in the same manner as the sensory feedforward input, the governing equations for the bump’s spatial position (*z*_*E*_) and amplitude (*U*_*E*_) can be updated directly. The resulting stochastic differential equations are:

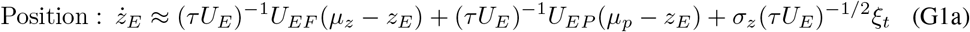

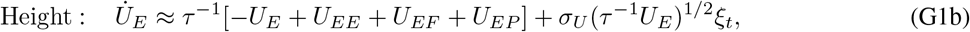

Here, *U*_*EP*_ and *µ*_*p*_ represent the precision and the mean of the non-uniform marginal prior, respectively. The drift term in the position dynamics now integrates both the sensory feedforward input and the external prior. As a result, when the bump position *z* reaches equilibrium, the network correctly samples from a posterior distribution that combines both sources of information:

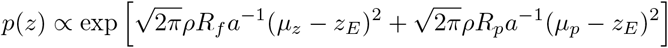

To validate this external prior configuration, we simulated the network dynamics with a spatially shifted prior. The prior input and the likelihood input were assigned equal intensities (*R*_*p*_ = *R*_*f*_ = 0.01). However, the prior’s mean (*µ*_*p*_ = 5) was deliberately shifted 5^*°*^ away from the likelihood’s mean (*µ*_*z*_ = 0).

As shown in the analysis of the sampling dynamics(Fig.A2), this offset configuration successfully preserves the network’s long-tail sampling behavior. The probability density of the bump amplitude (*U*_*E*_) exhibits right-skewed fluctuations that are best captured by a Gamma distribution, rather than a standard Gaussian fit. And the distribution of the sampling step sizes significantly deviates from simple diffusion. It tightly aligns with a Student-*t* distribution, confirming the presence of heavy-tailed jump dynamics. Plotting the CCDF of the step sizes on a log-log scale reveals a distinct linear relationship in the tail region. The empirical data yields a tail exponent of Θ = 1.83, mathematically demonstrating that the network utilizes long-range, Lévy-flight-like jumps to sample the modified posterior space efficiently.

