## Supplementary Codes for "Scale-mixture Langevin sampling in the subspaces of recurrent cortical circuit dynamics": 1d.py

```
import os
import glob
import tempfile
import gc
import traceback
import concurrent.futures
import multiprocessing as mp

import numpy as np
import pandas as pd
from scipy.stats import gamma, t, norm, linregress
from scipy.signal import savgol_filter
from scipy.interpolate import RectBivariateSpline, RegularGridInterpolator

import matplotlib.pyplot as plt
import matplotlib.colors as mcolors
import matplotlib.ticker as ticker
import seaborn as sns
from matplotlib.ticker import MaxNLocator
from matplotlib.colors import LinearSegmentedColormap, TwoSlopeNorm

from tqdm import tqdm

from cann_simulator import CANNSimulator 


def is_valid_npz(path, required_keys=None):
    required_keys = required_keys or ["time", "z_list", "Ue_list", "delta_z", "Rf", "k", "wee"]
    if not os.path.exists(path):
        return False
    if os.path.getsize(path) == 0:
        return False
    try:
        with np.load(path, allow_pickle=True) as data:
            keys = set(data.files)
        return all(k in keys for k in required_keys)
    except Exception:
        return False


def safe_save_npz(final_path, **arrays):
    folder = os.path.dirname(final_path) or "."
    fd, tmp_path = tempfile.mkstemp(dir=folder, suffix=".npz")
    os.close(fd)
    try:
        np.savez_compressed(tmp_path, **arrays)
        os.replace(tmp_path, final_path)
    except Exception:
        if os.path.exists(tmp_path):
            os.remove(tmp_path)
        raise


def prepare_integrated_data_1d(time, z_list, Ue_list, delta_z, params, win_bins=1):
    """Aligns, smooths, and cleans the 1D state series (No dx/dy)."""
    recording_start = params.get("recording_start", 0.0)
    L_state = min(len(time), len(z_list), len(Ue_list), len(delta_z))
    
    time_s = np.asarray(time[:L_state])
    U_s = np.asarray(Ue_list[:L_state])
    dz_s = np.asarray(delta_z[:L_state])

    U_dec = sliding_average(U_s, win_bins)
    dz_dec = sliding_average(dz_s, win_bins)
    time_dec = time_s[win_bins - 1:] 

    mask_dec = time_dec >= recording_start
    U_dec_clean = U_dec[mask_dec]
    U_dec_clean = U_dec_clean[np.isfinite(U_dec_clean) & (U_dec_clean > 0)]
    
    dz_dec_clean = dz_dec[mask_dec]
    dz_dec_clean = dz_dec_clean[np.isfinite(dz_dec_clean) & (np.abs(dz_dec_clean) > 1e-12)]

    return time_s, time_dec, U_dec_clean, dz_dec_clean


def rolling_mean_time(data, window, axis=0):
    """
    Sliding mean with stride=1 over time axis.
    - 1D: (T,) -> (T-window+1,)
    - 2D: (T,S) -> (T-window+1,S)
    """
    arr = np.asarray(data)
    if window <= 1:
        return arr

    if arr.ndim == 1:
        if arr.shape[0] < window:
            return np.empty((0,), dtype=float)
        c = np.cumsum(np.insert(arr.astype(float), 0, 0.0))
        return (c[window:] - c[:-window]) / window

    if axis != 0:
        arr = np.swapaxes(arr, 0, axis)

    T = arr.shape[0]
    if T < window:
        out = np.empty((0,) + arr.shape[1:], dtype=float)
        if axis != 0:
            out = np.swapaxes(out, 0, axis)
        return out

    c = np.cumsum(arr.astype(float), axis=0)
    c = np.concatenate([np.zeros((1,) + arr.shape[1:], dtype=float), c], axis=0)
    out = (c[window:] - c[:-window]) / window

    if axis != 0:
        out = np.swapaxes(out, 0, axis)
    return out


def sliding_average(arr, window):
    arr = np.asarray(arr)
    if len(arr) < window:
        return np.array([])
    return np.convolve(arr, np.ones(window)/window, mode='valid')


def fit_gamma_u(U):
    """Gamma fit (shape=kappa, scale=theta) on U>0."""
    U = np.asarray(U)
    U = U[np.isfinite(U) & (U > 0)]

    if U.size < 20:
        return np.nan, np.nan, np.nan

    try:
        shape_u, _, scale_u = gamma.fit(U, floc=0)
        loglik = np.sum(gamma.logpdf(U, a=shape_u, loc=0, scale=scale_u))
        aic = 2 * 2 - 2 * loglik
        return float(shape_u), float(scale_u), float(aic)
    except Exception:
        return np.nan, np.nan, np.nan


def generate_and_save_heatmaps_1d(time_array, Oc, U_field, params, Rf, wee, k, output_dir, win_bins=1):
    """Plots and saves the 1D spatio-temporal heatmaps."""
    if Oc is None or len(Oc) == 0:
        return
    
    def _ensure_time_first(mat, T):
        if mat is None:
            return None
        arr = np.asarray(mat)
        if arr.ndim != 2:
            return arr
        if arr.shape[0] != T and arr.shape[1] == T:
            return arr.T
        return arr
    
    Oc = _ensure_time_first(Oc, len(time_array))
    U_field = _ensure_time_first(U_field, len(time_array))
    time_s = np.asarray(time_array)[:len(Oc)]
    time_dec = time_s[win_bins - 1:]
    
    fig_hm = plt.figure(figsize=(18, 5), constrained_layout=True)
    gs_hm = fig_hm.add_gridspec(1, 3)
    ax_r = fig_hm.add_subplot(gs_hm[0, 0])
    ax_u = fig_hm.add_subplot(gs_hm[0, 1])
    ax_snap = fig_hm.add_subplot(gs_hm[0, 2])
    
    fig_hm.suptitle(f"1D Network Activity (Rf={Rf:.6f}, k={k:.7f}, wee={wee:.6f})", fontsize=16)

    def plot_map_rolling(ax, data, title, cmap):
        if data is None or len(data) == 0: return
        arr = np.asarray(data)[:len(time_s)]
        arr_roll = rolling_mean_time(arr, window=win_bins, axis=0) 
        if arr_roll.size == 0: return
        
        vmin, vmax = np.percentile(arr_roll, [1, 99])
        im = ax.imshow(arr_roll.T, aspect="auto", origin="lower",
                       extent=[time_dec[0], time_dec[-1], -180, 180],
                       vmin=vmin, vmax=vmax, cmap=cmap)
        ax.set_title(title)
        ax.set_xlabel("Time (s)")
        ax.set_ylabel("Position (deg)")
        plt.colorbar(im, ax=ax)

    plot_map_rolling(ax_r, Oc, "Firing Rate r(x,t)", "cividis")
    plot_map_rolling(ax_u, U_field, "Synaptic Input u(x,t)", "plasma")

    r_arr = np.asarray(Oc)[:len(time_s)]
    r_roll = rolling_mean_time(r_arr, window=win_bins, axis=0)
    if r_roll.size > 0:
        profile = r_roll[-1]
        x_deg = np.linspace(-180, 180, profile.shape[0])
        ax_snap.plot(x_deg, profile, linewidth=2, color='darkblue')
        ax_snap.set_title(f"Decoded profile (last window)")
        ax_snap.set_xlabel("Position (deg)")
        ax_snap.set_xlim(-180, 180)
        ax_snap.grid(True, alpha=0.3)

    os.makedirs(output_dir, exist_ok=True)
    fig_hm_path = os.path.join(output_dir, f"heatmaps_1d_Rf_{Rf:.6f}_k_{k:.7f}_wee_{wee:.6f}.svg")
    fig_hm.savefig(fig_hm_path, dpi=150, bbox_inches="tight")
    plt.close(fig_hm)


def fit_linear_tail(x, tail_q=0.95):
    """
    Isolates the tail of the data and fits a linear slope 
    in log-log space (Empirical CCDF).
    """
    x = np.asarray(x)
    x = np.abs(x) 
    x = x[np.isfinite(x) & (x > 1e-12) & (x<=180)]
    
    if x.size < 200:
        return dict(slope=np.nan, intercept=np.nan, u=np.nan, r_value=np.nan, tail_ok=False)

    # 1. Define the start of the tail
    u = float(np.quantile(x, tail_q))
    
    # 2. Extract and sort the tail data
    x_tail = np.sort(x[x > u])
    
    if x_tail.size < 10:
        return dict(slope=np.nan, intercept=np.nan, u=u, r_value=np.nan, tail_ok=False)

    # 3. Calculate the Empirical CCDF for the tail relative to the *entire* dataset
    N = len(x)
    ranks = np.searchsorted(np.sort(x), x_tail, side='right')
    ccdf_tail = (N - ranks + 1) / N

    # 4. Transform to log10-log10 space
    log_x = np.log10(x_tail)
    log_y = np.log10(ccdf_tail)

    # 5. Fit the linear slope
    try:
        slope, intercept, r_value, p_value, std_err = linregress(log_x, log_y)
        return dict(
            slope=slope,
            intercept=intercept, 
            r_value=r_value,
            u=u, 
            tail_ok=True
        )
    except:
        return dict(slope=np.nan, intercept=np.nan, u=u, r_value=np.nan, tail_ok=False)


def fit_t_and_evt(x, min_q=0.70, max_q=0.98, num_steps=50):
    """
    Global Student-t fit + automated Linear Tail fit for 1D magnitudes.
    Automatically finds the optimal tail threshold 'u' by finding the straightest 
    line in log-log space (maximizing R-squared).
    """
    x = np.asarray(x)
    x = np.abs(x)
    x = x[np.isfinite(x) & (x > 1e-12) &(x<=180)] 
    
    if x.size < 200:
        return dict(
            nu=np.nan, loc=np.nan, scale=np.nan, alpha_from_nu=np.nan,
            alpha_evt=np.nan, u=np.nan, tail_ok=False,
            linear_slope=np.nan, linear_intercept=np.nan, linear_r_squared=np.nan
        )

    # 1. Fit Student-t
    try:
        nu, loc, scale = t.fit(x)
        nu, loc, scale = float(nu), float(loc), float(scale)
    except:
        nu, loc, scale = np.nan, float(np.median(x)), np.nan

    alpha_from_nu = float(min(nu, 2.0)) if np.isfinite(nu) else np.nan

    # 2. Automated Linear Threshold Selection (Maximizing R^2)
    candidate_qs = np.linspace(min_q, max_q, num_steps)
    
    best_r_squared = -np.inf
    best_slope = np.nan
    best_intercept = np.nan
    best_u = np.nan
    best_r_val = np.nan
    
    for q_cand in candidate_qs:
        result = fit_linear_tail(x, tail_q=q_cand)
        
        if not result['tail_ok']:
            continue
            
        r_val = result['r_value']
        r_squared = r_val ** 2
        
        if r_squared > best_r_squared:
            best_r_squared = r_squared
            best_slope = result['slope']
            best_intercept = result['intercept']
            best_u = result['u']
            best_r_val = r_val

    # 3. Fallback if no valid candidates were found
    if np.isinf(best_r_squared) or np.isnan(best_r_squared):
        fallback_result = fit_linear_tail(x, tail_q=max_q)
        if not fallback_result['tail_ok']:
            return dict(
                nu=nu, loc=loc, scale=scale, alpha_from_nu=alpha_from_nu,
                alpha_evt=np.nan, u=np.nan, tail_ok=False,
                linear_slope=np.nan, linear_intercept=np.nan, linear_r_squared=np.nan
            )
        else:
            best_slope = fallback_result['slope']
            best_intercept = fallback_result['intercept']
            best_u = fallback_result['u']
            best_r_val = fallback_result['r_value']

    # 4. Calculate alpha (tail index) from the linear slope. 
    # For a CCDF, alpha is typically the negative of the slope.
    alpha_evt = float(min(-best_slope, 2.0)) if (np.isfinite(best_slope) and best_slope < 0) else 2.0
    # Get aic for the t fit
    aic = 2 * 3 - 2 * t.logpdf(x, df=nu, loc=loc, scale=scale).sum()  # 3 parameters: nu, loc, scale
    return dict(
        nu=nu, loc=loc, scale=scale, alpha_from_nu=alpha_from_nu,
        alpha_evt=alpha_evt, u=best_u, tail_ok=True,
        linear_slope=best_slope, linear_intercept=best_intercept, 
        linear_r_squared=(best_r_val**2), aic=aic
    )


def fit_gaussian(data):
    """Fits a Gaussian to the data and returns mean and std."""
    data = np.asarray(data)
    data = data[np.isfinite(data)]
    if data.size < 20:
        return np.nan, np.nan
    mean = float(np.mean(data))
    std = float(np.std(data))
    aic = 2 * 2 - 2 * np.sum(-0.5 * np.log(2 * np.pi * std**2) - 0.5 * ((data - mean) ** 2) / (std**2))
    return mean, std, aic


def _grid_key_1d(Rf, other, rf_decimals=3, other_decimals=7):
    return (round(float(Rf), rf_decimals), round(float(other), other_decimals))


def perform_integrated_analysis_1d(
    time, z_list, Ue_list, delta_z, params, Rf, wee, k, output_dir,
    save_diagnostic_plot=True, plot_range=180
):
    """Pure offline statistical analyzer for 1D step sizes and bump heights."""
    win_bins = 1
    time_s, time_dec, U, dz = prepare_integrated_data_1d(time, z_list, Ue_list, delta_z, params, win_bins)

    kappa_fit, theta_fit, aic_U = fit_gamma_u(U)

    dz_signed = dz[np.isfinite(dz) & (np.abs(dz) > 1e-12)]
    if dz_signed.size > 10:
        nu_signed, loc_signed, scale_signed = t.fit(dz_signed)
    else:
        nu_signed, loc_signed, scale_signed = np.nan, np.nan, np.nan

    dz_mag = np.abs(dz_signed - (loc_signed if np.isfinite(loc_signed) else 0))
    t_evt = fit_t_and_evt(dz_mag)

    if t_evt:
        t_evt["nu"] = float(nu_signed)
        t_evt["loc"] = float(loc_signed)
        t_evt["scale"] = float(scale_signed)

    # Wrapped Gaussian fit for z_list
    L = params.get("position_range", 360.0)
    low_z = -L / 2.0
    high_z = L / 2.0
    
    mean_z, var_z, std_z = fit_wrapped_gaussian_1d(z_list, low=low_z, high=high_z)
    
    aic_z = np.nan
    if np.isfinite(mean_z) and np.isfinite(var_z) and var_z > 0:
        pdf_vals = wrapped_gaussian_pdf_1d(z_list, mean_z, var_z, low=low_z, high=high_z)
        pdf_vals_clean = pdf_vals[pdf_vals > 1e-12]
        if len(pdf_vals_clean) > 0:
            nll_z = -np.sum(np.log(pdf_vals_clean))
            aic_z = 2 * 2 + 2 * nll_z 

    # Extract Gaussian parameters
    mean_dz, std_dz, aic_dz = fit_gaussian(dz)
    mean_U, std_U, aic_U_gauss = fit_gaussian(U)

    fig_dist_path = ''

    if save_diagnostic_plot:
        fig = plt.figure(figsize=(18, 5), constrained_layout=True)
        gs = fig.add_gridspec(1, 4)
        ax_dist_u = fig.add_subplot(gs[0, 0])
        ax_dist_dz = fig.add_subplot(gs[0, 1])
        ax_loglog = fig.add_subplot(gs[0, 2])
        ax_dist_z = fig.add_subplot(gs[0, 3])

        fig.suptitle(f"1D Distributions (Rf={Rf:.6f}, k={k:.7f}, wee={wee:.6f})", fontsize=18)

        # 1. Bump Height (U) Plot
        sns.histplot(U, bins=50, kde=False, stat="density", ax=ax_dist_u, color="lightsteelblue", edgecolor="None", linewidth=0)
        if U.size > 0:
            x_u = np.linspace(np.min(U)*0.9, np.max(U) * 1.1, 300)
            
            # Plot Gamma
            if np.isfinite(kappa_fit) and np.isfinite(theta_fit):
                ax_dist_u.plot(x_u, gamma.pdf(x_u, kappa_fit, 0, theta_fit), 
                               color="#1f4e79", ls='--', lw=3, label=f"Gamma fit (κ={kappa_fit:.2f}, θ={theta_fit:.4f})")
            
            # Plot Gaussian overlay
            if np.isfinite(mean_U) and np.isfinite(std_U) and std_U > 0:
                ax_dist_u.plot(x_u, norm.pdf(x_u, mean_U, std_U),
                               color="#d17c00", ls='--', lw=3, label=f"Gauss fit (μ={mean_U:.2f}, σ={std_U:.2f})")
            ax_dist_u.legend()
        ax_dist_u.set_title("Smoothed Bump Height U")

        # 2. Step Size (dz) Plot
        sns.histplot(dz, bins=100, binrange=(max(np.min(dz),-plot_range), min(np.max(dz),plot_range)), kde=False, stat="density", ax=ax_dist_dz, color="lightsteelblue", edgecolor="None", linewidth=0)
        if dz.size > 0:
            # Bound the theoretical curves to match the x-axis limits
            x_dz = np.linspace(max(np.min(dz),-plot_range), min(np.max(dz),plot_range), 1000)
            
            # Plot Student-t
            if t_evt and np.isfinite(t_evt.get("nu")):
                ax_dist_dz.plot(x_dz, t.pdf(x_dz, t_evt["nu"], t_evt["loc"], t_evt["scale"]),
                                color="#1f4e79", ls='--', lw=3, label=f"t fit (ν={t_evt['nu']:.2f}, loc={t_evt['loc']:.2f})")
            
            # Plot Gaussian overlay
            if np.isfinite(mean_dz) and np.isfinite(std_dz) and std_dz > 0:
                ax_dist_dz.plot(x_dz, norm.pdf(x_dz, mean_dz, std_dz),
                                color="#d17c00", ls='--', lw=3,  label=f"Gauss fit (μ={mean_dz:.2f}, σ={std_dz:.2f})")
            ax_dist_dz.legend()
            
        ax_dist_dz.set_title("Smoothed Step Size Δz")
        ax_dist_dz.set_xlim(-plot_range, plot_range)

        # 3. Log-Log Tail Diagnostics
        if dz.size > 0 and t_evt and np.isfinite(t_evt.get("loc")):
            loc_val = t_evt["loc"]
            rdata = np.abs(dz - loc_val)
            rdata = rdata[np.isfinite(rdata) & (rdata > 0)]

            if rdata.size > 5:
                r_sorted = np.sort(rdata)
                N = len(r_sorted)
                ccdf_emp = (N - np.arange(1, N + 1) + 1) / N
                plot_mask = r_sorted < 180
                r_sorted = r_sorted[plot_mask]
                ccdf_emp = ccdf_emp[plot_mask]

                # Sparsify arrays to prevent large file sizes
                max_points = 400
                if len(r_sorted) > max_points:
                    step = len(r_sorted) // max_points
                    r_plot = r_sorted[::step]
                    ccdf_plot = ccdf_emp[::step]
                    
                    if r_plot[-1] != r_sorted[-1]:
                        r_plot = np.append(r_plot, r_sorted[-1])
                        ccdf_plot = np.append(ccdf_plot, ccdf_emp[-1])
                else:
                    r_plot = r_sorted
                    ccdf_plot = ccdf_emp

                ax_loglog.loglog(r_plot, ccdf_plot, "o", ms=4, mfc="white", mec="0.45", mew=1.0, alpha=0.65, label="Empirical CCDF")

                r_grid = np.logspace(np.log10(r_sorted.min()), np.log10(r_sorted.max()), 400)
                ccdf_t = 2.0 * (1.0 - t.cdf(r_grid, df=t_evt["nu"], loc=0, scale=t_evt["scale"]))
                ax_loglog.loglog(r_grid, ccdf_t, "-", color="#1f4e79", lw=2, label=f"t CCDF (ν={t_evt['nu']:.2f})")

                if t_evt.get("tail_ok"):
                    u_val = t_evt["u"]
                    slope = t_evt["linear_slope"]
                    intercept = t_evt["linear_intercept"]
                    r2 = t_evt.get("linear_r_squared", np.nan)

                    if np.isfinite(slope) and np.isfinite(intercept):
                        r_tail = np.logspace(np.log10(max(u_val, np.min(rdata))), np.log10(np.max(rdata)), 300)
                        ccdf_tail = (10 ** intercept) * (r_tail ** slope)
                        ax_loglog.plot(r_tail, ccdf_tail, "--", color="#d17c00", lw=2, label=f"Linear Tail (α={-slope:.2f}, R²={r2:.2f}, u={u_val:.2f})")
                        ax_loglog.axvline(u_val, color="0.65", lw=1.6, ls=":", alpha=0.9)

                ax_loglog.legend(fontsize=9)

        ax_loglog.set_title("Tail diagnostics (Smoothed series)")
        ax_loglog.set_xlabel("|Δz - loc|")
        ax_loglog.set_ylabel("CCDF")
        ax_loglog.set_xlim(left=1e-2)
        ax_loglog.set_ylim(bottom=1e-3)

        # 4. Stimulus Position (Wrapped Gauss)
        sns.histplot(z_list, bins=50, binrange=(low_z, high_z), kde=False, stat="density", ax=ax_dist_z, color="orange", alpha=0.3)
        ax_dist_z.set_title("Stimulus Position z")
        if np.isfinite(mean_z) and np.isfinite(var_z) and var_z > 0:
            x_z = np.linspace(low_z, high_z, 300)
            y_z = wrapped_gaussian_pdf_1d(x_z, mean_z, var_z, low=low_z, high=high_z)
            ax_dist_z.plot(x_z, y_z, "r-", lw=3, label=f"Wrapped Gauss (μ={mean_z:.2f}, σ={std_z:.2f})")
            ax_dist_z.legend()
            ax_dist_z.set_xlim(low_z, high_z) 

        os.makedirs(output_dir, exist_ok=True)
        fig_dist_path = os.path.join(output_dir, f"diagnostics_1d_Rf_{Rf:.6f}_k_{k:.7f}_wee_{wee:.6f}.svg")
        fig.savefig(fig_dist_path, dpi=150, bbox_inches="tight")
        plt.close(fig)

    return {
        "kappa_fit_decoded": kappa_fit,
        "theta_fit_decoded": theta_fit,
        "nu_fit_decoded_dz": t_evt.get("nu", np.nan) if t_evt else np.nan,
        "alpha_evt_dz": t_evt.get("alpha_evt", np.nan) if t_evt else np.nan,
        "slope_evt_dz": t_evt.get("linear_slope", np.nan) if t_evt else np.nan,
        "fig_dist_path": fig_dist_path,
        "aic_U_gamma": aic_U,
        "aic_U_gauss": aic_U_gauss,
        "aic_z_gauss": aic_z,
        "aic_dz_gauss": aic_dz,
        "aic_dz_t": t_evt.get("aic", np.nan) if t_evt else np.nan
    }


def fit_wrapped_gaussian_1d(z, low=-180.0, high=180.0):
    """Fit a wrapped Gaussian (wrapped normal) from samples on a ring [low, high)."""
    z = np.asarray(z, dtype=float)
    z = z[np.isfinite(z)]
    if z.size < 20:
        return np.nan, np.nan, np.nan

    L = high - low
    z_wrap = ((z - low) % L) + low

    theta = 2.0 * np.pi * (z_wrap - low) / L

    C = np.mean(np.cos(theta))
    S = np.mean(np.sin(theta))
    R = np.sqrt(C**2 + S**2)
    R = np.clip(R, 1e-12, 1.0)

    mu_theta = np.arctan2(S, C)
    if mu_theta < 0:
        mu_theta += 2.0 * np.pi

    sigma2_theta = -2.0 * np.log(R)
    mu_wrap = low + (L * mu_theta / (2.0 * np.pi))
    var_wrap = (L / (2.0 * np.pi))**2 * sigma2_theta
    std_wrap = np.sqrt(var_wrap)

    return float(mu_wrap), float(var_wrap), float(std_wrap)


def wrapped_gaussian_pdf_1d(x, mu, var, low=-180.0, high=180.0, n_wraps=7):
    """Wrapped Gaussian density on [low, high)."""
    x = np.asarray(x, dtype=float)
    L = high - low
    var = max(float(var), 1e-12)

    mu = ((mu - low) % L) + low
    k = np.arange(-n_wraps, n_wraps + 1, dtype=float)

    diff = x[..., None] - mu + k * L
    pdf = np.exp(-0.5 * diff**2 / var) / np.sqrt(2.0 * np.pi * var)
    return np.sum(pdf, axis=-1)


def simulate_plot_and_save_task_1d(args):
    params, Rf, wee, k, raw_data_dir, plot_dir = args
    filename = f"raw_data_1d_Rf_{Rf:.6f}_k_{k:.7f}_wee_{wee:.6f}.npz"
    save_path = os.path.join(raw_data_dir, filename)

    if is_valid_npz(save_path):
        return save_path

    if os.path.exists(save_path):
        try:
            os.remove(save_path)
        except OSError:
            pass

    try:
        local_params = params.copy()
        local_params["k"] = k
        local_params["normalization_k"] = k   

        simulator = CANNSimulator(local_params)
        results = simulator.run_simulation(
            Rf=Rf,
            Wei=0,
            ff_scale=local_params.get("feedforward_scale", 0.71648),
            Wee=wee,
        )

        delta_z = results["delta_z"]
        delta_z_values = np.insert(delta_z, 0, np.nan)
        time_array = np.linspace(0, local_params["simulation_time"], len(delta_z_values))

        generate_and_save_heatmaps_1d(
            time_array=time_array,
            Oc=results.get("E_FR"),
            U_field=results.get("Synaptic_Input"),
            params=local_params,
            Rf=Rf,
            wee=wee,
            k=k,
            output_dir=plot_dir,
        )

        safe_save_npz(
            save_path,
            time=time_array,
            z_list=results["E_stim"],
            Ue_list=results["E_bump_height"],
            delta_z=delta_z_values,
            Rf=np.array(Rf),
            k=np.array(k),
            wee=np.array(wee),
        )

        del results
        del simulator
        del delta_z
        del delta_z_values
        del time_array
        del local_params
        
        plt.close('all')
        gc.collect()

        return save_path

    except Exception as e:
        print(f"Simulation Error (Rf={Rf:.6f}, k={k:.7f}, wee={wee:.6f}): {e}")
        plt.close('all')
        gc.collect()
        return None


def generate_simulation_data_1d(params, target_wee, Rf_list, k_list, n_jobs=-1, raw_data_dir='output_net/raw_npz/1d/', plot_dir='output_net/plots/1d/'):
    os.makedirs(raw_data_dir, exist_ok=True)
    os.makedirs(plot_dir, exist_ok=True)
    max_workers = mp.cpu_count() if n_jobs == -1 else n_jobs
    
    tasks = [(params, Rf, target_wee, k, raw_data_dir, plot_dir) for Rf in Rf_list for k in k_list]
    saved_paths = []
    
    print(f"🚀 [1D] Starting {len(tasks)} simulations across {max_workers} CPU cores...")
    with concurrent.futures.ProcessPoolExecutor(max_workers=max_workers) as executor:
        futures = [executor.submit(simulate_plot_and_save_task_1d, task) for task in tasks]
        for future in tqdm(concurrent.futures.as_completed(futures), total=len(futures), desc="Simulating 1D"):
            try:
                res = future.result() 
                if res is not None: saved_paths.append(res)
            except Exception as e:
                print(f"\nWorker process crashed: {e}")
                
    return saved_paths


def analyze_saved_data_task_1d(args):
    data_path, params, plot_output_dir, save_diagnostic_plot, plot_range = args
    try:
        with np.load(data_path, allow_pickle=True) as data:
            Rf = float(data['Rf'])
            wee = float(data['wee'])
            k = float(data['k'])

            out = perform_integrated_analysis_1d(
                time=data['time'],
                z_list=data['z_list'],
                Ue_list=data['Ue_list'],
                delta_z=data['delta_z'],
                params=params,
                Rf=Rf,
                wee=wee,
                k=k,
                output_dir=plot_output_dir,
                save_diagnostic_plot=save_diagnostic_plot,
                plot_range=180
            )

            return {
                'k': k,
                'Rf': Rf,
                'wee': wee,
                'alpha_eff': out.get('alpha_evt_dz', np.nan),
                'kappa_fit': out.get('kappa_fit_decoded', np.nan),
                'theta_fit_decoded': out.get('theta_fit_decoded', np.nan),
                'nu_fit_decoded_dz': out.get('nu_fit_decoded_dz', np.nan),
                'alpha_evt_dz': out.get('alpha_evt_dz', np.nan),
                'slope_evt_dz': out.get('slope_evt_dz', np.nan),
                'fig_dist_path': out.get('fig_dist_path', ''),
                'aic_U_gamma': out.get('aic_U_gamma', np.nan),
                'aic_U_gauss': out.get('aic_U_gauss', np.nan),
                'aic_z_gauss': out.get('aic_z_gauss', np.nan),
                'aic_dz_gauss': out.get('aic_dz_gauss', np.nan),
                'aic_dz_t': out.get('aic_dz_t', np.nan),
            }
    except Exception as e:
        print(f"Analysis Error on {os.path.basename(data_path)}: {e}")
        return None


def run_offline_analysis_1d(data_paths, params, target_wee, k_list=None, n_jobs=-1, plot_dir='output_net/plots_1d/', plot_range=180):
    if not data_paths:
        return None

    os.makedirs(plot_dir, exist_ok=True)
    max_workers = mp.cpu_count() if n_jobs == -1 else n_jobs
    tasks = [(path, params, plot_dir, plot_range) for path in data_paths]
    results = []

    print(f"📊 [1D] Analyzing {len(tasks)} datasets across {max_workers} CPU cores.")
    with concurrent.futures.ProcessPoolExecutor(max_workers=max_workers) as executor:
        futures = [executor.submit(analyze_saved_data_task_1d, task) for task in tasks]
        for future in tqdm(concurrent.futures.as_completed(futures), total=len(futures), desc="Analyzing 1D"):
            try:
                res = future.result()
                if res is not None:
                    results.append(res)
            except Exception as e:
                print(f"\nAnalysis worker crashed: {e}")

    if not results:
        return None

    summary_df = pd.DataFrame(results).sort_values(by=['Rf', 'k'])
    csv_path = os.path.join(plot_dir, f'summary_1d_fixed_wee_{target_wee:.6f}.csv')
    summary_df.to_csv(csv_path, index=False)
    print(f"✅ [1D] Summary CSV successfully saved to {csv_path}")
    return summary_df


def _simulate_for_k(k_val, params, Rf, wee):
    """Top-level helper function to run the simulator for a specific k value."""
    local_params = params.copy()
    local_params['normalization_k'] = k_val
    local_params['k'] = k_val 
    
    simulator = CANNSimulator(local_params)
    simulator.initialize_network()
    
    res = simulator.run_simulation(
        Rf=Rf, 
        Wei=0, 
        ff_scale=local_params.get('feedforward_scale', 0.71648), 
        k=k_val,
        Wee=wee,
        noise_type='free'
    )
    return res


def _simulate_for_wee(wee_val, params, Rf, k_val):
    """Top-level helper function to run the simulator for a specific wee value."""
    local_params = params.copy()
    local_params['normalization_k'] = k_val
    local_params['k'] = k_val
    
    simulator = CANNSimulator(local_params)
    simulator.initialize_network()
    
    res = simulator.run_simulation(
        Rf=Rf, 
        Wei=0, 
        ff_scale=local_params.get('feedforward_scale', 0.71648), 
        k=k_val,
        Wee=wee_val,
        noise_type='free'
    )
    return res


def plot_R_U2_vary_wee(params, k_val, Rf, recording_start=50, dt=0.01, wee_list=None):
    """Generates steady-state U_E and R_E by varying wee while mapping superlinear region."""
    params['simulation_time'] = 500
    a = params['gaussian_width_exc']
    
    sim_temp = CANNSimulator(params)
    critical_weight = sim_temp.params['critical_weight']
    rho = sim_temp.params.get('neuron_density', 0.5) 
    
    Ue_steady_list = []
    Relist_steady_list = []
    
    if wee_list is None:
        wee_list = np.linspace(1.0, 3.0, 30)

    print(f"Running simulations for {len(wee_list)} values of wee with fixed Rf={Rf}, k={k_val}...")

    with mp.get_context('spawn').Pool() as pool:
        results = pool.starmap(
            _simulate_for_wee,
            [(wee_val, params, Rf, k_val) for wee_val in wee_list]
        )
        
    for single_run_result in results:
        Ue_timeseries = single_run_result['E_bump_height']
        Relist_timeseries = np.sum(single_run_result['E_FR'], axis=0) / (np.sqrt(np.pi) * 2 * a * rho)
        
        start_index = int(recording_start / dt)
        Ue_steady = np.average(Ue_timeseries[start_index:])
        Relist_steady = np.average(Relist_timeseries[start_index:])

        Ue_steady_list.append(Ue_steady)
        Relist_steady_list.append(Relist_steady)

    os.makedirs('output_net/plots', exist_ok=True)
    
    plt.figure(figsize=(10, 6))
    plt.plot(wee_list/critical_weight, Ue_steady_list, marker='o', linestyle='-', label=r'Steady-state $U_E$')
    plt.plot(wee_list/critical_weight, Relist_steady_list, marker='x', linestyle='--', label=r'Steady-state $R_E$')
    plt.xlabel(r'Excitatory coupling $w_{ee}$')
    plt.ylabel('Steady-state value')
    plt.title(f'Steady States vs wee (k={k_val}, Rf={Rf})')
    plt.legend()
    plt.grid(True)
    plt.savefig(f"output_net/plots/steady_states_vs_wee_k_{k_val}_Rf_{Rf}.svg", dpi=300, bbox_inches='tight')
    plt.close()

    csv_filename = f'output_net/plots/steady_state_wee_Ue_Re_k_{k_val}_Rf_{Rf}.csv'
    pd.DataFrame({
        'wee': wee_list,
        'Rf': Rf,
        'k': k_val,
        'Ue_steady': Ue_steady_list,
        'Re_steady': Relist_steady_list
    }).to_csv(csv_filename, index=False)
    
    wee_arr = np.asarray(wee_list, dtype=float)
    Re_arr = np.asarray(Relist_steady_list, dtype=float)
    Ue_arr = np.asarray(Ue_steady_list, dtype=float)

    start_Ue, end_Ue, info = find_superlinear_region_sigmoid(
        x=Ue_arr, y=Re_arr,
        smooth_window=9, polyorder=3,
        slope_on_frac=0,  
        min_run=3
    )

    if start_Ue is None or end_Ue is None:
        print("[WARN] superlinear region not detected.")
    else:
        sort_idx = np.argsort(Ue_arr)
        Ue_sorted = Ue_arr[sort_idx]
        wee_sorted = wee_arr[sort_idx]
        Re_sorted = Re_arr[sort_idx]

        start_wee = float(np.interp(start_Ue, Ue_sorted, wee_sorted))
        end_wee = float(np.interp(end_Ue, Ue_sorted, wee_sorted))

        start_Re = float(np.interp(start_Ue, Ue_sorted, Re_sorted))
        end_Re = float(np.interp(end_Ue, Ue_sorted, Re_sorted))

        nbins = 40
        R_bin_edges = np.linspace(start_Re, end_Re, nbins + 1)
        R_centers = 0.5 * (R_bin_edges[:-1] + R_bin_edges[1:])

        mask = (Ue_arr >= start_Ue) & (Ue_arr <= end_Ue)
        Re_supra = Re_arr[mask]
        wee_supra = wee_arr[mask]
        Ue_supra = Ue_arr[mask]

        sort_supra = np.argsort(Re_supra)
        Re_supra, wee_supra, Ue_supra = Re_supra[sort_supra], wee_supra[sort_supra], Ue_supra[sort_supra]

        wee_series = np.interp(R_centers, Re_supra, wee_supra, left=np.nan, right=np.nan)
        Ue_on_R = np.interp(R_centers, Re_supra, Ue_supra, left=np.nan, right=np.nan)

        print("\n=== superlinear mapping: fixed R bins -> corresponding wee ===")
        print(f"superlinear U_E range: [{start_Ue:.6g}, {end_Ue:.6g}]")
        print(f"superlinear wee range: [{start_wee:.6g}, {end_wee:.6g}]")
        
        df_map = pd.DataFrame({
            "R_center": R_centers,
            "wee_superlinear": wee_series,
            "Ue_superlinear": Ue_on_R,
            "k": k_val,
            "Rf": Rf
        })
        df_map.to_csv(f"output_net/plots/superlinear_Rbins_to_wee_k_{k_val}_Rf_{Rf}.csv", index=False)

        plt.figure(figsize=(10, 6))
        plt.plot(R_centers, wee_series, marker='o', linestyle='-')
        plt.xlabel(r'Fixed $R_E$ bin center')
        plt.ylabel(r'Corresponding $w_{ee}$ (superlinear only)')
        plt.title(r'New scale: $w_{ee}$ reparameterized by fixed $R_E$ bins (superlinear)')
        plt.grid(True)
        plt.savefig(f"output_net/plots/new_scale_wee_vs_fixed_Re_k_{k_val}_Rf_{Rf}.svg", dpi=300, bbox_inches='tight')
        plt.close()

        plt.figure(figsize=(10, 6))
        plt.plot(R_centers, Ue_on_R, marker='x', linestyle='--')
        plt.xlabel(r'Fixed $R_E$ bin center')
        plt.ylabel(r'$U_E$ (superlinear branch)')
        plt.title(r'superlinear branch reparameterized by fixed $R_E$ bins: $U_E(R_E)$')
        plt.grid(True)
        plt.savefig(f"output_net/plots/new_scale_Ue_vs_fixed_Re_k_{k_val}_Rf_{Rf}.svg", dpi=300, bbox_inches='tight')
        plt.close()


def _odd_window(n, max_n):
    n = int(n)
    if n < 3: n = 3
    if n % 2 == 0: n += 1
    if n > max_n:
        n = max_n if max_n % 2 == 1 else max_n - 1
    return max(3, n)


def find_superlinear_region_sigmoid(
    x, y,
    smooth_window=9,
    polyorder=3,
    slope_on_frac=0.25,
    min_run=3
):
    """Detects early superlinear region for a sigmoid-like y(x)."""
    x = np.asarray(x, dtype=float)
    y = np.asarray(y, dtype=float)

    order = np.argsort(x)
    x = x[order]; y = y[order]
    finite = np.isfinite(x) & np.isfinite(y)
    x = x[finite]; y = y[finite]
    if len(x) < 7:
        return None, None, {"reason": "too_few_points"}

    w = _odd_window(smooth_window, len(y))
    if w <= polyorder:
        polyorder = max(1, min(3, w - 1))
    y_s = savgol_filter(y, window_length=w, polyorder=polyorder, mode="interp")

    dy = np.gradient(y_s, x)
    d2y = np.gradient(dy, x)

    i_inf = int(np.nanargmax(dy))
    max_slope = float(dy[i_inf])
    if not np.isfinite(max_slope) or max_slope <= 0:
        return None, None, {"reason": "nonpositive_max_slope"}

    slope_thr = slope_on_frac * max_slope

    cand = (np.arange(len(x)) <= i_inf) & (d2y > 0) & (dy >= slope_thr)
    idx = np.where(cand)[0]
    if len(idx) == 0:
        return None, None, {"reason": "no_candidates", "i_inf": i_inf}

    runs = []
    start = idx[0]
    prev = idx[0]
    for k in idx[1:]:
        if k == prev + 1:
            prev = k
        else:
            runs.append((start, prev))
            start = k
            prev = k
    runs.append((start, prev))

    runs_sorted = sorted(runs, key=lambda ab: (ab[1]-ab[0], ab[1]), reverse=True)
    start_i, end_i = runs_sorted[0]
    if (end_i - start_i + 1) < min_run:
        return None, None, {"reason": "run_too_short", "i_inf": i_inf, "run": (start_i, end_i)}

    end_i = min(end_i, i_inf)

    if start_i >= end_i:
        return None, None, {"reason": "start_ge_end", "i_inf": i_inf, "run": (start_i, end_i)}

    info = {"x_sorted": x, "y_smooth": y_s, "dy": dy, "d2y": d2y, "i_inf": i_inf, "max_slope": max_slope}
    return float(x[start_i]), float(x[end_i]), info


def plot_R_U2_vary_k(params, wee, Rf, recording_start=50, dt=0.01, k_list=None):
    """Generates steady-state U_E and R_E by varying normalization constant k."""
    default_k = 0.0005
    params['simulation_time'] = 500
    a = params['gaussian_width_exc']
    
    sim_temp = CANNSimulator(params)
    rho = sim_temp.params.get('neuron_density', 0.5) 
    
    Ue_steady_list = []
    Relist_steady_list = []

    print(f"Running simulations for {len(k_list)} values of k with fixed Rf={Rf}, wee={wee}...")

    with mp.get_context('spawn').Pool() as pool:
        results = pool.starmap(
            _simulate_for_k,
            [(k_val, params, Rf, wee) for k_val in k_list]
        )
        
    for single_run_result in results:
        Ue_timeseries = single_run_result['E_bump_height']
        Relist_timeseries = np.sum(single_run_result['E_FR'], axis=0) / (np.sqrt(np.pi) * 2 * a * rho)
        
        start_index = int(recording_start / dt)
        Ue_steady = np.average(Ue_timeseries[start_index:])
        Relist_steady = np.average(Relist_timeseries[start_index:])

        Ue_steady_list.append(Ue_steady)
        Relist_steady_list.append(Relist_steady)

    os.makedirs('output_net', exist_ok=True)
    plt.figure(figsize=(10, 6))
    plt.plot(k_list/default_k, Ue_steady_list, marker='o', linestyle='-', label=r'Steady-state $U_E$')
    plt.plot(k_list/default_k, Relist_steady_list, marker='x', linestyle='--', label=r'Steady-state $R_E$')
    plt.xlabel(r'Normalization $k$ (relative to default) default $k={:.1e}$'.format(default_k))
    plt.ylabel('Steady-state value')
    plt.title(f'Steady States vs Normalization k (wee={wee}, Rf={Rf})')
    plt.legend()
    plt.grid(True)
    plt.savefig(f"output_net/steady_states_vs_k_wee_{wee}_Rf_{Rf}.svg", dpi=300, bbox_inches='tight')

    csv_filename = f'output_net/steady_state_k_Ue_Re_wee_{wee}_Rf_{Rf}.csv'
    pd.DataFrame({
        'k': k_list,
        'Rf': Rf,
        'wee': wee,
        'Ue_steady': Ue_steady_list,
        'Re_steady': Relist_steady_list
    }).to_csv(csv_filename, index=False)
    
    k_arr = np.asarray(k_list, dtype=float)
    Re_arr = np.asarray(Relist_steady_list, dtype=float)
    Ue_arr = np.asarray(Ue_steady_list, dtype=float)

    start_Ue, end_Ue, info = find_superlinear_region_sigmoid(
        x=Ue_arr, y=Re_arr,
        smooth_window=9, polyorder=3,
        slope_on_frac=0,  
        min_run=3
    )

    if start_Ue is None or end_Ue is None:
        print("[WARN] superlinear region not detected.")
    else:
        sort_idx = np.argsort(Ue_arr)
        Ue_sorted = Ue_arr[sort_idx]
        k_sorted = k_arr[sort_idx]
        Re_sorted = Re_arr[sort_idx]

        start_k = float(np.interp(start_Ue, Ue_sorted, k_sorted))
        end_k = float(np.interp(end_Ue, Ue_sorted, k_sorted))

        start_Re = float(np.interp(start_Ue, Ue_sorted, Re_sorted))
        end_Re = float(np.interp(end_Ue, Ue_sorted, Re_sorted))

        nbins = 40
        R_bin_edges = np.linspace(start_Re, end_Re, nbins + 1)
        R_centers = 0.5 * (R_bin_edges[:-1] + R_bin_edges[1:])

        mask = (Ue_arr >= start_Ue) & (Ue_arr <= end_Ue)
        Re_supra = Re_arr[mask]
        k_supra = k_arr[mask]
        Ue_supra = Ue_arr[mask]

        sort_supra = np.argsort(Re_supra)
        Re_supra, k_supra, Ue_supra = Re_supra[sort_supra], k_supra[sort_supra], Ue_supra[sort_supra]

        k_series = np.interp(R_centers, Re_supra, k_supra, left=np.nan, right=np.nan)
        Ue_on_R = np.interp(R_centers, Re_supra, Ue_supra, left=np.nan, right=np.nan)

        print("\n=== superlinear mapping: fixed R bins -> corresponding k ===")
        print(f"superlinear U_E range: [{start_Ue:.6g}, {end_Ue:.6g}]")
        print(f"superlinear k range: [{start_k:.6g}, {end_k:.6g}]")
        
        df_map = pd.DataFrame({
            "R_center": R_centers,
            "k_superlinear": k_series,
            "Ue_superlinear": Ue_on_R,
            "wee": wee,
            "Rf": Rf
        })
        df_map.to_csv(f"output_net/plots/superlinear_Rbins_to_k_wee_{wee}_Rf_{Rf}.csv", index=False)

        plt.figure(figsize=(10, 6))
        plt.plot(R_centers, k_series/default_k, marker='o', linestyle='-')
        plt.xlabel(r'Fixed $R_E$ bin center')
        plt.ylabel(r'Corresponding $k$ (superlinear only) (relative to default)')
        plt.title('New scale: $k$ reparameterized by fixed $R_E$ bins (superlinear)')
        plt.grid(True)
        plt.savefig(f"output_net/plots/new_scale_k_vs_fixed_Re_wee_{wee}_Rf_{Rf}.svg", dpi=300, bbox_inches='tight')
        plt.close()

        plt.figure(figsize=(10, 6))
        plt.plot(R_centers, Ue_on_R, marker='x', linestyle='--')
        plt.xlabel(r'Fixed $R_E$ bin center')
        plt.ylabel(r'$U_E$ (superlinear branch)')
        plt.title('superlinear branch reparameterized by fixed $R_E$ bins: $U_E(R_E)$')
        plt.grid(True)
        plt.savefig(f"output_net/plots/new_scale_Ue_vs_fixed_Re_wee_{wee}_Rf_{Rf}.svg", dpi=300, bbox_inches='tight')
        plt.close()


def _collect_or_build_summary_1d(
    params,
    target_wee,
    Rf_list,
    k_list,
    raw_data_dir='output_net/raw_npz/1d/',
    plot_dir='output_net/plots/1d/',
    n_jobs=-1,
    force_reanalyze=False,
    plot_range=180
):
    os.makedirs(raw_data_dir, exist_ok=True)
    os.makedirs(plot_dir, exist_ok=True)

    target_pairs = {(float(Rf), float(k)) for Rf in Rf_list for k in k_list}
    required_cols = {'Rf', 'k', 'aic_U_gamma', 'aic_U_gauss', 'aic_dz_t', 'aic_dz_gauss', 'slope_evt_dz'}

    df_parts = []
    if not force_reanalyze:
        for f in glob.glob(os.path.join(plot_dir, 'summary_1d_fixed_wee_*.csv')):
            try:
                tmp = pd.read_csv(f)
                if required_cols.issubset(tmp.columns):
                    if 'wee' in tmp.columns:
                        tmp = tmp[np.isclose(tmp['wee'], target_wee)]
                    df_parts.append(tmp)
            except Exception as e:
                print(f"Warning: failed to read {f}: {e}")

    df = pd.concat(df_parts, ignore_index=True) if df_parts else pd.DataFrame()

    if not df.empty:
        subset_cols = ['Rf', 'k', 'wee'] if 'wee' in df.columns else ['Rf', 'k']
        df = df.drop_duplicates(subset=subset_cols, keep='last')
        have_pairs = {(float(rf), float(k)) for rf, k in zip(df['Rf'], df['k'])}
    else:
        have_pairs = set()

    missing_pairs = sorted(target_pairs - have_pairs)

    if missing_pairs:
        print(f"Need {len(missing_pairs)} missing (Rf, k) cases.")

        missing_sim_tasks = []
        for Rf, k in missing_pairs:
            fn = f'raw_data_1d_Rf_{Rf:.6f}_k_{k:.7f}_wee_{target_wee:.6f}.npz'
            p = os.path.join(raw_data_dir, fn)

            if not is_valid_npz(p):
                if os.path.exists(p):
                    print(f"Removing corrupted raw file: {p}")
                    try:
                        os.remove(p)
                    except OSError as e:
                        print(f"Warning: could not remove {p}: {e}")
                missing_sim_tasks.append((params, Rf, target_wee, k, raw_data_dir, plot_dir))

        if missing_sim_tasks:
            print(f"Running {len(missing_sim_tasks)} missing/corrupted simulations...")
            max_workers = mp.cpu_count() if n_jobs == -1 else n_jobs
            with concurrent.futures.ProcessPoolExecutor(max_workers=max_workers) as executor:
                futures = [executor.submit(simulate_plot_and_save_task_1d, task) for task in missing_sim_tasks]
                for future in tqdm(concurrent.futures.as_completed(futures), total=len(futures), desc="Simulating 1D"):
                    try:
                        future.result()
                    except Exception as e:
                        print(f"\nSimulation worker crashed: {e}")

        all_npz_paths = []
        for Rf, k in target_pairs:
            fn = f'raw_data_1d_Rf_{Rf:.6f}_k_{k:.7f}_wee_{target_wee:.6f}.npz'
            p = os.path.join(raw_data_dir, fn)
            if is_valid_npz(p):
                all_npz_paths.append(p)

        if all_npz_paths:
            print(f"Analyzing {len(all_npz_paths)} raw datasets...")
            max_workers = mp.cpu_count() if n_jobs == -1 else n_jobs
            tasks = [(path, params, plot_dir, plot_range) for path in all_npz_paths]
            results = []

            with concurrent.futures.ProcessPoolExecutor(max_workers=max_workers) as executor:
                futures = [executor.submit(analyze_saved_data_task_1d, task) for task in tasks]
                for future in tqdm(concurrent.futures.as_completed(futures), total=len(futures), desc="Analyzing 1D"):
                    try:
                        res = future.result()
                        if res is not None:
                            results.append(res)
                    except Exception as e:
                        print(f"\nAnalysis worker crashed: {e}")

            if results:
                df_new = pd.DataFrame(results).sort_values(['Rf', 'k'])
                summary_path = os.path.join(plot_dir, f'summary_1d_fixed_wee_{target_wee:.6f}_grid.csv')
                df_new.to_csv(summary_path, index=False)
                print(f"Saved merged summary to {summary_path}")

                df = pd.concat([df, df_new], ignore_index=True) if not df.empty else df_new
                subset_cols = ['Rf', 'k', 'wee'] if 'wee' in df.columns else ['Rf', 'k']
                df = df.drop_duplicates(subset=subset_cols, keep='last')

    if df.empty:
        raise ValueError("No valid summary data available after checking CSV, NPZ, and simulations.")

    df = df[df.apply(lambda row: (float(row['Rf']), float(row['k'])) in target_pairs, axis=1)].copy()
    if 'wee' in df.columns:
        df = df[np.isclose(df['wee'], target_wee)]

    return df.sort_values(['Rf', 'k']).reset_index(drop=True)


def signed_log10(x):
    x = np.asarray(x, dtype=float)
    return np.sign(x) * np.log10(1 + np.abs(x))


def plot_signedlog_heatmap(pivot, ax, center=0.0, cbar_label='signed log10(1+|ΔAIC|)'):
    z = signed_log10(pivot.to_numpy())
    lim = np.nanmax(np.abs(z[np.isfinite(z)]))
    if not np.isfinite(lim) or lim == 0:
        lim = 1.0

    cmap = LinearSegmentedColormap.from_list(
        "blue_white_red",
        ["blue", "white", "red"]
    )
    norm = TwoSlopeNorm(vmin=-lim, vcenter=center, vmax=lim)

    sns.heatmap(
        pd.DataFrame(z, index=pivot.index, columns=pivot.columns),
        ax=ax,
        cmap=cmap,
        norm=norm,
        annot=False,
        cbar_kws={'label': cbar_label}
    )
    ax.invert_yaxis()


def plot_bof_vs_k_rf(
    params,
    target_wee,
    Rf_list,
    k_list,
    raw_data_dir='output_net/raw_npz/1d/',
    plot_dir='output_net/plots/1d/',
    n_jobs=-1,
    metric='delta_aic',
    force_reanalyze=False,
    save=True,
):
    """
    Heatmaps across (Rf, k):
      - U: Gaussian vs Gamma
      - dz: Gaussian vs Student-t
    """
    df = _collect_or_build_summary_1d(
        params=params, target_wee=target_wee, Rf_list=Rf_list, k_list=k_list,
        raw_data_dir=raw_data_dir, plot_dir=plot_dir, n_jobs=n_jobs, force_reanalyze=force_reanalyze,
    )

    req = ['Rf', 'k', 'aic_U_gamma', 'aic_U_gauss', 'aic_dz_t', 'aic_dz_gauss']
    df = df.dropna(subset=req).copy()
    if df.empty:
        raise ValueError("No valid rows available for best-of-fit plotting.")

    df['delta_aic_U'] = - df['aic_U_gauss'] + df['aic_U_gamma']
    df['delta_aic_dz'] = - df['aic_dz_gauss'] + df['aic_dz_t']
    df['winner_U'] = (df['delta_aic_U'] < 0).astype(float)
    df['winner_dz'] = (df['delta_aic_dz'] < 0).astype(float)

    if metric == 'delta_aic':
        valU, valdz = 'delta_aic_U', 'delta_aic_dz'
        titleU = r'U fit: $\Delta$AIC = AIC(Gamma)- AIC(Gaussian)'
        titledz = r'$\Delta z$ fit: $\Delta$AIC = IC(Student-t) - AIC(Gaussian)'
        fmt = '.2f'
    elif metric == 'winner':
        valU, valdz = 'winner_U', 'winner_dz'
        titleU = 'U fit winner (1 = Gamma better)'
        titledz = 'Δz fit winner (1 = Student-t better)'
        fmt = '.0f'
    else:
        raise ValueError("metric must be 'delta_aic' or 'winner'")

    pivot_U = df.pivot_table(index='Rf', columns='k', values=valU, aggfunc='mean').sort_index()
    pivot_dz = df.pivot_table(index='Rf', columns='k', values=valdz, aggfunc='mean').sort_index()

    fig, axes = plt.subplots(1, 2, figsize=(15, 6), constrained_layout=True)

    if metric == 'delta_aic':
        limit_U = np.nanpercentile(np.abs(pivot_U.values), 95)
        limit_dz = np.nanpercentile(np.abs(pivot_dz.values), 95)
        
        limit_U = max(limit_U, 1.0)
        limit_dz = max(limit_dz, 1.0)

        sns.heatmap(
            pivot_U, ax=axes[0], cmap='RdBu_r', center=0,
            vmin=-limit_U, vmax=limit_U, annot=False, fmt=fmt, cbar_kws={'label': r'$\Delta$AIC'}
        )
        sns.heatmap(
            pivot_dz, ax=axes[1], cmap='RdBu_r', center=0,
            vmin=-limit_dz, vmax=limit_dz, annot=False, fmt=fmt, cbar_kws={'label': r'$\Delta$AIC'}
        )
    else:
        sns.heatmap(
            pivot_U, ax=axes[0], cmap='RdBu_r', center=0.5,
            vmin=0, vmax=1, annot=False, fmt=fmt, cbar_kws={'label': 'Winner'}
        )
        sns.heatmap(
            pivot_dz, ax=axes[1], cmap='RdBu_r', center=0.5,
            vmin=0, vmax=1, annot=False, fmt=fmt, cbar_kws={'label': 'Winner'}
        )

    axes[0].set_title(titleU)
    axes[0].set_xlabel('k')
    axes[0].set_ylabel('Rf')
    axes[0].invert_yaxis()

    axes[1].set_title(titledz)
    axes[1].set_xlabel('k')
    axes[1].set_ylabel('Rf')
    axes[1].invert_yaxis()

    fig.suptitle(f'Best-of-fit comparison across k and Rf (wee={target_wee:.6f})', fontsize=16)

    if save:
        out = os.path.join(plot_dir, f'bof_vs_k_rf_{metric}_wee_{target_wee:.6f}.svg')
        fig.savefig(out, dpi=150, bbox_inches='tight')
        print(f"Saved: {out}")

    return fig, df, pivot_U, pivot_dz


def plot_tail_slope_vs_k_rf(
    params,
    target_wee,
    Rf_list,
    k_list,
    raw_data_dir='output_net/raw_npz/1d/',
    plot_dir='output_net/plots/1d/',
    n_jobs=-1,
    value_col='slope_evt_dz',
    force_reanalyze=False,
    save=True,
):
    df = _collect_or_build_summary_1d(
        params=params, target_wee=target_wee, Rf_list=Rf_list, k_list=k_list,
        raw_data_dir=raw_data_dir, plot_dir=plot_dir, n_jobs=n_jobs, force_reanalyze=force_reanalyze,
    )

    if value_col not in df.columns:
        raise ValueError(f"{value_col} not found in summary dataframe.")

    df = df.dropna(subset=['Rf', 'k', value_col]).copy()
    if df.empty:
        raise ValueError(f"No valid rows available for {value_col}.")

    pivot = df.pivot_table(index='Rf', columns='k', values=value_col, aggfunc='mean').sort_index()

    fig, ax = plt.subplots(figsize=(8, 6), constrained_layout=True)

    spread = np.nanpercentile(np.abs(pivot.values - (-2.0)), 95)
    spread = max(spread, 1.0)

    norm = mcolors.TwoSlopeNorm(vcenter=-2.0, vmin=-2.0 - spread, vmax=-2.0 + spread)

    sns.heatmap(
        pivot,
        ax=ax,
        cmap='RdBu_r',
        norm=norm,
        annot=False,
        fmt='.2f',
        cbar_kws={'label': value_col}
    )

    ax.set_title(f'Step-size tail metric across k and Rf: {value_col} (wee={target_wee:.6f})')
    ax.set_xlabel('k')
    ax.set_ylabel('Rf')
    ax.invert_yaxis()

    if save:
        out = os.path.join(plot_dir, f'{value_col}_vs_k_rf_wee_{target_wee:.6f}.svg')
        fig.savefig(out, dpi=150, bbox_inches='tight')
        print(f"Saved: {out}")

    return fig, df, pivot


def _wrap_to_ring(z, position_min=-180.0, position_max=180.0):
    """Wrap scalar/array z back to the ring interval [position_min, position_max)."""
    L = position_max - position_min
    return ((z - position_min) % L) + position_min


def simulation_for_z_fixed_U(
    tau, rho, w_ef, sigma_z,
    total_time, dt,
    fixed_U,
    Rf,
    input_center=0.0,
    z0=0.0,
    position_min=-180.0,
    position_max=180.0,
    wrap_ring=True,
    rng=None
):
    """Simulate only z dynamics with bump height fixed at a constant value."""
    if rng is None:
        rng = np.random.default_rng()

    num_steps = int(np.round(total_time / dt)) + 1
    time = np.linspace(0, total_time, num_steps)
    z_list = np.zeros(num_steps, dtype=float)
    z_list[0] = z0

    U_const = max(1e-9, float(fixed_U))
    sqrt_dt = np.sqrt(dt)
    rho_sqrt2 = rho / np.sqrt(2.0)

    def _wrap_to_ring(z):
        L = position_max - position_min
        return ((z - position_min) % L) + position_min

    for i in range(num_steps - 1):
        z_curr = z_list[i]

        drift_z = (rho_sqrt2 * w_ef * Rf / (tau * U_const)) * (input_center - z_curr)
        noise_z = sigma_z / np.sqrt(tau * U_const) * rng.normal() * sqrt_dt

        z_next = z_curr + drift_z * dt + noise_z
        if wrap_ring:
            z_next = _wrap_to_ring(z_next)

        z_list[i + 1] = z_next

    Ue = np.full(num_steps, U_const, dtype=float)
    return time, Ue, z_list


def wrap_to_interval_1d(x, low=-180.0, high=180.0):
    """Wrap values onto [low, high)."""
    L = high - low
    return ((np.asarray(x) - low) % L) + low


def make_wrapped_target_1d(param_dict, Rf, nbins=180, n_wraps=7):
    low = float(param_dict.get('position_min', -180.0))
    high = float(param_dict.get('position_max', 180.0))

    bin_edges = np.linspace(low, high, nbins + 1)
    bin_centers = 0.5 * (bin_edges[:-1] + bin_edges[1:])

    mu_th = float(param_dict.get('input_position', 0.0))
    rho = param_dict['num_neurons'] / (param_dict['position_max'] - param_dict['position_min'])
    a = param_dict['gaussian_width_exc']

    var_th = 1.0 / (np.sqrt(2.0 * np.pi) * rho * Rf / a)
    var_th = max(var_th, 1e-12)

    q_density = wrapped_gaussian_pdf_1d(
        bin_centers,
        mu=mu_th,
        var=var_th,
        low=low,
        high=high,
        n_wraps=n_wraps,
        normalize=False
    )

    q_target = q_density / np.sum(q_density)
    q_target = np.maximum(q_target, 1e-300)
    q_target = q_target / np.sum(q_target)

    return bin_edges, bin_centers, q_target, mu_th, var_th


def compute_wrapped_kl_divergence_fast(
    param_dict,
    Se,
    Rf,
    mode='cumulative',
    nbins=180,
    n_wraps=5,
    pseudocount=0.0,
    return_target=False
):
    """KL divergence on a circular domain using a wrapped-Gaussian target."""
    Se = np.asarray(Se, dtype=float)
    if Se.ndim == 1:
        Se = Se[None, :]

    low = float(param_dict.get('position_min', -180.0))
    high = float(param_dict.get('position_max', 180.0))

    bin_edges, bin_centers, q_target, mu_th, var_th = make_wrapped_target_1d(
        param_dict, Rf, nbins=nbins, n_wraps=n_wraps
    )

    T = Se.shape[1]
    counts_t = np.zeros((T, nbins), dtype=float)

    for t in range(T):
        x = Se[:, t]
        x = x[np.isfinite(x)]
        if x.size == 0:
            continue
        x = wrap_to_interval_1d(x, low=low, high=high)
        counts_t[t], _ = np.histogram(x, bins=bin_edges)

    if mode == 'cumulative':
        counts = np.cumsum(counts_t, axis=0)
    elif mode == 'instantaneous':
        counts = counts_t
    else:
        raise ValueError("mode must be 'cumulative' or 'instantaneous'")

    if pseudocount > 0:
        counts = counts + pseudocount

    total = np.sum(counts, axis=1, keepdims=True)
    total = np.maximum(total, 1.0)

    p_emp = counts / total

    q = np.maximum(q_target, 1e-300)
    kl = np.zeros(T, dtype=float)

    for t in range(T):
        p = p_emp[t]
        mask = p > 0
        if np.any(mask):
            kl[t] = np.sum(p[mask] * (np.log(p[mask]) - np.log(q[mask])))

    if return_target:
        return kl, bin_centers, q_target

    return kl


def compute_kl_divergence_fast(param_dict, Se, Rf, mode='cumulative'):
    """Fast KL divergence between empirical z samples and the feedforward Gaussian target."""
    Se = np.asarray(Se, dtype=float)
    if Se.ndim == 1:
        Se = Se[None, :]

    rho = param_dict['num_neurons'] / (param_dict['position_max'] - param_dict['position_min'])
    a = param_dict['gaussian_width_exc']
    vth = 1.0 / (np.sqrt(2.0 * np.pi) * rho * Rf / a)
    mth = float(param_dict['input_position'])

    if mode == 'instantaneous':
        mean_t = np.nanmean(Se, axis=0)
        var_t = np.nanvar(Se, axis=0)
    elif mode == 'cumulative':
        finite_mask = np.isfinite(Se)
        x = np.where(finite_mask, Se, 0.0)

        col_sum = np.sum(x, axis=0)
        col_sq_sum = np.sum(x**2, axis=0)
        col_n = np.sum(finite_mask, axis=0)

        cum_sum = np.cumsum(col_sum)
        cum_sq_sum = np.cumsum(col_sq_sum)
        cum_n = np.cumsum(col_n)

        cum_n = np.maximum(cum_n, 1)
        mean_t = cum_sum / cum_n
        var_t = cum_sq_sum / cum_n - mean_t**2
    else:
        raise ValueError("mode must be 'cumulative' or 'instantaneous'")

    var_t = np.maximum(var_t, 1e-10)

    kl = 0.5 * (
        np.log(vth / var_t)
        + (var_t + (mean_t - mth)**2) / vth
        - 1.0
    )
    return kl


def plot_kl_div_with_lan_net(
    params,
    Rf,
    wee,
    k=None,
    num_trials=30,
    stride=10,
    mode='instantaneous',
    plot_dir='output_net/plots/1d/',
    save=True,
    seed=None
):
    """
    Compare KL-vs-time for:
      1) full network simulation (CANN)
      2) reduced z dynamics with fixed bump height U = <U>_network
    """
    os.makedirs(plot_dir, exist_ok=True)

    local_params = params.copy()
    if k is None:
        k = local_params.get('k', local_params.get('normalization_k', 0.0005))
    local_params['k'] = k

    tau = local_params['time_constant_exc']
    total_time = 10
    dt = local_params['time_step']
    a_e = local_params['gaussian_width_exc']
    rho = local_params['num_neurons'] / (local_params['position_max'] - local_params['position_min'])

    ff_scale = local_params.get('feedforward_scale', 0.71648)
    mu_in = float(local_params.get('input_position', 0.0))
    recording_start = 0
    local_params['simulation_time'] = total_time

    position_min = float(local_params.get('position_min', -180.0))
    position_max = float(local_params.get('position_max', 180.0))

    F = float(local_params.get('Fano_factor', 0.5))
    sigma_z = np.sqrt(8.0 * a_e * F / (3.0 * np.sqrt(3.0 * np.pi)))

    z_net_trials = []
    U_net_trials = []

    for trial in range(num_trials):
        simulator = CANNSimulator(local_params)
        results = simulator.run_simulation(
            Rf=Rf,
            Wei=0,
            ff_scale=ff_scale,
            Wee=wee
        )

        z_net = np.asarray(results['E_stim'], dtype=float)
        U_net = np.asarray(results['E_bump_height'], dtype=float)

        z_net_trials.append(z_net)
        U_net_trials.append(U_net)

    min_len_full = min(len(x) for x in z_net_trials)
    z_net_trials = np.vstack([x[:min_len_full] for x in z_net_trials])
    U_net_trials = np.vstack([x[:min_len_full] for x in U_net_trials])

    time_full = np.arange(min_len_full) * dt

    mask_rec = time_full >= recording_start
    if np.any(mask_rec):
        U_bar = float(np.nanmean(U_net_trials[:, mask_rec]))
    else:
        U_bar = float(np.nanmean(U_net_trials))

    z_net_trials = z_net_trials[:, ::stride]
    time_dec = time_full[::stride]

    z0 = float(np.nanmean(z_net_trials[:, 0]))

    z_lan_trials = []

    for trial in range(num_trials):
        rng = np.random.default_rng(None if seed is None else seed + trial)

        time_red, U_red, z_red = simulation_for_z_fixed_U(
            tau=tau,
            rho=rho,
            w_ef=ff_scale,
            sigma_z=sigma_z,
            total_time=total_time,
            dt=dt,
            fixed_U=U_bar,
            Rf=Rf,
            input_center=mu_in,
            z0=z0,
            position_min=position_min,
            position_max=position_max,
            wrap_ring=True,
            rng=rng
        )

        z_lan_trials.append(np.asarray(z_red, dtype=float)[:min_len_full:stride])

    z_lan_trials = np.vstack(z_lan_trials)

    min_len = min(z_net_trials.shape[1], z_lan_trials.shape[1], len(time_dec))
    z_net_trials = z_net_trials[:, :min_len]
    z_lan_trials = z_lan_trials[:, :min_len]
    time_dec = time_dec[:min_len]

    kl_net = compute_wrapped_kl_divergence_fast(
        local_params,
        z_net_trials,
        Rf=Rf,
        mode=mode,
        nbins=180,
        n_wraps=7
    )

    kl_lan = compute_wrapped_kl_divergence_fast(
        local_params,
        z_lan_trials,
        Rf=Rf,
        mode=mode,
        nbins=180,
        n_wraps=7
    )

    fig, ax = plt.subplots(figsize=(8, 5))
    ax.plot(time_dec, kl_net, lw=2, label='Network simulation')
    ax.plot(time_dec, kl_lan, lw=2, label=fr'Reduced SDE, fixed $U=\langle U \rangle$ = {U_bar:.6f}')
    ax.axvline(recording_start, linestyle='--', alpha=0.5, label='recording_start')

    ax.set_xlabel('Time')
    ax.set_ylabel('KL divergence')
    ax.set_title(
        f'KL vs time: network vs fixed-U reduced dynamics\n'
        f'Rf={Rf:.6f}, k={k:.7f}, wee={wee:.6f}, mode={mode}'
    )
    ax.grid(True, alpha=0.3)
    ax.legend()

    df_out = pd.DataFrame({
        'time': time_dec,
        'kl_network': kl_net,
        'kl_lan_fixedU': kl_lan
    })

    if save:
        fig_path = os.path.join(
            plot_dir,
            f'kl_div_net_vs_lan_Rf_{Rf:.6f}_k_{k:.7f}_wee_{wee:.6f}.svg'
        )
        csv_path = os.path.join(
            plot_dir,
            f'kl_div_net_vs_lan_Rf_{Rf:.6f}_k_{k:.7f}_wee_{wee:.6f}.csv'
        )
        fig.savefig(fig_path, dpi=150, bbox_inches='tight')
        df_out.to_csv(csv_path, index=False)
        print(f"Saved figure to: {fig_path}")
        print(f"Saved data to:   {csv_path}")

    return fig, df_out


def replot_kl_div_from_csv(csv_path, save_path=None):
    df = pd.read_csv(csv_path)
    time = df['time'].to_numpy()
    kl_net = df['kl_network'].to_numpy()
    kl_lan = df['kl_lan_fixedU'].to_numpy()
    
    mask = time <= 10
    fig, ax = plt.subplots(figsize=(8, 5))
    ax.plot(time[mask], kl_net[mask], lw=2, label='Network simulation')
    ax.plot(time[mask], kl_lan[mask], lw=2, label=r'Reduced SDE, fixed U')

    ax.set_xlabel('Time')
    ax.set_ylabel('KL divergence')
    ax.grid(True, alpha=0.3)
    ax.legend()

    if save_path is not None:
        fig.savefig(save_path, dpi=150, bbox_inches='tight')

    return fig


def _collect_or_build_summary_1d_fixed_k(
    params,
    target_k,
    Rf_list,
    wee_list,
    raw_data_dir='output_net/raw_npz/1d/',
    plot_dir='output_net/plots/1d/',
    n_jobs=-1,
    force_reanalyze=False,
    save_diagnostic_plot=False,
    save_summary_csv=True,
    atol_k=1e-12,
    plot_range=180
):
    """
    Return a clean summary dataframe for the requested (Rf, wee) grid at fixed k.
    Uses existing summary CSVs if possible; otherwise falls back to raw npz,
    and runs only genuinely missing simulations/analyses.
    """
    os.makedirs(raw_data_dir, exist_ok=True)
    os.makedirs(plot_dir, exist_ok=True)

    target_pairs = {
        _grid_key_1d(Rf, wee, rf_decimals=3, other_decimals=3)
        for Rf in Rf_list for wee in wee_list
    }
    required_cols = {
        'Rf', 'k', 'wee',
        'aic_U_gamma', 'aic_U_gauss',
        'aic_dz_t', 'aic_dz_gauss',
        'slope_evt_dz'
    }

    df_parts = []
    if not force_reanalyze:
        for f in glob.glob(os.path.join(plot_dir, 'summary_1d_fixed_k_*.csv')):
            try:
                tmp = pd.read_csv(f)
                if required_cols.issubset(tmp.columns):
                    tmp = tmp[np.isclose(tmp['k'], target_k, rtol=0, atol=atol_k)]
                    if not tmp.empty:
                        df_parts.append(tmp)
            except Exception as e:
                print(f"Warning: failed to read {f}: {e}")

        for f in glob.glob(os.path.join(plot_dir, 'summary_1d_fixed_wee_*.csv')):
            try:
                tmp = pd.read_csv(f)
                if required_cols.issubset(tmp.columns):
                    tmp = tmp[np.isclose(tmp['k'], target_k, rtol=0, atol=atol_k)]
                    if not tmp.empty:
                        df_parts.append(tmp)
            except Exception as e:
                print(f"Warning: failed to read {f}: {e}")

    df = pd.concat(df_parts, ignore_index=True) if df_parts else pd.DataFrame()

    if not df.empty:
        df = df.drop_duplicates(subset=['Rf', 'wee', 'k'], keep='last')
        have_pairs = {
            _grid_key_1d(rf, wee, rf_decimals=3, other_decimals=3)
            for rf, wee in zip(df['Rf'], df['wee'])
        }
    else:
        have_pairs = set()

    missing_pairs = sorted(target_pairs - have_pairs)

    if missing_pairs:
        print(f"Need {len(missing_pairs)} missing (Rf, wee) cases at k={target_k:.7f}.")

        missing_sim_tasks = []
        for Rf_key, wee_key in missing_pairs:
            fn = f'raw_data_1d_Rf_{Rf_key:.6f}_k_{target_k:.7f}_wee_{wee_key:.6f}.npz'
            p = os.path.join(raw_data_dir, fn)
            if not os.path.exists(p):
                missing_sim_tasks.append((params, Rf_key, wee_key, target_k, raw_data_dir, plot_dir))

        if missing_sim_tasks:
            print(f"Running {len(missing_sim_tasks)} missing simulations...")
            max_workers = mp.cpu_count() if n_jobs == -1 else n_jobs
            with concurrent.futures.ProcessPoolExecutor(max_workers=max_workers) as executor:
                futures = [executor.submit(simulate_plot_and_save_task_1d, task) for task in missing_sim_tasks]
                for future in tqdm(concurrent.futures.as_completed(futures), total=len(futures), desc="Simulating 1D"):
                    try:
                        future.result()
                    except Exception as e:
                        print(f"\nSimulation worker crashed: {e}")

        missing_npz_paths = []
        for Rf_key, wee_key in missing_pairs:
            fn = f'raw_data_1d_Rf_{Rf_key:.6f}_k_{target_k:.7f}_wee_{wee_key:.6f}.npz'
            p = os.path.join(raw_data_dir, fn)
            if os.path.exists(p):
                missing_npz_paths.append(p)

        if missing_npz_paths:
            print(f"Analyzing {len(missing_npz_paths)} missing datasets...")
            max_workers = mp.cpu_count() if n_jobs == -1 else n_jobs
            tasks = [(path, params, plot_dir, save_diagnostic_plot, plot_range) for path in missing_npz_paths]
            results = []

            with concurrent.futures.ProcessPoolExecutor(max_workers=max_workers) as executor:
                futures = [executor.submit(analyze_saved_data_task_1d, task) for task in tasks]
                for future in tqdm(concurrent.futures.as_completed(futures), total=len(futures), desc="Analyzing 1D"):
                    try:
                        res = future.result()
                        if res is not None:
                            results.append(res)
                    except Exception as e:
                        print(f"\nAnalysis worker crashed: {e}")

            if results:
                df_new = pd.DataFrame(results).sort_values(['Rf', 'wee'])
                if save_summary_csv:
                    summary_path = os.path.join(plot_dir, f'summary_1d_fixed_k_{target_k:.7f}_grid.csv')
                    df_new.to_csv(summary_path, index=False)
                    print(f"Saved merged summary to {summary_path}")

                df = pd.concat([df, df_new], ignore_index=True) if not df.empty else df_new
                df = df.drop_duplicates(subset=['Rf', 'wee', 'k'], keep='last')

    if df.empty:
        raise ValueError("No valid 1D summary data available after checking CSV, NPZ, and simulations.")

    df = df[np.isclose(df['k'], target_k, rtol=0, atol=atol_k)].copy()

    keep_mask = [
        _grid_key_1d(rf, wee, rf_decimals=3, other_decimals=3) in target_pairs
        for rf, wee in zip(df['Rf'], df['wee'])
    ]
    df = df[keep_mask].copy()

    if df.empty:
        raise ValueError("No rows matched the requested (Rf, wee) grid at the target k.")

    return df.sort_values(['Rf', 'wee']).reset_index(drop=True)


def _run_and_save_worker(task_args):
    """
    Top-level worker function to prevent multiprocessing pickling errors.
    """
    Rf_val, params, ff_scale, wee, k, raw_data_dir = task_args
    
    simulator = CANNSimulator(params)
    res = simulator.run_simulation(float(Rf_val), 0, float(ff_scale), float(wee), k, "normal")
    
    save_fn = f'raw_data_1d_Rf_{float(Rf_val):.6f}_k_{float(k):.7f}_wee_{round(float(wee), 3):.6f}.npz'
    save_path = os.path.join(raw_data_dir, save_fn)
    
    time_array = res.get("time", np.array([]))
    delta_z_values = res.get("delta_z", np.array([]))
    E_stim_array = res.get("E_stim", np.array([]))
    
    safe_save_npz(
        save_path,
        time=time_array,
        z_list=E_stim_array,
        Ue_list=res["E_bump_height"],
        delta_z=delta_z_values,
        E_FR=res["E_FR"],
        Rf=np.array(Rf_val),
        k=np.array(k),
        wee=np.array(wee)
    )
    
    return save_path


def find_left_superlinear_region(
    x,
    y,
    smooth_window=5,
    polyorder=2,
    slope_frac=0.10,
    min_points=3,
):
    """Find the early superlinear region of y(x)."""
    x = np.asarray(x, dtype=float)
    y = np.asarray(y, dtype=float)

    finite = np.isfinite(x) & np.isfinite(y)
    x = x[finite]
    y = y[finite]

    if len(x) < min_points:
        return None, None, {}

    order = np.argsort(x)
    x = x[order]
    y = y[order]

    x_unique, inverse = np.unique(x, return_inverse=True)
    y_unique = np.array([np.mean(y[inverse == i]) for i in range(len(x_unique))])

    x = x_unique
    y = y_unique
    n = len(x)

    if n < min_points:
        return None, None, {}

    smooth_window = int(smooth_window)
    if smooth_window % 2 == 0:
        smooth_window += 1

    smooth_window = min(smooth_window, n if n % 2 == 1 else n - 1)

    if smooth_window >= 3 and smooth_window > polyorder:
        y_smooth = savgol_filter(y, window_length=smooth_window, polyorder=polyorder)
    else:
        y_smooth = y.copy()

    dx = np.diff(x)
    dy = np.diff(y_smooth)

    valid_dx = dx > 1e-12
    if np.sum(valid_dx) < min_points - 1:
        return None, None, {}

    x_mid = 0.5 * (x[:-1] + x[1:])
    slope = np.full_like(dx, np.nan, dtype=float)
    slope[valid_dx] = dy[valid_dx] / dx[valid_dx]

    if not np.any(np.isfinite(slope)):
        return None, None, {}

    max_slope = np.nanmax(slope)

    if not np.isfinite(max_slope) or max_slope <= 0:
        return None, None, {
            "x": x,
            "y_smooth": y_smooth,
            "x_mid": x_mid,
            "slope": slope,
        }

    peak_interval = int(np.nanargmax(slope))
    end_idx = peak_interval + 1

    active = slope[: peak_interval + 1] > slope_frac * max_slope

    if np.any(active):
        start_interval = int(np.argmax(active))
        start_idx = start_interval
    else:
        start_idx = 0

    if end_idx - start_idx + 1 < min_points:
        start_idx = max(0, end_idx - min_points + 1)

    start_x = x[start_idx]
    end_x = x[end_idx]

    info = {
        "x": x,
        "y": y,
        "y_smooth": y_smooth,
        "x_mid": x_mid,
        "slope": slope,
        "peak_interval": peak_interval,
        "start_idx": start_idx,
        "end_idx": end_idx,
    }

    return start_x, end_x, info


def plot_R_U2(params, wee, recording_start=50, dt=0.1, Rflist=None, max_workers=None):
    params_new = params.copy()
    params_new['simulation_time'] = 500

    simulator2 = CANNSimulator(params_new)
    critical_weight = simulator2.params['critical_weight']
    
    a = params_new['gaussian_width_exc']
    rho = 0.5
    k = params_new["normalization_k"]
    ff_scale = params_new.get('feedforward_scale', 0.71648)
    
    raw_data_dir = 'output_net/raw_npz/1dUR/'
    plot_dir = 'output_net/plots/1d/'
    os.makedirs(raw_data_dir, exist_ok=True)
    os.makedirs(plot_dir, exist_ok=True)
    
    Ue_steady_list = []
    Relist_steady_list = []
    
    missing_sim_tasks = []
    for Rf in Rflist:
        fn = f'raw_data_1d_Rf_{float(Rf):.6f}_k_{float(k):.7f}_wee_{round(float(wee), 3):.6f}.npz'
        p = os.path.join(raw_data_dir, fn)
        if not os.path.exists(p):
            missing_sim_tasks.append((Rf, params_new, ff_scale, wee, k, raw_data_dir))

    if missing_sim_tasks:
        print(f"Running {len(missing_sim_tasks)} missing simulations...")
        
        with concurrent.futures.ProcessPoolExecutor(max_workers=max_workers) as executor:
            future_to_rf = {
                executor.submit(_run_and_save_worker, task): task[0] 
                for task in missing_sim_tasks
            }
            
            for future in tqdm(concurrent.futures.as_completed(future_to_rf), total=len(future_to_rf), desc="Simulating 1D"):
                Rf_val = future_to_rf[future]
                try:
                    future.result()
                except Exception as e:
                    print(f"\nSimulation worker crashed for Rf={Rf_val}: {e}")
                    print(traceback.format_exc())

    successful_Rflist = []
    
    for Rf in Rflist:
        fn = f'raw_data_1d_Rf_{float(Rf):.6f}_k_{float(k):.7f}_wee_{round(float(wee), 3):.6f}.npz'
        p = os.path.join(raw_data_dir, fn)
        
        if os.path.exists(p):
            try:
                data = np.load(p)
                
                Ue_timeseries = data['Ue_list'] 
                E_FR = data['E_FR']
                
                Relist_timeseries = np.sum(E_FR, axis=0) / (np.sqrt(np.pi) * 2 * a * float(rho))
                
                start_index = int(recording_start / dt)
                Ue_steady = np.average(Ue_timeseries[start_index:])
                Relist_steady = np.average(Relist_timeseries[start_index:])

                Ue_steady_list.append(Ue_steady)
                Relist_steady_list.append(Relist_steady)
                successful_Rflist.append(Rf)
            except Exception as e:
                print(f"Failed to load or process data for Rf={Rf}: {e}")
        else:
            print(f"Warning: File {p} still missing. Skipping Rf={Rf}.")

    Rflist = np.array(successful_Rflist)

    if len(Rflist) == 0:
        print("No successful data to plot. Exiting function.")
        return

    pd.DataFrame({
        'Rf': Rflist,
        'wee': wee,
        'Ue_steady': Ue_steady_list,
        'Re_steady': Relist_steady_list
    }).to_csv(os.path.join(plot_dir, f'steady_state_Rf_Ue_Re_wee_{wee}_Rf_test.csv'), index=False)

    Ue_arr = np.asarray(Ue_steady_list, dtype=float)
    Re_arr = np.asarray(Relist_steady_list, dtype=float)
    Rflist = np.asarray(Rflist, dtype=float)

    start_Ue, end_Ue, info = find_left_superlinear_region(
        x=Ue_arr,
        y=Re_arr,
        smooth_window=5,
        polyorder=2,
        slope_frac=0.10,
        min_points=3,
    )

    if start_Ue is not None and end_Ue is not None:
        superlinear_mask = (Ue_arr >= start_Ue) & (Ue_arr <= end_Ue)
        linear_sublinear_mask = ~superlinear_mask

        start_rf = np.min(Rflist[superlinear_mask])
        end_rf = np.max(Rflist[superlinear_mask])

        print(
            f"superlinear region found: "
            f"U_E = {start_Ue:.4f} to {end_Ue:.4f}, "
            f"R_f = {start_rf:.4f} to {end_rf:.4f}"
        )
    else:
        superlinear_mask = np.zeros_like(Ue_arr, dtype=bool)
        linear_sublinear_mask = ~superlinear_mask
        start_rf = None
        end_rf = None

        print("No clear superlinear region found.")
    
    # First Plot
    plt.figure(figsize=(12, 7))
    plt.scatter(Ue_steady_list, Relist_steady_list, c=Rflist, cmap='cividis', s=50)
    
    cbar = plt.colorbar()
    cbar.set_label(r'Input Strength ($R_f$)')
    if start_rf is not None and end_rf is not None:
        plt.axvspan(
            start_Ue,
            end_Ue,
            alpha=0.15,
            label="Detected superlinear region"
        )
        plt.title(f'superlinear Increase of R_E vs. U_E ({start_rf:.2f}-{end_rf:.2f}) for wee={wee/critical_weight:.2f}')
    else:
        plt.title(f'R_E vs. U_E for wee={wee/critical_weight:.2f}')
    plt.xlabel(r'Bump Amplitude ($U_E$)')
    plt.ylabel(r'Mean Firing Rate ($R_E$)')
    plt.legend()
    plt.savefig(os.path.join(plot_dir, f'steady_state_Rf_Ue_Re_wee_{wee/critical_weight:.2f}.svg'), dpi=300, bbox_inches='tight')
    
    # Second Plot
    plt.figure(figsize=(10, 6))
    plt.plot(Rflist, Ue_steady_list, marker='o', linestyle='-', label='$U_E$ (Bump Amplitude)')
    plt.plot(Rflist, Relist_steady_list, marker='x', linestyle='--', label='$R_E$ (Mean Firing Rate)')
    if start_rf is not None and end_rf is not None:
        plt.title(f'Steady-State Response vs. Input Strength ($R_f$) ({start_rf:.2f}-{end_rf:.2f}) for wee={wee/critical_weight:.2f}')
    else:
        plt.title(f'Steady-State Response vs. Input Strength ($R_f$) for wee={wee/critical_weight:.2f}')
    plt.xlabel(r'Input Strength ($R_f$)')
    plt.ylabel('Steady-State Value')
    plt.legend()
    plt.grid(True)
    plt.savefig(os.path.join(plot_dir, f'steady_state_Rf_vs_steady_state_wee_{wee/critical_weight:.2f}.svg'), dpi=300, bbox_inches='tight')
    
    # Third Plot
    plt.figure(figsize=(10, 6))
    plt.scatter(Ue_steady_list, Relist_steady_list, c=Rflist, cmap='cividis', s=50)
    plt.colorbar(label='Input Rf')
    if start_rf is not None and end_rf is not None:
        plt.title(f'Steady-State R_E vs U_E with Varying Input Rf_{start_rf:.2f}-{end_rf:.2f} for wee={wee/critical_weight:.2f}')
    else:
        plt.title(f'Steady-State R_E vs U_E with Varying Input Rf for wee={wee/critical_weight:.2f}')
    plt.xlabel('$U_E$')
    plt.ylabel('$R_E$')
    plt.grid(True)
    plt.savefig(os.path.join(plot_dir, f'steady_state_Ue_Re_scatter_wee_{wee/critical_weight:.2f}.svg'), dpi=300, bbox_inches='tight')
    
    plt.close('all') 
    print("Done! Plots saved successfully.")


def _format_slope_tick(x, pos):
    """Keep integers clean, allow 2 decimal places for fractions."""
    if abs(x - round(x)) < 1e-5:
        return f"{int(round(x))}"
    return f"{x:.2f}"


def _format_bof_tick(x, pos=None):
    """Force integers for large AIC values, allow decimals for tiny deltas, use scientific for extremes."""
    if abs(x) >= 10000:
        return f"{x:.1e}"
    if abs(x) >= 10 or abs(x - round(x)) < 1e-5:
        return f"{int(round(x))}"
    return f"{x:.2f}"


def _get_symlog_levels(vmin, vmax, linthresh=10.0, num=100):
    """Generates smoothly distributed log-levels for contourf so transitions aren't blocky."""
    if vmin >= vmax:
        return np.linspace(vmin - 1, vmax + 1, num)
        
    def forward(x):
        return np.sign(x) * np.log10(1 + np.abs(x) / linthresh)
        
    def inverse(x):
        return np.sign(x) * linthresh * (10**np.abs(x) - 1)
        
    t_min = forward(vmin)
    t_max = forward(vmax)
    t_levels = np.linspace(t_min, t_max, num)
    return inverse(t_levels)


def _get_asymmetric_cmap(vmin, vmax, center=0.0, cmap_name='RdBu_r', norm=None):
    """
    Dynamically rebuilds a divergent colormap so that the center color (white) 
    is strictly pinned to 'center', even if vmin and vmax are wildly asymmetric.
    """
    if norm is None:
        norm = mcolors.Normalize(vmin=vmin, vmax=vmax)
    
    zero_pt = norm(center)
    
    if zero_pt <= 0.0 or zero_pt >= 1.0:
        return plt.get_cmap(cmap_name)
        
    base_cmap = plt.get_cmap(cmap_name)
    colors = []
    
    for x in np.linspace(0, 1, 256):
        if x < zero_pt:
            c_idx = 0.5 * (x / zero_pt)
        else:
            c_idx = 0.5 + 0.5 * ((x - zero_pt) / (1.0 - zero_pt))
        colors.append(base_cmap(c_idx))
        
    return mcolors.LinearSegmentedColormap.from_list(f'asym_{cmap_name}', colors)


def _smooth_grid(x, y, z, zoom=5):
    """
    Uses strict Linear interpolation.
    Prevents the 'roller coaster' overshoot on extreme cliffs, 
    guaranteeing the smoothed grid NEVER exceeds raw max/min values.
    """
    interp = RegularGridInterpolator((y, x), z, method='linear')
    
    x_smooth = np.linspace(x.min(), x.max(), len(x) * zoom)
    y_smooth = np.linspace(y.min(), y.max(), len(y) * zoom)
    X_smooth, Y_smooth = np.meshgrid(x_smooth, y_smooth)
    
    pts = np.array([Y_smooth.ravel(), X_smooth.ravel()]).T
    z_smooth = interp(pts).reshape(X_smooth.shape)
    
    return X_smooth, Y_smooth, z_smooth


def plot_tail_slope_vs_wee_rf(
    params,
    target_k,
    Rf_list,
    wee_list,
    raw_data_dir='output_net/raw_npz/1d/',
    plot_dir='output_net/plots/1d/',
    n_jobs=-1,
    value_col='slope_evt_dz',
    force_reanalyze=False,
    save=True,
    save_diagnostic_plot=False,
    show_wee_in_wc=True,
):
    df = _collect_or_build_summary_1d_fixed_k(
        params=params, target_k=target_k, Rf_list=Rf_list, wee_list=wee_list,
        raw_data_dir=raw_data_dir, plot_dir=plot_dir, n_jobs=n_jobs,
        force_reanalyze=force_reanalyze, save_diagnostic_plot=save_diagnostic_plot,
        save_summary_csv=True,
    )

    if value_col not in df.columns:
        raise ValueError(f"{value_col} not found in summary dataframe.")

    df = df.dropna(subset=['Rf', 'wee', value_col]).copy()
    if df.empty:
        raise ValueError(f"No valid rows available for {value_col}.")

    pivot = df.pivot_table(index='Rf', columns='wee', values=value_col, aggfunc='mean').sort_index()
    
    pivot = pivot.interpolate(method='linear', axis=1, limit_direction='both')
    pivot = pivot.interpolate(method='linear', axis=0, limit_direction='both')
    pivot = pivot.bfill(axis=0).ffill(axis=0).bfill(axis=1).ffill(axis=1)
    
    y_vals = pivot.index.values.astype(float)
    x_vals = pivot.columns.values.astype(float)
    
    if show_wee_in_wc:
        sim_temp = CANNSimulator(params)
        wc = float(sim_temp.params['critical_weight'])
        x_vals = x_vals / wc
        x_label = r'$w_{ee}\;(\times w_c)$'
    else:
        x_label = 'wee'
        
    Z = pivot.values
    
    X_smooth, Y_smooth, Z_smooth = _smooth_grid(x_vals, y_vals, Z, zoom=8)
    
    fig, ax = plt.subplots(figsize=(8, 6), constrained_layout=True)
    
    z_min = np.nanpercentile(Z, 5) 
    z_max = np.nanpercentile(Z, 95)
    
    vmin = min(z_min, -2.0 - 1e-5)
    vmax = max(z_max, -2.0 + 1e-5)
    
    levels_smooth = np.unique(np.linspace(vmin, vmax, 100))
    
    cf = ax.contourf(X_smooth, Y_smooth, Z_smooth, levels=levels_smooth, cmap='YlGnBu', extend='both')
    cf.set_edgecolor("face") 
    
    step = 2.0 if (vmax - vmin) > 15 else 1.0
    cbar_ticks = np.arange(np.ceil(vmin), np.floor(vmax) + 0.1, step)
    if -2.0 not in cbar_ticks and vmin <= -2.0 <= vmax:
        cbar_ticks = np.sort(np.append(cbar_ticks, -2.0))
        
    cbar = fig.colorbar(cf, ax=ax, ticks=cbar_ticks)
    cbar.ax.yaxis.set_major_formatter(ticker.FuncFormatter(_format_slope_tick))
    
    line_levels = np.unique(np.linspace(np.ceil(vmin), np.floor(vmax), 8))
    
    if len(line_levels) > 0:
        c_lines = ax.contour(X_smooth, Y_smooth, Z_smooth, levels=line_levels, colors='black', linewidths=1, alpha=0.5)
        ax.clabel(c_lines, inline=True, fontsize=10, fmt='%.1f')

    if vmin <= -2.0 <= vmax:
        c_line_target = ax.contour(X_smooth, Y_smooth, Z_smooth, levels=[-2.0], colors='black', linewidths=2)
        ax.clabel(c_line_target, inline=True, fontsize=12, fmt='-2.0')
        
    ax.set_title(f'Step-size tail metric across wee and Rf: {value_col} (k={target_k:.7f})')
    ax.set_xlabel(x_label)
    ax.set_ylabel('Rf')

    if save:
        suffix = '_wc' if show_wee_in_wc else ''
        out = os.path.join(plot_dir, f'{value_col}_vs_wee_rf_k_{target_k:.7f}{suffix}.svg')
        fig.savefig(out, dpi=150, bbox_inches='tight')
        print(f"Saved: {out}")

    return fig, df, pivot


def plot_bof_vs_wee_rf(
    params,
    target_k,
    Rf_list,
    wee_list,
    raw_data_dir='output_net/raw_npz/1d/',
    plot_dir='output_net/plots/1d/',
    n_jobs=-1,
    metric='delta_aic',
    force_reanalyze=False,
    save=True,
    save_diagnostic_plot=False,
    show_wee_in_wc=True,
):
    df = _collect_or_build_summary_1d_fixed_k(
        params=params, target_k=target_k, Rf_list=Rf_list, wee_list=wee_list,
        raw_data_dir=raw_data_dir, plot_dir=plot_dir, n_jobs=n_jobs,
        force_reanalyze=force_reanalyze, save_diagnostic_plot=save_diagnostic_plot,
        save_summary_csv=True,
    )

    req = ['Rf', 'wee', 'aic_U_gamma', 'aic_U_gauss', 'aic_dz_t', 'aic_dz_gauss']
    df = df.dropna(subset=req).copy()
    if df.empty:
        raise ValueError("No valid rows available for best-of-fit plotting.")

    df['delta_aic_U'] = df['aic_U_gamma'] - df['aic_U_gauss']
    df['delta_aic_dz'] =  df['aic_dz_t'] - df['aic_dz_gauss']
    df['winner_U'] = (df['delta_aic_U'] < 0).astype(float)
    df['winner_dz'] = (df['delta_aic_dz'] < 0).astype(float)

    if metric == 'delta_aic':
        valU, valdz = 'delta_aic_U', 'delta_aic_dz'
        titleU = r'U fit: $\Delta$AIC = AIC(Gamma) - AIC(Gaussian)'
        titledz = r'$\Delta z$ fit: $\Delta$AIC = AIC(Student-t) - AIC(Gauss)'
        
        base_blues = plt.get_cmap('Blues_r')
        cmap_dz = mcolors.LinearSegmentedColormap.from_list('trunc_Blues_r', base_blues(np.linspace(0, 0.85, 256)))
        
    elif metric == 'winner':
        valU, valdz = 'winner_U', 'winner_dz'
        titleU = 'U fit winner (1 = Gamma better)'
        titledz = 'Δz fit winner (1 = Student-t better)'
        cmap_U = 'cividis'
        base_blues = plt.get_cmap('Blues_r')
        cmap_dz = mcolors.LinearSegmentedColormap.from_list('trunc_Blues_r', base_blues(np.linspace(0, 0.85, 256)))
    else:
        raise ValueError("metric must be 'delta_aic' or 'winner'")

    pivot_U = df.pivot_table(index='Rf', columns='wee', values=valU, aggfunc='mean').sort_index()
    pivot_dz = df.pivot_table(index='Rf', columns='wee', values=valdz, aggfunc='mean').sort_index()

    for pivot in [pivot_U, pivot_dz]:
        pivot.interpolate(method='linear', axis=1, limit_direction='both', inplace=True)
        pivot.interpolate(method='linear', axis=0, limit_direction='both', inplace=True)
        pivot.bfill(axis=0).ffill(axis=0).bfill(axis=1).ffill(axis=1).fillna(0, inplace=True)

    y_vals = pivot_U.index.values.astype(float)
    x_vals = pivot_U.columns.values.astype(float)
    
    if show_wee_in_wc:
        sim_temp = CANNSimulator(params)
        wc = float(sim_temp.params['critical_weight'])
        x_vals = x_vals / wc
        x_label = r'$w_{ee}\;(\times w_c)$'
    else:
        x_label = 'wee'

    ZU = pivot_U.values
    Zdz = pivot_dz.values

    X_smooth, Y_smooth, ZU_smooth = _smooth_grid(x_vals, y_vals, ZU, zoom=8)
    _, _, Zdz_smooth = _smooth_grid(x_vals, y_vals, Zdz, zoom=8)

    fig, axes = plt.subplots(1, 2, figsize=(15, 6), constrained_layout=True)
    
    if metric == 'delta_aic':
        linthresh = 10.0 
        
        vmin_U = min(np.nanpercentile(ZU, 2), -linthresh)
        vmax_U = max(np.nanpercentile(ZU, 98), linthresh)
        
        norm_U = mcolors.SymLogNorm(linthresh=linthresh, vmin=vmin_U, vmax=vmax_U, base=10)
        cmap_U = _get_asymmetric_cmap(vmin_U, vmax_U, center=0.0, cmap_name='RdBu_r', norm=norm_U)
        
        levels_U = np.unique(_get_symlog_levels(vmin_U, vmax_U, linthresh=linthresh, num=100))
        locator_U = ticker.SymmetricalLogLocator(linthresh=linthresh, base=10)
        ticks_U = locator_U.tick_values(vmin_U, vmax_U)
        
        vmin_dz = np.nanpercentile(Zdz, 2)
        vmax_dz = np.nanmax(Zdz) 
        if vmax_dz <= vmin_dz: 
            vmax_dz = vmin_dz + 1.0 
        
        norm_dz = None 
        levels_dz = np.unique(np.linspace(vmin_dz, vmax_dz, 100))
        ticks_dz = np.unique(np.linspace(vmin_dz, vmax_dz, 8))
        
    else:
        levels_U = np.unique(np.linspace(np.nanpercentile(ZU, 5), np.nanpercentile(ZU, 95), 100))
        levels_dz = np.unique(np.linspace(np.nanpercentile(Zdz, 2), np.nanmax(Zdz), 100))
        norm_U, norm_dz = None, None
        ticks_U, ticks_dz = None, None
        cmap_U = 'cividis'

    cf_U = axes[0].contourf(X_smooth, Y_smooth, ZU_smooth, levels=levels_U, norm=norm_U, cmap=cmap_U, extend='both')
    cf_U.set_edgecolor("face")
        
    cbar_U = fig.colorbar(cf_U, ax=axes[0], ticks=ticks_U)
    if metric == 'delta_aic':
        cbar_U.ax.yaxis.set_major_formatter(ticker.FuncFormatter(_format_bof_tick))
        line_lvls_U = np.unique(_get_symlog_levels(np.nanmin(ZU_smooth), np.nanmax(ZU_smooth), linthresh=linthresh, num=12))
    else:
        line_lvls_U = np.unique(np.linspace(np.nanmin(ZU_smooth), np.nanmax(ZU_smooth), 9))
        
    if len(line_lvls_U) > 0:
        cl_U = axes[0].contour(X_smooth, Y_smooth, ZU_smooth, levels=line_lvls_U, colors='black', linewidths=1, alpha=0.5)
        axes[0].clabel(cl_U, inline=True, fontsize=20, fmt=_format_bof_tick if metric == 'delta_aic' else '%.1f')
    
    if np.nanmin(ZU) <= 0.0 <= np.nanmax(ZU):
        cl_U_0 = axes[0].contour(X_smooth, Y_smooth, ZU_smooth, levels=[0.0], colors='black', linewidths=2)
        axes[0].clabel(cl_U_0, inline=True, fontsize=12, fmt='0')

    axes[0].set_title(titleU)
    axes[0].set_xlabel(x_label)
    axes[0].set_ylabel('Rf')

    cf_dz = axes[1].contourf(X_smooth, Y_smooth, Zdz_smooth, levels=levels_dz, norm=norm_dz, cmap=cmap_dz, extend='both')
    cf_dz.set_edgecolor("face")
        
    cbar_dz = fig.colorbar(cf_dz, ax=axes[1], ticks=ticks_dz)
    if metric == 'delta_aic':
        cbar_dz.ax.yaxis.set_major_formatter(ticker.FuncFormatter(_format_bof_tick))
        line_lvls_dz = np.unique(np.linspace(np.nanmin(Zdz_smooth), np.nanmax(Zdz_smooth), 8))
    else:
        line_lvls_dz = np.unique(np.linspace(np.nanmin(Zdz_smooth), np.nanmax(Zdz_smooth), 9))
        
    if len(line_lvls_dz) > 0:
        cl_dz = axes[1].contour(X_smooth, Y_smooth, Zdz_smooth, levels=line_lvls_dz, colors='black', linewidths=1, alpha=0.5)
        axes[1].clabel(cl_dz, inline=True, fontsize=10, fmt=_format_bof_tick if metric == 'delta_aic' else '%.1f')

    axes[1].set_title(titledz)
    axes[1].set_xlabel(x_label)
    axes[1].set_ylabel('Rf')

    fig.suptitle(f'Best-of-fit comparison across wee and Rf (k={target_k:.7f})', fontsize=16)

    if save:
        suffix = '_wc' if show_wee_in_wc else ''
        out = os.path.join(plot_dir, f'bof_vs_wee_rf_{metric}_k_{target_k:.7f}{suffix}.svg')
        fig.savefig(out, dpi=150, bbox_inches='tight')
        print(f"Saved: {out}")
        
    return fig, df, pivot_U, pivot_dz


def plot_one_simulation_Lan(params, Rf=10, test_eq="normal", tstart=0, tend=100):
    """
    Plot one simulation for Langevin sampling

    Args:
    - params: dictionary containing simulation parameters
    - Rf: float, input intensity for Langevin sampling
    - test_eq: str, type of input used for initialization
    - tstart: int, start time for plotting
    - tend: int, end time for plotting
    """
    params['simulation_time'] = tend
    simulator = CANNSimulator(params)
    results_Lan = simulator.run_simulation(
        Rf=Rf,
        Wei=0, 
        ff_scale=params['feedforward_scale'],
        Wee=simulator.params['recurrent_weight_e2e'],
        test_eq=test_eq
    )
    plot_results(results_Lan, "Langevin", tstart=tstart, tend=tend)


def plot_results(results, sampling, tstart=0, tend=100):
    """Visualization methods"""
    plot_firing_rates(results, sampling, tstart=tstart, tend=tend)    
    plot_bump_position_change(results, sampling, tstart=tstart, tend=tend)
    plot_bump_height(results, sampling, tstart=tstart, tend=tend)


def plot_bump_position_change(results, sampling, tstart=0, tend=100):
    """Plot bump position change over time"""
    t = np.arange(0, tend-tstart, results['params']['time_step'])
    figbp, axbp = plt.subplots(figsize=(2.5, 1.75))
    figbp.subplots_adjust(left=0.1, bottom=0.3, right=0.8, top=0.9, wspace=0.04, hspace=0.3)
    axbp.plot(t, results["delta_z"][(int(tstart/results['params']['time_step'])-1):(int(tend/results['params']['time_step'])-1)], label='Bump Position Change')
    axbp.set_xlim(0, tend-tstart)
    plt.title(f'Bump Position Change Over Time for {sampling} Sampling')
    plt.xlabel(r'Time /$\tau$') 
    plt.ylabel(r'Bump Position (degrees)')
    plt.savefig(f'one_simulation_bump_position_{sampling}.svg')
    plt.close()


def plot_firing_rates(results, sampling, tstart=0, tend=100):
    """Plot firing rates"""
    fig1, ax1 = plt.subplots(figsize=(2.5, 1.75))
    E_FR = results['E_FR'][:, int(tstart/results['params']['time_step']):int(tend/results['params']['time_step'])]
    plt.imshow(E_FR, aspect='auto', cmap=plt.get_cmap('Blues'),
        extent=[0, tend-tstart, -180, 180], origin='lower', vmin=0, vmax=100)
    ax1.set_yticks([-180, -120, -60, 0, 60, 120, 180])
    ax1.set_xlim(left=0)
    plt.colorbar(label='Firing Rate', shrink=0.5, ticks=[0, 50, 100])
    plt.title(f'Excitatory Population Firing Rate for {sampling} Sampling')
    plt.xlabel('Time Steps')
    plt.ylabel('Neuron Index')
    plt.savefig(f'one_simulation_FR_all_{sampling}.svg')
    plt.close()


def plot_bump_height(results, sampling, tstart=0, tend=100):
    """Plot bump height over time"""
    t = np.arange(0, tend-tstart, results['params']['time_step'])
    figbh, axbh = plt.subplots(figsize=(2.5, 1.75))
    figbh.subplots_adjust(left=0.1, bottom=0.3, right=0.8, top=0.9, wspace=0.04, hspace=0.3)
    axbh.plot(t, results['E_bump_height'][int(tstart/results['params']['time_step']):int(tend/results['params']['time_step'])], label='Bump Height')
    axbh.set_xlim(0, tend-tstart)
    plt.title(f'Bump Height Over Time for {sampling} Sampling')
    plt.xlabel(r'Time /$\tau$') 
    plt.ylabel(r'Bump Height')
    plt.savefig(f'one_simulation_bump_height_{sampling}.svg')
    plt.close()


def plot_dynamic_abs_stepsize_vs_invSqrtU(params, wee, Rf, recording_start=50, dt=0.1, n_bins=15):
    """
    Analyzes a SINGLE simulation run. Bins the instantaneous U_E(t) values,
    calculates the Mean Absolute Step Size <|Delta z(t)|> for each bin, 
    and plots it against 1/sqrt(U_E).
    """
    params_new = params.copy()
    tau = params_new['time_constant_exc']
    a_e = params_new['gaussian_width_exc']
    F = params_new.get('Fano_factor', 0.5)
    k = params_new.get("normalization_k", 0.0005)
    
    raw_data_dir = 'output_net/raw_npz/1dUR/'
    plot_dir = 'output_net/plots/1d/'
    os.makedirs(plot_dir, exist_ok=True)
    
    fn = f'raw_data_1d_Rf_{float(Rf):.6f}_k_{float(k):.7f}_wee_{round(float(wee), 3):.6f}.npz'
    p = os.path.join(raw_data_dir, fn)
    
    if not os.path.exists(p):
        print(f"Error: File {p} not found. Run the simulation for this Rf first.")
        return

    data = np.load(p)
    Ue_timeseries = data['Ue_list'] 
    delta_z = data['delta_z']
    
    start_index = int(recording_start / dt)
    Ue_valid = Ue_timeseries[start_index+1:]
    dz_valid = delta_z[start_index:]
    
    mask = np.isfinite(Ue_valid) & np.isfinite(dz_valid)
    Ue_valid = Ue_valid[mask]
    dz_valid = dz_valid[mask]

    u_min, u_max = np.percentile(Ue_valid, 1), np.percentile(Ue_valid, 99)
    bins = np.linspace(u_min, u_max, n_bins + 1)
    bin_centers = 0.5 * (bins[1:] + bins[:-1])
    
    indices = np.digitize(Ue_valid, bins)
    
    abs_dz_mean_list = []
    inv_sqrt_U_centers = []
    
    for i in range(1, len(bins)):
        dz_in_bin = dz_valid[indices == i]
        
        if len(dz_in_bin) > 50:
            abs_dz_mean_list.append(np.mean(np.abs(dz_in_bin)))
            inv_sqrt_U_centers.append(1.0 / np.sqrt(bin_centers[i-1]))
            
    inv_sqrt_U_arr = np.array(inv_sqrt_U_centers)
    abs_dz_mean_arr = np.array(abs_dz_mean_list)

    sigma_z_sq = (8.0 * a_e * F) / (3.0 * np.sqrt(3.0 * np.pi))
    theoretical_var_per_step = (sigma_z_sq * dt) / tau
    
    theory_slope = np.sqrt(theoretical_var_per_step) * np.sqrt(2.0 / np.pi)

    fig, ax = plt.subplots(figsize=(4.5, 4))
    
    ax.scatter(inv_sqrt_U_arr, abs_dz_mean_arr, color='#ff7f0e', s=60, label='Empirical (Binned)', zorder=3)
    
    x_grid = np.linspace(0, np.max(inv_sqrt_U_arr) * 1.05, 100)
    theory_line = theory_slope * x_grid
    
    ax.plot(x_grid, theory_line, color='black', linewidth=2, label='Theory (Gaussian)', zorder=2)

    ax.set_xlabel(r'$1 / \sqrt{U_E(t)}$ (Binned Instantaneous Inverse Sqrt Bump)')
    ax.set_ylabel(r'$\langle |\Delta z| \rangle$ (Mean Abs Step)')
    ax.set_title(f"Dynamic Step Size vs $1/\\sqrt{{U_E}}$\n(Rf={Rf})")
    ax.spines['top'].set_visible(False)
    ax.spines['right'].set_visible(False)
    ax.set_xlim(left=0)
    ax.set_ylim(bottom=0)
    
    ax.legend(frameon=False, loc='best', fontsize=10)
    
    plt.tight_layout()
    save_path = os.path.join(plot_dir, f'dynamic_abs_stepsize_vs_invSqrtU_Rf_{Rf}.svg')
    plt.savefig(save_path, dpi=300, transparent=True)
    plt.close()
    
    print(f"Saved dynamic absolute step size vs 1/sqrt(U) plot to {save_path}")


def plot_abs_stepsize_vs_inv_sqrt_U(params, wee, recording_start=50, dt=0.1, Rflist=None, max_workers=None):
    """
    Plots the Mean Absolute Step Size <|Delta z|> versus 1/sqrt(U_E).
    This provides a highly intuitive linear plot verifying the state-dependent noise.
    """
    params_new = params.copy()
    params_new['simulation_time'] = 500

    simulator2 = CANNSimulator(params_new)
    critical_weight = simulator2.params['critical_weight']
    tau = params_new['time_constant_exc']
    a_e = params_new['gaussian_width_exc']
    F = params_new.get('Fano_factor', 0.5)
    k = params_new.get("normalization_k", 0.0005)
    ff_scale = params_new.get('feedforward_scale', 0.71648)
    
    raw_data_dir = 'output_net/raw_npz/1dUR/'
    plot_dir = 'output_net/plots/1d/'
    os.makedirs(raw_data_dir, exist_ok=True)
    os.makedirs(plot_dir, exist_ok=True)
    
    abs_dz_mean_list = []
    inv_sqrt_Ue_list = []
    
    if Rflist is None:
        Rflist = np.linspace(0.01, 1.0, 20)
        
    missing_sim_tasks = []
    for Rf in Rflist:
        fn = f'raw_data_1d_Rf_{float(Rf):.6f}_k_{float(k):.7f}_wee_{round(float(wee), 3):.6f}.npz'
        p = os.path.join(raw_data_dir, fn)
        if not os.path.exists(p):
            missing_sim_tasks.append((Rf, params_new, ff_scale, wee, k, raw_data_dir))

    if missing_sim_tasks:
        print(f"Running {len(missing_sim_tasks)} missing simulations for |dz| vs 1/sqrt(U)...")
        with concurrent.futures.ProcessPoolExecutor(max_workers=max_workers) as executor:
            future_to_rf = {
                executor.submit(_run_and_save_worker, task): task[0] 
                for task in missing_sim_tasks
            }
            for future in tqdm(concurrent.futures.as_completed(future_to_rf), total=len(future_to_rf), desc="Simulating 1D"):
                pass

    successful_Rflist = []
    for Rf in Rflist:
        fn = f'raw_data_1d_Rf_{float(Rf):.6f}_k_{float(k):.7f}_wee_{round(float(wee), 3):.6f}.npz'
        p = os.path.join(raw_data_dir, fn)
        
        if os.path.exists(p):
            try:
                data = np.load(p)
                Ue_timeseries = data['Ue_list'] 
                delta_z = data['delta_z']
                
                start_index = int(recording_start / dt)
                
                valid_dz = delta_z[start_index:]
                valid_dz = valid_dz[np.isfinite(valid_dz)]
                
                Ue_steady = np.average(Ue_timeseries[start_index:])
                
                mean_abs_dz = np.mean(np.abs(valid_dz))
                
                inv_sqrt_Ue_list.append(1.0 / np.sqrt(Ue_steady))
                abs_dz_mean_list.append(mean_abs_dz)
                successful_Rflist.append(Rf)
            except Exception as e:
                print(f"Failed to process data for Rf={Rf}: {e}")

    if not successful_Rflist:
        print("No successful data to plot.")
        return

    inv_sqrt_Ue_arr = np.array(inv_sqrt_Ue_list)
    abs_dz_mean_arr = np.array(abs_dz_mean_list)

    sigma_z_sq = (8.0 * a_e * F) / (3.0 * np.sqrt(3.0 * np.pi))
    theoretical_var_per_step = (sigma_z_sq * dt) / tau
    
    theoretical_slope = np.sqrt(theoretical_var_per_step) * np.sqrt(2.0 / np.pi)

    fig, ax = plt.subplots(figsize=(3.5, 3))
    
    ax.scatter(inv_sqrt_Ue_arr, abs_dz_mean_arr, color='#ff7f0e', s=35, label='Sim', zorder=3)
    
    x_grid = np.linspace(0, np.max(inv_sqrt_Ue_arr) * 1.05, 100)
    theory_line = theoretical_slope * x_grid
    ax.plot(x_grid, theory_line, color='black', linewidth=1.5, label='Theory', zorder=2)
    
    ax.set_xlabel(r'$1 / \sqrt{U_E}$')
    ax.set_ylabel(r'$\langle |\Delta z| \rangle$ (Mean Abs Step)')
    ax.spines['top'].set_visible(False)
    ax.spines['right'].set_visible(False)
    ax.set_xlim(left=0)
    ax.set_ylim(bottom=0)
    
    ax.legend(frameon=False, loc='best', fontsize=12, handlelength=1.5)
    
    plt.tight_layout()
    save_path = os.path.join(plot_dir, f'abs_stepsize_vs_invSqrtU_wee_{wee/critical_weight:.2f}.svg')
    plt.savefig(save_path, dpi=300, transparent=True)
    plt.close()
    
    print(f"Saved absolute step size plot to {save_path}")


# ==========================================
# EXECUTION BLOCK
# ==========================================
if __name__ == '__main__':
    params = {
        'time_constant_exc': 1.0,
        'position_max': 180.0,
        'position_min': -180.0,
        'gaussian_width_exc': 40.0,
        'gaussian_width_ES': 5.0,
        'num_neurons': 180,
        'simulation_time':  5000,
        'time_step': 0.1,
        'recording_start': 500,
        'Fano_factor': 0.5,
        'normalization_k': 0.0005,
        'inhibitory_gain': 10,
        'input_position': 20,
        'feedforward_scale': 0.71648,
        't_steady': 50,
        'initial_mean_eq': 0,
        'initial_var_eq': 30,
        'initial_scale_eq': 1e-1,
        'tail_quantile': 0.99,
        "decode_win_bins": 1
    }
    # Just comment out the other parts if you want a specific figure.
    simulator = CANNSimulator(params)
    Rf_list = np.linspace(0.01, 0.1, 5).tolist() + np.linspace(0.2, 1.0, 5).tolist()
 
    target_wee = 0.4 * simulator.params['critical_weight']
    wc = simulator.params['critical_weight']

    # Figure 2 C
    plot_abs_stepsize_vs_inv_sqrt_U(params, wee=target_wee, recording_start=50, dt=0.1, Rflist=Rf_list, max_workers=4)
    
    # Figure 2 EF
    Rf_list_diag = [0.01, 2.5]
    df_tail = _collect_or_build_summary_1d_fixed_k(
        params=params,
        target_k=params['normalization_k'],
        Rf_list=Rf_list_diag,
        wee_list=[target_wee],
        raw_data_dir='output_net/raw_npz/1d3/',
        plot_dir='output_net/plots/1d3/',
        n_jobs=2,
        force_reanalyze=True,
        save_diagnostic_plot=True,
        save_summary_csv=False,
    )
    
    weelist = [0.1*wc, 0.2*wc, 0.3*wc, 0.4*wc, 0.5*wc, 0.6*wc, 0.7*wc, 0.8*wc, 0.9*wc, 1.0*wc] 
    
    # Figure 2B upper panel
    plot_R_U2(params, target_wee, Rflist=Rf_list, recording_start=50, dt=0.1, max_workers=-1)
    
    # Figure 3: BOF and Tail Slope across Rf and wee
    fig1, df_bof, pivot_U, pivot_dz = plot_bof_vs_wee_rf(
        params=params,
        target_k=params['normalization_k'],
        Rf_list=Rf_list,
        wee_list=weelist,
        raw_data_dir='output_net/raw_npz/1d/',
        plot_dir='output_net/plots/1d/',
        n_jobs=2,
        metric='delta_aic',
        force_reanalyze=False,          
        save=True,
        save_diagnostic_plot=False,
        show_wee_in_wc=True
    )

    fig2, df_tail, pivot_tail = plot_tail_slope_vs_wee_rf(
        params=params,
        target_k=params['normalization_k'],
        Rf_list=Rf_list,
        wee_list=weelist,
        raw_data_dir='output_net/raw_npz/1d/',
        plot_dir='output_net/plots/1d/',
        n_jobs=2,
        value_col='slope_evt_dz',   
        force_reanalyze=False,      
        save=True,
        save_diagnostic_plot=False,
        show_wee_in_wc=True
    )

    plt.close(fig1)
    plt.close(fig2)    
    
    # Figure 2 G
    plot_kl_div_with_lan_net(
        params=params,
        Rf=0.1,
        wee=target_wee,
        k=params['normalization_k'],
        num_trials=1000,
        stride=1,
        mode='cumulative',
        plot_dir='output_net/plots/1d2/',
        save=True,
        seed=10
    )
```
