## Supplementary Codes for "Scale-mixture Langevin sampling in the subspaces of recurrent cortical circuit dynamics": cann_simulator.py

```
import sys
import pickle
import concurrent.futures
import multiprocessing as mp
from functools import partial
from collections import defaultdict

import numpy as np
import numpy.matlib
import matplotlib.pyplot as plt
from scipy.stats import multivariate_normal
from matplotlib.collections import LineCollection
from tqdm import tqdm


class CANNSimulator:
    """
    A simulator for Continuous Attractor Neural Network (CANN) models.

    This class provides methods to initialize network parameters, run simulations,
    and analyze the results of CANN models. It supports various configurations
    and allows for the calibration of feedforward weights.

    Attributes:
        params (dict): Dictionary of simulation parameters.
        conn_params (dict): Dictionary of connection parameters.
    """

    def __init__(self, params=None):
        """
        Initialize the CANNSimulator with given parameters.

        Parameters:
            params (dict, optional): Dictionary of parameters to override default values.
        """
        default_params = {
            'time_constant_exc': 1.0,
            'position_max': 180.0,
            'position_min': -180.0,
            'gaussian_width_exc': 40.0,
            'gaussian_width_ES': 20.0,
            'num_neurons': 180,
            'simulation_time': 100.0,
            'time_step': 0.01,
            'recording_start': 20,
            'Fano_factor': 0.05,
            'normalization_k': 0.0005,
            'inhibitory_gain': 10,
            'input_position': 0,
            'feedforward_scale': 0.58214787,
            't_steady': 20,
            'initial_mean_eq': 0,
            'initial_var_eq': 60,
            'initial_scale_eq': 1e-1
        }
        self.params = {**default_params, **(params or {})}
        self._compute_parameters()
        self.conn_params = None
        self.initialize_network()

    def _compute_parameters(self):
        """Initialize and compute all simulation parameters."""
        p = self.params

        p['time_constant_som'] = p['time_constant_exc']
        p['gaussian_width_SE'] = np.sqrt(p['gaussian_width_exc'] ** 2 - p['gaussian_width_ES'] ** 2)
        p['gaussian_width_SS'] = np.sqrt((p['gaussian_width_exc'] ** 2 + p['gaussian_width_ES'] ** 2) / 2)

        p['position_range'] = p['position_max'] - p['position_min']
        p['neuron_density'] = p['num_neurons'] / p['position_range']
        
        p['PrefStim'] = np.linspace(
            p['position_min'], p['position_max'], p['num_neurons'] + 1
        )[1:p['num_neurons'] + 1]
        
        p['max_inhibition'] = 1 / (12 * p['neuron_density'])
        
        numerator = 8 * np.sqrt(2 * np.pi) * p['normalization_k'] * p['gaussian_width_exc']
        p['critical_weight'] = np.sqrt(numerator / p['neuron_density'])

        p['recurrent_weight_e2e'] = 0.5 * p['critical_weight']
        p['recurrent_weight_e2i'] = 0.5 * p['critical_weight']
        
        denominator = 2 * np.sqrt(np.pi) * p['normalization_k'] * p['gaussian_width_exc']
        p['feedforward_intensity_scale'] = (p['critical_weight'] / denominator)

    def initialize_network(self):
        """Initialize network connectivity kernels."""
        param_dict = self.params
        PrefStim = param_dict["PrefStim"]
        a_SE = param_dict["gaussian_width_SE"]
        xrange = param_dict['position_range']
        aE = param_dict["gaussian_width_exc"]
        a_ES = param_dict['gaussian_width_ES']

        W_angle = np.angle(np.exp((1j * (PrefStim - PrefStim[0]) * (2 * np.pi / xrange)))) * xrange / (2 * np.pi)
        
        W_kerFtE = np.exp(-W_angle**2 / (2 * aE**2)) / (np.sqrt(2 * np.pi) * aE)    
        WE = np.fft.fft(W_kerFtE)
        
        W_kerFtES = np.exp(-W_angle**2 / (2 * a_ES**2)) / (np.sqrt(2 * np.pi) * a_ES)   
        WES = np.fft.fft(W_kerFtES)
        
        W_kerFtSE = np.exp(-W_angle**2 / (2 * a_SE**2)) / (np.sqrt(2 * np.pi) * a_SE) 
        WSE = np.fft.fft(W_kerFtSE)

        self.conn_params = {"E Kernel": WE, 'ES Kernel': WES, 'SE Kernel': WSE}

    def _temporal_ff(self, Rf, T, mu):
        """Calculate temporal feedforward input."""
        a = self.params["gaussian_width_exc"]
        xrange = self.params['position_range']
        PrefStim = self.params['PrefStim']
        dt = self.params['time_step']
        N = self.params['num_neurons']
        
        x1 = np.repeat(mu, N)    
        pos1 = np.subtract(x1, PrefStim)
        pos1 = np.angle(np.exp(1j * pos1 * (2 * np.pi) / xrange)) * xrange / (2 * np.pi)
        pos1 = np.expand_dims(pos1, axis=1)
        pos1 = np.matlib.repmat(pos1, 1, int(T / dt))
        Ipos1 = Rf * np.exp(-(pos1**2) / (4 * a**2))

        return Ipos1
    
    def run_simulation(self, Rf, Wei, ff_scale, Wee, k=None, test_eq="normal", noise_type="normal", uniF=0.1):
        """Main simulation function."""
        param_dict = self.params
        N = param_dict["num_neurons"]
        T = param_dict["simulation_time"]
        dt = param_dict["time_step"]
        a = param_dict["gaussian_width_exc"]
        aS = param_dict["gaussian_width_SS"]
        
        if k is None:
            k = param_dict["normalization_k"]
        else:
            param_dict["normalization_k"] = k
            
        gain = param_dict["inhibitory_gain"]
        rho = param_dict["neuron_density"]
        F = param_dict["Fano_factor"]
        mu = param_dict['input_position']
        t_steady = param_dict['t_steady']
        tauE = param_dict["time_constant_exc"]
        tauS = param_dict['time_constant_som']
        WE = self.conn_params['E Kernel']
        WES = self.conn_params['ES Kernel']
        WSE = self.conn_params['SE Kernel']
        Wie = param_dict["recurrent_weight_e2i"]

        U = np.zeros([N, int(T/dt)])
        V = np.zeros([N, int(T/dt)])
        Oc = np.zeros([N, int(T/dt)])
        Vc = np.zeros([N, int(T/dt)])

        if test_eq == 'eq':        
            initial_mean = param_dict['initial_mean_eq']
            initial_var = param_dict['initial_var_eq']
            shift = np.random.normal(initial_mean, initial_var)
            Iext1 = self._temporal_ff(Rf, t_steady, int(shift)/2*2)
            Iext2 = self._temporal_ff(Rf, T-t_steady, mu)
            Iext = np.concatenate((Iext1, Iext2), axis=1)
        elif test_eq == 'non-eq':
            initial_scale = param_dict['initial_scale_eq']
            small_input = initial_scale
            Iext1 = small_input * np.ones([N, int(t_steady/dt)])
            Iext2 = self._temporal_ff(Rf, T-t_steady, mu)
            Iext = np.concatenate((Iext1, Iext2), axis=1)
        else:
            Iext = self._temporal_ff(Rf, T, mu)

        for i in range(int(T/dt) - 1):
            noisevr = U[:, i].clip(min=0)
            
            O = Oc[:, i]
            Oin = Vc[:, i]
            
            FRE = np.real(np.fft.ifft(WE * np.fft.fft(O)))
            FRSE = np.real(np.fft.ifft(WSE * np.fft.fft(O)))
            FRI = np.real(np.fft.ifft(WES * np.fft.fft(Oin)))

            if noise_type == "normal":
                U[:, i+1] = ((1 - dt/tauE) * U[:, i]) + (Wee * FRE * dt/tauE) + (Wei * FRI * dt/tauE) + (ff_scale * Iext[:, i] * dt/tauE) + (np.sqrt(F * noisevr * dt/tauE) * np.random.normal(0, 1, N))
                V[:, i+1] = ((1 - dt/tauS) * V[:, i]) + (Wie * FRSE * (dt/tauS))

                U[:, i+1] = U[:, i+1].clip(min=0)
                V[:, i+1] = V[:, i+1].clip(min=0)
                Ocn = U[:, i+1].clip(min=0) ** 2
                Ocd = np.multiply(k * rho, np.sum(Ocn, axis=0))
                Oc[:, i+1] = Ocn / (Ocd + 1)
                
            elif noise_type == "uniform":
                U[:, i+1] = ((1 - dt/tauE) * U[:, i]) + (Wee * FRE * dt/tauE) + (Wei * FRI * dt/tauE) + (ff_scale * Iext[:, i] * dt/tauE) + (np.sqrt(F * np.sum(noisevr) * uniF * dt/tauE) * np.random.normal(0, 1, N))
                V[:, i+1] = ((1 - dt/tauS) * V[:, i]) + (Wie * FRSE * (dt/tauS))

                U[:, i+1] = U[:, i+1].clip(min=0)
                V[:, i+1] = V[:, i+1].clip(min=0)                
                Ocn = U[:, i+1].clip(min=0) ** 2
                Ocd = np.multiply(k * rho, np.sum(Ocn, axis=0))
                Oc[:, i+1] = Ocn / (Ocd + 1)   
                
            elif noise_type == 'free':
                U[:, i+1] = ((1 - dt/tauE) * U[:, i]) + (Wee * FRE * dt/tauE) + (Wei * FRI * dt/tauE) + (ff_scale * Iext[:, i] * dt/tauE)
                V[:, i+1] = ((1 - dt/tauS) * V[:, i]) + (Wie * FRSE * (dt/tauS))

                U[:, i+1] = U[:, i+1].clip(min=0)
                V[:, i+1] = V[:, i+1].clip(min=0)
                Ocn = U[:, i+1].clip(min=0) ** 2
                Ocd = np.multiply(k * rho, np.sum(Ocn, axis=0))
                Oc[:, i+1] = Ocn / (Ocd + 1)                
                
            elif noise_type == "pv":
                U[:, i+1] = ((1 - dt/tauE) * U[:, i]) + (Wee * FRE * dt/tauE) + (Wei * FRI * dt/tauE) + (ff_scale * Iext[:, i] * dt/tauE) + (np.sqrt(F * noisevr * dt/tauE) * np.random.normal(0, 1, N))
                V[:, i+1] = ((1 - dt/tauS) * V[:, i]) + (Wie * FRSE * (dt/tauS))

                U[:, i+1] = U[:, i+1].clip(min=0)
                V[:, i+1] = V[:, i+1].clip(min=0)
                Ocn = U[:, i+1].clip(min=0) ** 2
                Ocd = np.multiply(k * rho, np.sum(Ocn, axis=0))
                if Ocd < 0:
                    Ocd = 0
                Oc[:, i+1] = Ocn / (Ocd + 1 + np.sqrt(Ocd) * np.random.normal(0, 1))
            else:
                raise ValueError("Invalid noise_type. Choose from 'normal', 'uniform', or 'pv'.")                  
            
            Vc[:, i+1] = gain * V[:, i+1]
        
        Oc = Oc.clip(min=0)
        
        return {
            'E_stim': self._calculate_stimulus(Oc),
            'delta_z': self._calculate_stimulus_change(Oc),
            'I_stim': self._calculate_stimulus(Vc),
            'E_bump_height': np.sum(U, axis=0) / (np.sqrt(np.pi) * 2 * a * rho),
            'I_bump_height': np.sum(V, axis=0) / (np.sqrt(np.pi) * 2 * aS * rho),
            'E_FR': Oc,
            'I_FR': Vc,
            'Synaptic_Input': U,
            'I_Synaptic_Input': V,
            'params': self.params
        }
         
    def _calculate_stimulus(self, activity):
        """Calculate stimulus position from activity."""
        pref_factor = self.params['PrefStim'] * np.pi / 180
        if np.all(activity == 0):
            return np.zeros(activity.shape[1])
        e = np.exp(1j * pref_factor) @ activity
        return np.angle(e, deg=True) 

    def _calculate_stimulus_change(self, activity):
        """Calculate change in stimulus position from activity."""
        pref_factor = self.params['PrefStim'] * np.pi / 180
        e = np.exp(1j * pref_factor) @ activity
        magnitudes = np.abs(e)
        
        normalized_e = np.zeros_like(e, dtype=complex)
        non_zero_mask = magnitudes > 0
        normalized_e[non_zero_mask] = e[non_zero_mask] / magnitudes[non_zero_mask]
        
        z_t = normalized_e[:-1]
        z_t_plus_1 = normalized_e[1:]
        delta_z_complex = z_t_plus_1 * np.conj(z_t)
        
        return np.angle(delta_z_complex, deg=True)

    def _get_uniF(self, Rf, Wei, ff_scale, Wee, test_eq, num_trials=100):
        """Calculate the Fano factor across trials for uniformity testing."""
        spike_counts = []

        for i in range(num_trials):
            result = self.run_simulation(Rf, Wei, ff_scale, Wee, test_eq, noise_type="uniform", uniF=0.1)
            firing_rates = result['Oc'] 
            target_neuron_rates = firing_rates[np.argmax(np.mean(firing_rates, axis=1)), :]
            count = np.sum(target_neuron_rates[int(self.params['recording_start'] / self.params['time_step']):])
            spike_counts.append(count)
            
        mean_count = np.mean(spike_counts)
        var_count = np.var(spike_counts)
        
        if mean_count == 0:
            return 0 
            
        return var_count / mean_count


def save_simulation_data_pickle(data, filename):
    """Save simulation data locally to a pickle file."""
    with open(filename, 'wb') as f:
        pickle.dump(data, f)   


def load_simulation_data_pickle(filename):
    """Load previously saved simulation data."""
    with open(filename, 'rb') as f:
        return pickle.load(f)


def bump_position_ham(params, Rf=10):
    """Helper wrapper function to run a standard simulated experiment setup."""
    simulator = CANNSimulator(params)
    simulator.initialize_network()
    results = simulator.run_simulation(
        Rf=Rf, 
        Wei=-0.6 * simulator.params['critical_weight'], 
        ff_scale=1.624,
        Wee=simulator.params['recurrent_weight_e2e']
    )
    
    ZE = results['E_stim']
    ZS = results['I_stim']
    UE = results['E_bump_height']
    UI = results['I_bump_height']
    return ZE, ZS, UE, UI


def _run_processpool_tasks(worker_fn, tasks, max_workers=None, desc="Simulating"):
    """Run top-level worker tasks with ProcessPoolExecutor and collect non-None results."""
    results = []
    with concurrent.futures.ProcessPoolExecutor(max_workers=max_workers) as executor:
        futures = [executor.submit(worker_fn, task) for task in tasks]

        for future in tqdm(
            concurrent.futures.as_completed(futures),
            total=len(futures),
            desc=desc,
            file=sys.stdout,
        ):
            try:
                res = future.result()
                if res is not None:
                    results.append(res)
            except Exception as e:
                print(f"\n⚠️ A worker process crashed during simulation: {e}")
    return results


def _calibration_bump_position_task(task):
    """Top-level worker to prevent pickling issues with multiprocessing pools."""
    params = dict(task["params"])
    simulator = CANNSimulator(params)
    result = simulator.run_simulation(
        Rf=task["Rf"],
        Wei=task["Wei"],
        ff_scale=task["ff_scale"],
        Wee=task["Wee"],
        k=task["k"],
        test_eq=task["test_eq"],
    )
    return result["E_stim"]


def _bump_height_position_task(task):
    """Top-level worker to calculate both bump height and position."""
    params = dict(task["params"])
    simulator = CANNSimulator(params)
    result = simulator.run_simulation(
        Rf=task["Rf"],
        Wei=task["Wei"],
        ff_scale=task["ff_scale"],
        Wee=task["Wee"],
        test_eq=task["test_eq"],
    )
    return {"E_bump_height": result["E_bump_height"], "E_stim": result["E_stim"]}


def calibrate_ff_scale(simulator, Rf_list=np.linspace(1, 25, 23), Wei=0, n_trials=5000, max_workers=None):
    """
    Calibrate feedforward weight using precision analysis (Standalone ProcessPoolExecutor version).

    Call as:
        new_ff = calibrate_ff_scale(simulator, Rf_list=..., Wei=..., n_trials=..., max_workers=...)
    """
    params = dict(simulator.params)
    t_record = params["recording_start"]
    dt = params["time_step"]
    rho = params["neuron_density"]
    a = params["gaussian_width_exc"]

    tasks = [
        {
            "params": params,
            "Rf": Rf,
            "Wei": Wei,
            "ff_scale": params["feedforward_scale"],
            "Wee": params["recurrent_weight_e2e"],
            "k": params["normalization_k"],
            "test_eq": "normal",
        }
        for Rf in Rf_list
        for _ in range(n_trials)
    ]

    results = _run_processpool_tasks(
        _calibration_bump_position_task,
        tasks,
        max_workers=max_workers,
        desc=f"Running {n_trials} trials per Rf",
    )

    record_precision_inv = []
    for res in results:
        Se = np.asarray(res).flatten()
        Se = Se[int(t_record / dt):]
        record_precision_inv.append(1.0 / np.var(Se))

    mean_precision_inv = []
    for i, _ in enumerate(Rf_list):
        block = record_precision_inv[i * n_trials : (i + 1) * n_trials]
        mean_precision_inv.append(np.mean(block))

    slope = np.polyfit(Rf_list, mean_precision_inv, 1)[0]
    theoretical_slope = np.sqrt(2 * np.pi) * rho / a
    new_ff = theoretical_slope / slope
    
    simulator.params["feedforward_scale"] = new_ff
    params["feedforward_scale"] = new_ff

    updated_tasks = [
        {
            "params": params,
            "Rf": Rf,
            "Wei": Wei,
            "ff_scale": new_ff,
            "Wee": params["recurrent_weight_e2e"],
            "k": params["normalization_k"],
            "test_eq": "normal",
        }
        for Rf in Rf_list
        for _ in range(n_trials)
    ]

    new_results = _run_processpool_tasks(
        _calibration_bump_position_task,
        updated_tasks,
        max_workers=max_workers,
        desc="Recalculating precision with updated ff_scale",
    )

    record_precision_inv_new = []
    for res in new_results:
        Se = np.asarray(res).flatten()
        Se = Se[int(t_record / dt):]
        record_precision_inv_new.append(1.0 / np.var(Se))

    mean_precision_inv_new = []
    for i, _ in enumerate(Rf_list):
        block = record_precision_inv_new[i * n_trials : (i + 1) * n_trials]
        mean_precision_inv_new.append(np.mean(block))

    plt.figure(figsize=(10, 6))
    plt.plot(
        Rf_list,
        theoretical_slope * np.array(Rf_list),
        label="Theoretical Prediction",
    )
    plt.plot(
        Rf_list,
        mean_precision_inv_new,
        "o",
        label=f"Simulation (mean over {n_trials} trials)",
    )
    plt.xlabel("Feedforward Intensity (Rf)")
    plt.ylabel("1 / Variance (Precision)")
    plt.title("Precision vs Input Intensity")
    plt.legend()
    plt.savefig(f"precision_vs_Rf_wei{Wei}_ff{new_ff:.3f}.svg")
    plt.close()

    return new_ff
```
