## Supplementary Codes for "Scale-mixture Langevin sampling in the subspaces of recurrent cortical circuit dynamics": 2d.py

```
import os
import glob
import traceback
import concurrent.futures
import multiprocessing as mp

import numpy as np
import pandas as pd
from scipy.optimize import minimize
from scipy.stats import gamma, norm, t, beta, weibull_min, kstest, linregress, genpareto, rayleigh
from scipy.interpolate import RegularGridInterpolator
from scipy.ndimage import zoom as scipy_zoom
from joblib import Parallel, delayed

import matplotlib.pyplot as plt
import matplotlib.colors as mcolors
import matplotlib.ticker as ticker
import seaborn as sns
from tqdm import tqdm

from cann_2d import CANNSimulator2D, calibrate_ff_scale_2d, measure_readout_mean_2d


# ============================================================
# UTILITY & FORMATTING FUNCTIONS
# ============================================================

def _grid_key_2d(Rf, wee, rf_decimals=3, wee_decimals=3):
    return (round(float(Rf), rf_decimals), round(float(wee), wee_decimals))


def _first_existing(candidates, columns, what):
    for c in candidates:
        if c in columns:
            return c
    raise ValueError(f"Could not find {what}. Tried: {candidates}")


def _resolve_2d_columns(df):
    cols = df.columns
    out = {
        "step_t": _first_existing(
            ["aic_dr_t2d", "aic_dr_t", "aic_dz_t", "aic_dd_t"],
            cols,
            "2D non-Gaussian step AIC column",
        ),
        "step_gauss": _first_existing(
            ["aic_dr_gauss", "aic_dz_gauss", "aic_dd_gauss"],
            cols,
            "2D Gaussian step AIC column",
        ),
        "tail_slope": None,
        "tail_alpha": None,
    }

    for c in ["slope_evt_dr", "slope_evt_dz", "slope_evt_dd"]:
        if c in cols:
            out["tail_slope"] = c
            break

    for c in ["alpha_evt_dr", "alpha_evt_dz", "alpha_evt_dd"]:
        if c in cols:
            out["tail_alpha"] = c
            break

    return out


def _format_slope_tick(x, pos=None):
    if abs(x - round(x)) < 1e-5:
        return f"{int(round(x))}"
    return f"{x:.2f}"


def _format_bof_tick(x, pos=None):
    if not np.isfinite(x):
        return ""
    ax = abs(x)
    if ax == 0:
        return "0"
    elif ax >= 1000:
        return f"{x:.0f}"
    elif ax >= 100:
        return f"{x:.0f}"
    elif ax >= 10:
        return f"{x:.1f}"
    elif ax >= 1:
        return f"{x:.2f}"
    else:
        return f"{x:.2g}"


def _get_symlog_levels(vmin, vmax, linthresh=10.0, num=100):
    if vmin >= vmax:
        return np.linspace(vmin - 1, vmax + 1, num)
        
    def forward(x):
        return np.sign(x) * np.log10(1 + np.abs(x) / linthresh)
        
    def inverse(x):
        return np.sign(x) * linthresh * (10**np.abs(x) - 1)
        
    t_min = forward(vmin)
    t_max = forward(vmax)
    t_levels = np.linspace(t_min, t_max, num)
    return inverse(t_levels)


def _soft_red_blue_cmap():
    return mcolors.LinearSegmentedColormap.from_list(
        "soft_red_blue",
        [
            (0.00, "#2166ac"),
            (0.25, "#67a9cf"),
            (0.50, "#ffffff"),
            (0.75, "#fddbc7"),
            (1.00, "#f4a582"),
        ],
        N=256,
    )


def _truncated_blues_r_cmap(light_cut=0.88):
    base = plt.get_cmap("Blues_r")
    colors = base(np.linspace(0.0, light_cut, 256))
    return mcolors.LinearSegmentedColormap.from_list("truncated_Blues_r", colors, N=256)


def _zero_centered_diverging_scale(Z, neg_pct=5, pos_pct=95, n_neg=60, n_pos=60, center=0.0):
    Z = np.asarray(Z, dtype=float)
    finite = Z[np.isfinite(Z)]

    if finite.size == 0:
        raise ValueError("Z contains no finite values.")

    neg_vals = finite[finite < center]
    pos_vals = finite[finite > center]

    vmin = np.nanpercentile(neg_vals, neg_pct) if neg_vals.size > 0 else center - 1.0
    vmax = np.nanpercentile(pos_vals, pos_pct) if pos_vals.size > 0 else center + 1.0

    if not np.isfinite(vmin) or vmin >= center:
        vmin = np.nanmin(finite)
        if vmin >= center:
            vmin = center - 1.0

    if not np.isfinite(vmax) or vmax <= center:
        vmax = np.nanmax(finite)
        if vmax <= center:
            vmax = center + 1.0

    if vmin == center: vmin = center - 1e-6
    if vmax == center: vmax = center + 1e-6

    norm = mcolors.TwoSlopeNorm(vmin=vmin, vcenter=center, vmax=vmax)

    neg_levels = np.linspace(vmin, center, n_neg, endpoint=False)
    pos_levels = np.linspace(center, vmax, n_pos)

    levels = np.unique(np.concatenate([neg_levels, pos_levels]))

    if len(levels) < 2:
        levels = np.array([vmin, vmax])

    ticks = np.unique(
        np.concatenate([
            np.linspace(vmin, center, 4, endpoint=False),
            np.linspace(center, vmax, 5),
        ])
    )

    line_levels = np.unique(
        np.concatenate([
            np.linspace(vmin, center, 5, endpoint=False),
            np.linspace(center, vmax, 5),
        ])
    )

    return levels, norm, ticks, line_levels, vmin, vmax


def _dr_blues_raw_delta_scale(Z, low_pct=2, high_pct=98, n_levels=100, n_ticks=6):
    Z = np.asarray(Z, dtype=float)
    finite = Z[np.isfinite(Z)]

    if finite.size == 0:
        raise ValueError("Z contains no finite values.")

    vmin = np.nanpercentile(finite, low_pct)
    vmax = np.nanpercentile(finite, high_pct)

    if not np.isfinite(vmin): vmin = np.nanmin(finite)
    if not np.isfinite(vmax): vmax = np.nanmax(finite)
    if vmax <= vmin: vmax = vmin + 1.0

    levels = np.linspace(vmin, vmax, n_levels)
    ticks = np.linspace(vmin, vmax, n_ticks)
    line_levels = np.linspace(vmin, vmax, n_ticks)

    norm = mcolors.Normalize(vmin=vmin, vmax=vmax)

    return levels, norm, ticks, line_levels, vmin, vmax


def _safe_contour_labels(ax, X, Y, Z, levels, fmt_func, linewidths=0.8, alpha=0.45, fontsize=9):
    Z = np.asarray(Z, dtype=float)
    if not np.any(np.isfinite(Z)):
        return None

    zmin, zmax = np.nanmin(Z), np.nanmax(Z)
    levels = np.asarray(levels, dtype=float)
    levels = levels[np.isfinite(levels)]
    levels = levels[(levels >= zmin) & (levels <= zmax)]
    levels = np.unique(levels)

    if len(levels) == 0:
        return None

    try:
        cl = ax.contour(X, Y, Z, levels=levels, colors="black", linewidths=linewidths, alpha=alpha)
        ax.clabel(cl, inline=True, inline_spacing=5, fontsize=fontsize, fmt=fmt_func)
        return cl
    except Exception:
        return None


def _set_contourf_edgecolor_face(cf):
    try:
        cf.set_edgecolor("face")
    except Exception:
        try:
            for coll in cf.collections:
                coll.set_edgecolor("face")
        except Exception:
            pass


def _smooth_grid_interp(x, y, z, zoom=5):
    interp = RegularGridInterpolator((y, x), z, method='linear')
    x_smooth = np.linspace(x.min(), x.max(), len(x) * zoom)
    y_smooth = np.linspace(y.min(), y.max(), len(y) * zoom)
    X_smooth, Y_smooth = np.meshgrid(x_smooth, y_smooth)
    pts = np.array([Y_smooth.ravel(), X_smooth.ravel()]).T
    z_smooth = interp(pts).reshape(X_smooth.shape)
    return X_smooth, Y_smooth, z_smooth


def _smooth_grid_zoom(x_vals, y_vals, Z, zoom=8, order=3):
    x_vals, y_vals, Z = np.asarray(x_vals, dtype=float), np.asarray(y_vals, dtype=float), np.asarray(Z, dtype=float)
    if Z.ndim != 2: raise ValueError("Z must be a 2D array.")
    
    if len(x_vals) < 2 or len(y_vals) < 2:
        X, Y = np.meshgrid(x_vals, y_vals)
        return X, Y, Z

    try:
        Z_smooth = scipy_zoom(Z, zoom=zoom, order=order)
        x_smooth = np.linspace(x_vals.min(), x_vals.max(), Z_smooth.shape[1])
        y_smooth = np.linspace(y_vals.min(), y_vals.max(), Z_smooth.shape[0])
        X_smooth, Y_smooth = np.meshgrid(x_smooth, y_smooth)
        return X_smooth, Y_smooth, Z_smooth
    except Exception:
        X, Y = np.meshgrid(x_vals, y_vals)
        return X, Y, Z


def _fill_pivot_nans_2d(pivot):
    pivot = pivot.copy()
    pivot.interpolate(method="linear", axis=1, limit_direction="both", inplace=True)
    pivot.interpolate(method="linear", axis=0, limit_direction="both", inplace=True)
    pivot = pivot.bfill(axis=0).ffill(axis=0).bfill(axis=1).ffill(axis=1).fillna(0.0)
    return pivot


# ============================================================
# STATISTICAL FITTING
# ============================================================

def _safe_aic_bic_from_logpdf(logpdf_vals, n_params):
    logpdf_vals = np.asarray(logpdf_vals, dtype=float)
    logpdf_vals = logpdf_vals[np.isfinite(logpdf_vals)]
    n = len(logpdf_vals)
    if n < 10:
        return np.nan, np.nan
    nll = -np.sum(logpdf_vals)
    aic = 2 * n_params + 2 * nll
    bic = n_params * np.log(n) + 2 * nll
    return float(aic), float(bic)


def _analyze_bh_U_distribution_fit(steady_state_data):
    x = np.asarray(steady_state_data, dtype=float)
    x = x[np.isfinite(x) & (x > 0)]
    if x.size < 10:
        return None

    out = {}
    try:
        gamma_params = gamma.fit(x, floc=0)
        gamma_logpdf = gamma.logpdf(x, *gamma_params)
        gamma_aic, gamma_bic = _safe_aic_bic_from_logpdf(gamma_logpdf, n_params=2)
        out["Gamma"] = {"params": gamma_params, "aic": gamma_aic, "bic": gamma_bic}
    except Exception:
        out["Gamma"] = {"params": (np.nan, 0.0, np.nan), "aic": np.nan, "bic": np.nan}

    try:
        norm_params = norm.fit(x)
        norm_logpdf = norm.logpdf(x, *norm_params)
        norm_aic, norm_bic = _safe_aic_bic_from_logpdf(norm_logpdf, n_params=2)
        out["Normal"] = {"params": norm_params, "aic": norm_aic, "bic": norm_bic}
    except Exception:
        out["Normal"] = {"params": (np.nan, np.nan), "aic": np.nan, "bic": np.nan}

    return out


def _analyze_signed_component_fit(x):
    x = np.asarray(x, dtype=float)
    x = x[np.isfinite(x)]
    if x.size < 10:
        return None

    out = {}
    try:
        t_params = t.fit(x)
        t_logpdf = t.logpdf(x, *t_params)
        t_aic, t_bic = _safe_aic_bic_from_logpdf(t_logpdf, n_params=3)
        out["Student_T"] = {"params": t_params, "aic": t_aic, "bic": t_bic}
    except Exception:
        out["Student_T"] = {"params": (np.nan, np.nan, np.nan), "aic": np.nan, "bic": np.nan}

    try:
        n_params = norm.fit(x)
        n_logpdf = norm.logpdf(x, *n_params)
        n_aic, n_bic = _safe_aic_bic_from_logpdf(n_logpdf, n_params=2)
        out["Normal"] = {"params": n_params, "aic": n_aic, "bic": n_bic}
    except Exception:
        out["Normal"] = {"params": (np.nan, np.nan), "aic": np.nan, "bic": np.nan}

    return out


def radial_t2d_pdf(r, kappa, A):
    r = np.asarray(r, dtype=float)
    out = np.zeros_like(r, dtype=float)
    m = r > 0
    out[m] = (2.0 * kappa / A) * r[m] * (1.0 + (r[m] ** 2) / A) ** (-(kappa + 1.0))
    return out


def radial_t2d_logpdf(r, kappa, A):
    r = np.asarray(r, dtype=float)
    out = np.full_like(r, -np.inf, dtype=float)
    m = r > 0
    out[m] = np.log(2.0 * kappa / A) + np.log(r[m]) - (kappa + 1.0) * np.log1p((r[m] ** 2) / A)
    return out


def _fit_radial_t2d(dr):
    dr = np.asarray(dr, dtype=float)
    dr = dr[np.isfinite(dr) & (dr > 1e-12)]
    if dr.size < 10:
        return {"kappa": np.nan, "A": np.nan, "aic": np.nan, "bic": np.nan}

    m2 = np.mean(dr ** 2)
    kappa0 = 3.0
    A0 = max((kappa0 - 1.0) * m2, 1e-8)

    def obj(log_params):
        log_kappa, log_A = log_params
        kappa = np.exp(log_kappa)
        A = np.exp(log_A)
        ll = radial_t2d_logpdf(dr, kappa, A)
        if not np.all(np.isfinite(ll)):
            return np.inf
        return -np.sum(ll)

    res = minimize(obj, x0=np.log([kappa0, A0]), method="L-BFGS-B", bounds=[(-10, 10), (-20, 20)])

    if not res.success:
        return {"kappa": np.nan, "A": np.nan, "aic": np.nan, "bic": np.nan}

    kappa_hat = float(np.exp(res.x[0]))
    A_hat = float(np.exp(res.x[1]))
    ll = radial_t2d_logpdf(dr, kappa_hat, A_hat)
    aic, bic = _safe_aic_bic_from_logpdf(ll, n_params=2)
    return {"kappa": kappa_hat, "A": A_hat, "aic": aic, "bic": bic}


def _fit_rayleigh_radius(dr):
    dr = np.asarray(dr, dtype=float)
    dr = dr[np.isfinite(dr) & (dr > 1e-12)]
    if dr.size < 10:
        return {"scale": np.nan, "aic": np.nan, "bic": np.nan}
    try:
        _, sigma = rayleigh.fit(dr, floc=0)
        ll = rayleigh.logpdf(dr, loc=0, scale=sigma)
        aic, bic = _safe_aic_bic_from_logpdf(ll, n_params=1)
        return {"scale": float(sigma), "aic": aic, "bic": bic}
    except Exception:
        return {"scale": np.nan, "aic": np.nan, "bic": np.nan}


def _fit_gaussian_radius(dr):
    dr = np.asarray(dr, dtype=float)
    dr = dr[np.isfinite(dr) & (dr > 1e-12)]
    if dr.size < 10:
        return {"loc": np.nan, "scale": np.nan, "aic": np.nan, "bic": np.nan}
    try:
        mu, sigma = norm.fit(dr)
        ll = norm.logpdf(dr, loc=mu, scale=sigma)
        aic, bic = _safe_aic_bic_from_logpdf(ll, n_params=2)
        return {"loc": float(mu), "scale": float(sigma), "aic": aic, "bic": bic}
    except Exception:
        return {"loc": np.nan, "scale": np.nan, "aic": np.nan, "bic": np.nan}


def fit_linear_tail(x, tail_q=0.95):
    x = np.asarray(x)
    x = x[np.isfinite(x) & (x > 1e-12)]
    if x.size < 200:
        return dict(slope=np.nan, intercept=np.nan, u=np.nan, r_value=np.nan, tail_ok=False)

    u = float(np.quantile(x, tail_q))
    x_tail = np.sort(x[x > u])
    
    if x_tail.size < 10:
        return dict(slope=np.nan, intercept=np.nan, u=u, r_value=np.nan, tail_ok=False)

    N = len(x)
    ranks = np.searchsorted(np.sort(x), x_tail, side='right')
    ccdf_tail = (N - ranks + 1) / N

    log_x = np.log10(x_tail)
    log_y = np.log10(ccdf_tail)

    try:
        slope, intercept, r_value, p_value, std_err = linregress(log_x, log_y)
        return dict(slope=slope, intercept=intercept, r_value=r_value, u=u, tail_ok=True)
    except:
        return dict(slope=np.nan, intercept=np.nan, u=u, r_value=np.nan, tail_ok=False)


def fit_evt_tail_metric(x, min_q=0.80, max_q=0.98, num_steps=50):
    x = np.asarray(x, dtype=float)
    x = x[np.isfinite(x) & (x > 1e-12)]

    if x.size < 200:
        return dict(alpha_evt=np.nan, linear_slope=np.nan, linear_intercept=np.nan, linear_r_squared=np.nan, u=np.nan, tail_ok=False)

    candidate_qs = np.linspace(min_q, max_q, num_steps)
    best_r2, best = -np.inf, None

    for q in candidate_qs:
        res = fit_linear_tail(x, tail_q=q)
        if not res["tail_ok"]: continue
        r2 = res["r_value"] ** 2
        if r2 > best_r2:
            best_r2, best = r2, res

    if best is None:
        return dict(alpha_evt=np.nan, linear_slope=np.nan, linear_intercept=np.nan, linear_r_squared=np.nan, u=np.nan, tail_ok=False)

    slope = best["slope"]
    alpha_evt = float(min(-slope, 2.0)) if (np.isfinite(slope) and slope < 0) else 2.0

    return dict(
        alpha_evt=alpha_evt,
        linear_slope=float(best["slope"]),
        linear_intercept=float(best["intercept"]),
        linear_r_squared=float(best["r_value"] ** 2),
        u=float(best["u"]),
        tail_ok=True,
    )


# ============================================================
# DATA PREPARATION & PROCESSING
# ============================================================

def rolling_mean_time(data, window, axis=0):
    arr = np.asarray(data)
    if window <= 1:
        return arr

    if arr.ndim == 1:
        if arr.shape[0] < window:
            return np.empty((0,), dtype=float)
        c = np.cumsum(np.insert(arr.astype(float), 0, 0.0))
        return (c[window:] - c[:-window]) / window

    if axis != 0:
        arr = np.swapaxes(arr, 0, axis)

    T = arr.shape[0]
    if T < window:
        out = np.empty((0,) + arr.shape[1:], dtype=float)
        if axis != 0: out = np.swapaxes(out, 0, axis)
        return out

    c = np.cumsum(arr.astype(float), axis=0)
    c = np.concatenate([np.zeros((1,) + arr.shape[1:], dtype=float), c], axis=0)
    out = (c[window:] - c[:-window]) / window

    if axis != 0:
        out = np.swapaxes(out, 0, axis)
    return out


def sliding_average(arr, window):
    arr = np.asarray(arr)
    if len(arr) < window: return np.array([])
    return np.convolve(arr, np.ones(window)/window, mode='valid')


def prepare_integrated_data(time, z_list, Ue_list, delta_z, params, win_bins=1):
    recording_start = params.get("recording_start", 0.0)

    z_arr = np.asarray(z_list, dtype=float)
    U_arr = np.asarray(Ue_list, dtype=float)
    dz_arr = np.asarray(delta_z)

    L = min(len(time), len(z_arr), len(U_arr), len(dz_arr))
    time_s = np.asarray(time[:L], dtype=float)
    z_s = z_arr[:L]
    U_s = U_arr[:L]
    dz_s = dz_arr[:L]

    if z_s.ndim != 2 or z_s.shape[1] != 2:
        raise ValueError("z_list must have shape (T, 2).")

    if dz_s.ndim == 2 and dz_s.shape[1] == 2:
        dx_s = dz_s[:, 0].astype(float)
        dy_s = dz_s[:, 1].astype(float)
        dr_s = np.sqrt(dx_s ** 2 + dy_s ** 2)
    else:
        dr_s = dz_s.reshape(-1).astype(float)
        dx_s = np.diff(z_s[:, 0], prepend=np.nan).astype(float)
        dy_s = np.diff(z_s[:, 1], prepend=np.nan).astype(float)

    U_dec  = sliding_average(U_s,  win_bins)
    dr_dec = sliding_average(dr_s, win_bins)
    dx_dec = sliding_average(dx_s, win_bins)
    dy_dec = sliding_average(dy_s, win_bins)
    time_dec = time_s[win_bins - 1:]

    valid = (
        (time_dec >= recording_start)
        & np.isfinite(U_dec)  & (U_dec > 0)
        & np.isfinite(dr_dec) & (dr_dec > 1e-12)
        & np.isfinite(dx_dec)
        & np.isfinite(dy_dec)
    )

    return time_s, time_dec[valid], U_dec[valid], dr_dec[valid], dx_dec[valid], dy_dec[valid]


def run_global_fits(U_dec_clean, dr_dec_clean, dx_dec_clean, dy_dec_clean):
    return {
        "U": _analyze_bh_U_distribution_fit(U_dec_clean) or {},
        "dr": {
            "Radial_T2D": _fit_radial_t2d(dr_dec_clean),
            "Normal": _fit_gaussian_radius(dr_dec_clean),
        },
        "dr_tail": fit_evt_tail_metric(dr_dec_clean) or {},
        "dx": (_analyze_signed_component_fit(dx_dec_clean) if dx_dec_clean.size > 0 else {}) or {},
        "dy": (_analyze_signed_component_fit(dy_dec_clean) if dy_dec_clean.size > 0 else {}) or {},
    }


def perform_integrated_analysis_2d(time, z_list, Ue_list, delta_z, params, Rf, wee, k, output_dir, save_diagnostic_plot=True, plot_range=180):
    win_bins = 1
    time_s, time_dec, U_dec, dr_dec, dx_dec, dy_dec = prepare_integrated_data(time, z_list, Ue_list, delta_z, params, win_bins)

    fits = run_global_fits(U_dec, dr_dec, dx_dec, dy_dec)
    fig_dist_path = ""

    if save_diagnostic_plot:
        fig_dist = plot_distributions(U_dec, dr_dec, dx_dec, dy_dec, fits, Rf, wee, k, win_bins, plot_range)
        os.makedirs(output_dir, exist_ok=True)
        fig_dist_path = os.path.join(output_dir, f"diagnostics_Rf_{Rf:.6f}_k_{k:.7f}_wee_{wee:.6f}.svg")
        fig_dist.savefig(fig_dist_path, dpi=150, bbox_inches="tight")
        plt.close(fig_dist)

    u_fit = fits.get("U") or {}
    dr_fit = fits.get("dr") or {}
    dx_fit = fits.get("dx") or {}
    dy_fit = fits.get("dy") or {}
    tail_fit = fits.get("dr_tail") or {}

    return {
        "Rf": Rf,
        "wee": wee,
        "k": k,
        "kappa_fit_U": u_fit.get("Gamma", {}).get("params", [np.nan])[0],
        "kappa_fit_dr": dr_fit.get("Radial_T2D", {}).get("kappa", np.nan),
        "A_fit_dr": dr_fit.get("Radial_T2D", {}).get("A", np.nan),
        "alpha_evt_dr": tail_fit.get("alpha_evt", np.nan),
        "slope_evt_dr": tail_fit.get("linear_slope", np.nan),
        "aic_U_gamma": u_fit.get("Gamma", {}).get("aic", np.nan),
        "aic_U_gauss": u_fit.get("Normal", {}).get("aic", np.nan),
        "aic_dx_t": dx_fit.get("Student_T", {}).get("aic", np.nan),
        "aic_dx_gauss": dx_fit.get("Normal", {}).get("aic", np.nan),
        "aic_dy_t": dy_fit.get("Student_T", {}).get("aic", np.nan),
        "aic_dy_gauss": dy_fit.get("Normal", {}).get("aic", np.nan),
        "aic_dr_t2d": dr_fit.get("Radial_T2D", {}).get("aic", np.nan),
        "aic_dr_gauss": dr_fit.get("Normal", {}).get("aic", np.nan),
        "fig_dist_path": fig_dist_path,
    }


def analyze_saved_data_task_2d(args):
    data_path, params, plot_output_dir, save_diagnostic_plot, plot_range = args
    try:
        with np.load(data_path, allow_pickle=True) as data:
            out = perform_integrated_analysis_2d(
                time=data['time'],
                z_list=data['z_list'],
                Ue_list=data['Ue_list'],
                delta_z=data['delta_z_mag'],
                params=params,
                Rf=float(data['Rf']),
                wee=float(data['wee']),
                k=float(data['k']),
                output_dir=plot_output_dir,
                save_diagnostic_plot=save_diagnostic_plot,
                plot_range=plot_range
            )
            return out
    except Exception as e:
        print(f"Analysis Error on {os.path.basename(data_path)}: {e}")
        return None


def _collect_or_build_summary_2d_fixed_k(
    params, target_k, Rf_list, wee_list, raw_data_dir="output_net/raw_npz/2d/",
    plot_dir="output_net/plots/2d/", n_jobs=-1, force_reanalyze=False,
    save_diagnostic_plot=False, save_summary_csv=True, atol_k=1e-12, plot_range=180
):
    os.makedirs(raw_data_dir, exist_ok=True)
    os.makedirs(plot_dir, exist_ok=True)

    target_pairs = {_grid_key_2d(Rf, wee, rf_decimals=3, wee_decimals=3) for Rf in Rf_list for wee in wee_list}
    required_cols = {"Rf", "k", "wee", "aic_U_gamma", "aic_U_gauss", "aic_dr_t2d", "aic_dr_gauss", "aic_dx_t", "aic_dx_gauss", "aic_dy_t", "aic_dy_gauss", "slope_evt_dr"}

    df_parts = []
    if not force_reanalyze:
        for pattern in ["summary_2d_fixed_k_*.csv", "summary_2d_fixed_wee_*.csv"]:
            for f in glob.glob(os.path.join(plot_dir, pattern)):
                try:
                    tmp = pd.read_csv(f)
                    if required_cols.issubset(tmp.columns):
                        tmp = tmp[np.isclose(tmp["k"], target_k, rtol=0, atol=atol_k)]
                        if not tmp.empty: df_parts.append(tmp)
                except Exception as e:
                    print(f"Warning: failed to read {f}: {e}")

    df = pd.concat(df_parts, ignore_index=True) if df_parts else pd.DataFrame()

    if not df.empty:
        df = df.drop_duplicates(subset=["Rf", "wee", "k"], keep="last")
        have_pairs = {_grid_key_2d(rf, wee, rf_decimals=3, wee_decimals=3) for rf, wee in zip(df["Rf"], df["wee"])}
    else:
        have_pairs = set()

    missing_pairs = sorted(target_pairs - have_pairs)

    if missing_pairs:
        missing_sim_tasks = []
        for Rf_key, wee_key in missing_pairs:
            fn = f"raw_data_2d_Rf_{Rf_key:.6f}_k_{target_k:.7f}_wee_{wee_key:.6f}.npz"
            p = os.path.join(raw_data_dir, fn)
            if not os.path.exists(p):
                wee_norm = wee_key / params.get("critical_weight", 1.0)
                missing_sim_tasks.append((params, Rf_key, wee_key, wee_norm, target_k, raw_data_dir, plot_dir))

        if missing_sim_tasks:
            max_workers = mp.cpu_count() if n_jobs == -1 else n_jobs
            with concurrent.futures.ProcessPoolExecutor(max_workers=max_workers) as executor:
                futures = [executor.submit(simulate_plot_and_save_task_2d, task) for task in missing_sim_tasks]
                for future in tqdm(concurrent.futures.as_completed(futures), total=len(futures), desc="Simulating 2D"):
                    try: future.result()
                    except Exception as e: print(f"\nSimulation worker crashed: {e}")

        missing_npz_paths = []
        for Rf_key, wee_key in missing_pairs:
            fn = f"raw_data_2d_Rf_{Rf_key:.6f}_k_{target_k:.7f}_wee_{wee_key:.6f}.npz"
            p = os.path.join(raw_data_dir, fn)
            if os.path.exists(p): missing_npz_paths.append(p)

        if missing_npz_paths:
            max_workers = mp.cpu_count() if n_jobs == -1 else n_jobs
            tasks = [(path, params, plot_dir, save_diagnostic_plot, plot_range) for path in missing_npz_paths]
            results = []
            with concurrent.futures.ProcessPoolExecutor(max_workers=max_workers) as executor:
                futures = [executor.submit(analyze_saved_data_task_2d, task) for task in tasks]
                for future in tqdm(concurrent.futures.as_completed(futures), total=len(futures), desc="Analyzing 2D"):
                    try:
                        res = future.result()
                        if res is not None: results.append(res)
                    except Exception as e:
                        print(f"\nAnalysis worker crashed: {e}")

            if results:
                df_new = pd.DataFrame(results).sort_values(["Rf", "wee"])
                if save_summary_csv:
                    summary_path = os.path.join(plot_dir, f"summary_2d_fixed_k_{target_k:.7f}_grid.csv")
                    df_new.to_csv(summary_path, index=False)

                df = pd.concat([df, df_new], ignore_index=True) if not df.empty else df_new
                df = df.drop_duplicates(subset=["Rf", "wee", "k"], keep="last")

    if df.empty:
        raise ValueError("No valid 2D summary data available after checking CSV, NPZ, and simulations.")

    df = df[np.isclose(df["k"], target_k, rtol=0, atol=atol_k)].copy()
    keep_mask = [_grid_key_2d(rf, wee, rf_decimals=3, wee_decimals=3) in target_pairs for rf, wee in zip(df["Rf"], df["wee"])]
    df = df[keep_mask].copy()

    if df.empty:
        raise ValueError("No rows matched the requested (Rf, wee) grid at the target k.")

    return df.sort_values(["Rf", "wee"]).reset_index(drop=True)


# ============================================================
# PLOTTING FUNCTIONS
# ============================================================

def plot_distributions(U, dr, dx, dy, fits, Rf, wee, k, win_bins, plot_range=180):
    fig = plt.figure(figsize=(24, 18), constrained_layout=True)
    gs = fig.add_gridspec(2, 4)

    ax_dist_dx = fig.add_subplot(gs[0, 0:2])
    ax_dist_dy = fig.add_subplot(gs[0, 2:4])    
    ax_dist_u  = fig.add_subplot(gs[1, 0])
    ax_dist_dz = fig.add_subplot(gs[1, 1])
    ax_loglog  = fig.add_subplot(gs[1, 2:])

    fig.suptitle(f"Distributions (Rf={Rf:.6f}, k={k:.7f}, wee={wee:.6f}) | win_bins={win_bins}", fontsize=22)

    def plot_thinned_empirical_ccdf(ax, r_sorted, ccdf_emp):
        max_points = 400
        if len(r_sorted) > max_points:
            step = len(r_sorted) // max_points
            r_plot = r_sorted[::step]
            ccdf_plot = ccdf_emp[::step]
            if r_plot[-1] != r_sorted[-1]:
                r_plot = np.append(r_plot, r_sorted[-1])
                ccdf_plot = np.append(ccdf_plot, ccdf_emp[-1])
        else:
            r_plot = r_sorted
            ccdf_plot = ccdf_emp
        ax.loglog(r_plot, ccdf_plot, "o", ms=4, mfc="white", mec="0.45", mew=1.0, alpha=0.65, label="Empirical CCDF")

    sns.histplot(U, bins=50, kde=False, stat="density", ax=ax_dist_u, color="lightsteelblue", edgecolor="None", linewidth=0)
    u_dict = fits.get("U") or {}
    
    if U.size > 0:
        x_u = np.linspace(np.min(U)*0.9, np.max(U) * 1.1, 300)
        u_gamma = u_dict.get("Gamma", {})
        k_val, loc_g, th = u_gamma.get("params", [np.nan, 0, np.nan])
        aic_gamma = u_gamma.get("aic", np.nan)
        if np.isfinite(k_val) and np.isfinite(th):
            ax_dist_u.plot(x_u, gamma.pdf(x_u, k_val, loc_g, th), color="#1f4e79", ls="--", lw=3, label=f"Gamma (AIC: {aic_gamma:.1f})")
            
        u_gauss = u_dict.get("Normal", {})
        g_params = u_gauss.get("params", [np.nan, np.nan])
        if len(g_params) < 2 or not np.isfinite(g_params[0]):
            g_mu, g_std = norm.fit(U)
            aic_gauss_u = 4 + 2 * -np.sum(norm.logpdf(U, g_mu, g_std))
        else:
            g_mu, g_std = g_params
            aic_gauss_u = u_gauss.get("aic", np.nan)

        if np.isfinite(g_std) and g_std > 0:
            ax_dist_u.plot(x_u, norm.pdf(x_u, g_mu, g_std), color="#d17c00", ls="--", lw=3, label=f"Gauss (AIC: {aic_gauss_u:.1f})")

        ax_dist_u.legend(fontsize=9)
    ax_dist_u.set_title("Smoothed Bump Height U")

    def plot_linear_tail_on_loglog(ax, data, fit_dict, label_prefix):
        if data.size > 0 and fit_dict is not None and fit_dict.get("tail_ok"):
            u_val = fit_dict["u"]
            slope = fit_dict["linear_slope"]
            intercept = fit_dict["linear_intercept"]
            r2 = fit_dict.get("linear_r_squared", np.nan)
            
            if np.isfinite(slope) and np.isfinite(intercept):
                r_tail = np.logspace(np.log10(max(u_val, np.min(data))), np.log10(np.max(data)), 300)
                ccdf_tail = (10 ** intercept) * (r_tail ** slope)
                lbl = f"{label_prefix} Tail (α={-slope:.2f}, R²={r2:.2f})"
                ax.plot(r_tail, ccdf_tail, "--", color="#005f73", lw=2, label=lbl)
                ax.axvline(u_val, color="0.65", lw=1.6, ls=":", alpha=0.9)

    def plot_histogram_diagnostics(ax, data, fit_dict, title):
        if data.size == 0: return
        fit_dict = fit_dict or {}
        st_dict = fit_dict.get("Student_T", {})
        nu, loc_val, scale_val = st_dict.get("params", [np.nan, np.nan, np.nan])
        clean_data = data[np.isfinite(data)]
        if clean_data.size < 5: return
        
        if not np.isfinite(nu) and clean_data.size > 10:
            nu, loc_val, scale_val = t.fit(clean_data)
            aic_t = 6 + 2 * -np.sum(t.logpdf(clean_data, nu, loc_val, scale_val))
        else:
            aic_t = st_dict.get("aic", np.nan)

        ax.hist(clean_data, bins='auto', density=True, alpha=0.5, color="lightsteelblue", label='Empirical')
        x_grid = np.linspace(clean_data.min(), clean_data.max(), 400)
        
        if np.isfinite(nu):
            pdf_t = t.pdf(x_grid, df=nu, loc=loc_val, scale=scale_val)
            ax.plot(x_grid, pdf_t, "-", color="#1f4e79", lw=2, label=f"Student-t (AIC: {aic_t:.1f})")
                
        gauss_dict = fit_dict.get("Normal", {})
        gauss_params = gauss_dict.get("params", [np.nan, np.nan])
        if len(gauss_params) < 2 or not np.isfinite(gauss_params[0]):
            g_mu, g_std = norm.fit(clean_data)
            aic_g = 4 + 2 * -np.sum(norm.logpdf(clean_data, g_mu, g_std))
        else:
            g_mu, g_std = gauss_params
            aic_g = gauss_dict.get("aic", np.nan)

        if np.isfinite(g_std) and g_std > 0:
            pdf_g = norm.pdf(x_grid, loc=g_mu, scale=g_std)
            ax.plot(x_grid, pdf_g, "--", color="#d17c00", lw=2, label=f"Gauss (AIC: {aic_g:.1f})")
        
        ax.set_title(f"{title} Distribution Diagnostics (t vs Gauss)")
        ax.set_xlabel(title)
        ax.set_ylabel("Density")
        ax.legend(fontsize=9)

    if fits.get("dx") is not None or dx.size > 0: 
        plot_histogram_diagnostics(ax_dist_dx, dx, fits.get("dx"), "dx")
    if fits.get("dy") is not None or dy.size > 0: 
        plot_histogram_diagnostics(ax_dist_dy, dy, fits.get("dy"), "dy")

    dr_dict = fits.get("dr", {})
    dr_t2d_dict = dr_dict.get("Radial_T2D", {})
    aic_t2d_dr = dr_t2d_dict.get("aic", np.nan)
    
    dr_gauss_dict = dr_dict.get("Normal", {})
    dr_gauss_params = dr_gauss_dict.get("params", [np.nan, np.nan])
    
    if dr.size > 5:
        if len(dr_gauss_params) < 2 or not np.isfinite(dr_gauss_params[0]):
            g_mu_dr, g_std_dr = norm.fit(dr)
            dr_clean = dr[dr > 0] 
            aic_gauss_dr = 4 + 2 * -np.sum(norm.logpdf(dr_clean, g_mu_dr, g_std_dr))
        else:
            g_mu_dr, g_std_dr = dr_gauss_params
            aic_gauss_dr = dr_gauss_dict.get("aic", np.nan)

        rdata = dr[np.isfinite(dr) & (dr > 0)]
        r_sorted = np.sort(rdata)
        N = len(r_sorted)
        ccdf_emp = (N - np.arange(1, N + 1) + 1) / N
        
        plot_mask = r_sorted < plot_range
        r_sorted = r_sorted[plot_mask]
        ccdf_emp = ccdf_emp[plot_mask]
        plot_thinned_empirical_ccdf(ax_loglog, r_sorted, ccdf_emp)
        
        if len(r_sorted) > 0:
            r_grid = np.logspace(np.log10(r_sorted.min()), np.log10(r_sorted.max()), 400)
            if np.isfinite(dr_t2d_dict.get("kappa", np.nan)):
                kappa = dr_t2d_dict["kappa"]
                A = dr_t2d_dict["A"]
                ccdf_t2d = (1.0 + (r_grid**2) / A) ** (-kappa)
                ax_loglog.loglog(r_grid, ccdf_t2d, "-", color="#1f4e79", lw=2, label=f"Radial T2D (AIC: {aic_t2d_dr:.1f})")
            if np.isfinite(g_std_dr) and g_std_dr > 0:
                ccdf_g_dr = 1.0 - norm.cdf(r_grid, loc=g_mu_dr, scale=g_std_dr)
                ax_loglog.loglog(r_grid, ccdf_g_dr, "--", color="#d17c00", lw=2, label=f"Gauss (AIC: {aic_gauss_dr:.1f})")
        
        if fits.get("dr_tail"):
            plot_linear_tail_on_loglog(ax_loglog, rdata, fits["dr_tail"], "dr")
        
        ax_loglog.set_xlabel("Step Size dr")
        ax_loglog.set_ylabel("CCDF")
        ax_loglog.legend(fontsize=9)
            
    ax_loglog.set_title("dr Tail Diagnostics (T2D vs Gauss)")

    max_dr = np.max(dr) if dr.size > 0 else plot_range
    sns.histplot(dr, bins=100, binrange=(0, min(max_dr, plot_range)), kde=False, stat="density", ax=ax_dist_dz, color="lightsteelblue", edgecolor="None", linewidth=0)
    
    if dr.size > 0:
        x_dr = np.linspace(0, min(max_dr, plot_range), 1000)
        if np.isfinite(dr_t2d_dict.get("kappa", np.nan)):
            kappa = dr_t2d_dict["kappa"]
            A = dr_t2d_dict["A"]
            ax_dist_dz.plot(x_dr, radial_t2d_pdf(x_dr, kappa, A), color="#1f4e79", ls="--", lw=3, label=f"Radial T2D (AIC: {aic_t2d_dr:.1f})")
        if np.isfinite(g_std_dr) and g_std_dr > 0:
            ax_dist_dz.plot(x_dr, norm.pdf(x_dr, g_mu_dr, g_std_dr), color="#d17c00", ls="--", lw=3, label=f"Gauss (AIC: {aic_gauss_dr:.1f})")
        ax_dist_dz.legend(fontsize=9)
            
    ax_dist_dz.set_title("Smoothed Step Size Magnitude (dr)")
    ax_dist_dz.set_xlim(0, plot_range)

    return fig


def plot_heatmaps(time_s, time_dec, Oc, win_bins, Rf, wee, k):
    fig_hm = plt.figure(figsize=(12, 4), constrained_layout=True)
    gs_hm = fig_hm.add_gridspec(1, 5)

    ax_heat_x = fig_hm.add_subplot(gs_hm[0, 0:2])
    ax_heat_y = fig_hm.add_subplot(gs_hm[0, 2:4]) 
    ax_snap   = fig_hm.add_subplot(gs_hm[0, 4]) 
    
    fig_hm.suptitle(f"2D Marginals & Snapshot (Rf={Rf:.6f}, k = {k:.7f}, wee={wee:.6f})", fontsize=16)

    def plot_marginal_map(ax, data, title, axis_to_sum, cmap="viridis"):
        if data is None or len(data) == 0:
            ax.set_axis_off()
            return
        arr_marginal = np.sum(np.asarray(data)[:len(time_s)], axis=axis_to_sum)
        arr_roll = rolling_mean_time(arr_marginal, window=win_bins, axis=0) 
        if arr_roll.size == 0:
            ax.set_title(f"{title} (window too large)")
            return
        vmin, vmax = np.percentile(arr_roll, [1, 99])
        im = ax.imshow(arr_roll.T, aspect="auto", origin="lower", extent=[time_dec[0], time_dec[-1], -180, 180], vmin=vmin, vmax=vmax, cmap=cmap)
        ax.set_title(title)
        ax.set_xlabel("Time (s)")
        ax.set_ylabel("Position (deg)")
        plt.colorbar(im, ax=ax, fraction=0.046, pad=0.04)

    if Oc is not None and len(Oc) > 0:
        plot_marginal_map(ax_heat_x, Oc, "Firing Rate X", axis_to_sum=2)
        plot_marginal_map(ax_heat_y, Oc, "Firing Rate Y", axis_to_sum=1, cmap="plasma")
        
        Oc_arr = np.asarray(Oc)[:len(time_s)]
        r_roll = rolling_mean_time(Oc_arr, window=win_bins, axis=0)
        if r_roll.size > 0:
            profile = r_roll[-1] 
            im_snap = ax_snap.imshow(profile.T, aspect="equal", origin="lower", extent=[-180, 180, -180, 180], cmap="viridis")
            ax_snap.set_title("Snapshot")
            plt.colorbar(im_snap, ax=ax_snap, fraction=0.046, pad=0.04)
        else:
            ax_snap.set_axis_off()
    else:
        ax_snap.set_axis_off()

    return fig_hm


def plot_metric_heatmap(data_matrix, x_vals, y_vals, title_str, cbar_label_str, wc=0.895612107936198, vmin=None, vmax=None):
    plt.figure(figsize=(10, 8))
    ax = sns.heatmap(
        data_matrix, 
        cmap='RdBu_r', 
        cbar_kws={'label': cbar_label_str},
        vmin=vmin, 
        vmax=vmax  
    )
    
    ax.set_title(title_str, fontsize=16)
    ax.set_xlabel(r'$w_{ee}(/wc)$', fontsize=12)
    ax.set_ylabel('$R_f$', fontsize=12)
    
    ax.set_xticks(np.linspace(0, len(x_vals)-1, 6, dtype=int))
    ax.set_xticklabels([f'{x/wc:.2f}' for x in x_vals[ax.get_xticks()]])
    ax.set_yticks(np.linspace(0, len(y_vals)-1, 6, dtype=int))
    ax.set_yticklabels([f'{y:.2f}' for y in y_vals[ax.get_yticks()]])
    
    plt.gca().invert_yaxis()
    plt.savefig(f"{title_str.replace(' ', '_').lower()}.svg", dpi=300, bbox_inches='tight')


def plot_tail_slope_vs_wee_rf_2d(
    params,
    target_k,
    Rf_list,
    wee_list,
    raw_data_dir="output_net/raw_npz/2d/",
    plot_dir="output_net/plots/2d/",
    n_jobs=-1,
    value_col=None,
    force_reanalyze=False,
    save=True,
    save_diagnostic_plot=False,
    show_wee_in_wc=True,
):
    plt.close('all') 
    
    df = _collect_or_build_summary_2d_fixed_k(
        params=params, target_k=target_k, Rf_list=Rf_list, wee_list=wee_list,
        raw_data_dir=raw_data_dir, plot_dir=plot_dir, n_jobs=n_jobs,
        force_reanalyze=force_reanalyze, save_diagnostic_plot=save_diagnostic_plot,
    )

    colmap = _resolve_2d_columns(df)
    if value_col is None:
        value_col = colmap["tail_slope"] if colmap["tail_slope"] is not None else colmap["tail_alpha"]

    if value_col is None or value_col not in df.columns:
        raise ValueError(f"{value_col} not found in summary dataframe.")

    df = df.dropna(subset=["Rf", "wee", value_col]).copy()
    if df.empty:
        raise ValueError(f"No valid 2D rows available for {value_col}.")

    pivot = df.pivot_table(index="Rf", columns="wee", values=value_col, aggfunc="mean").sort_index()

    y_vals = pivot.index.values.astype(float)
    x_vals = pivot.columns.values.astype(float)

    if show_wee_in_wc:
        sim_temp = CANNSimulator2D(params)
        wc = float(sim_temp.params["critical_weight"])
        x_vals = x_vals / wc
        x_label = r"$w_{ee}\;(\times w_c)$"
    else:
        x_label = "wee"

    Z = pivot.values
    X_smooth, Y_smooth, Z_smooth = _smooth_grid_interp(x_vals, y_vals, Z, zoom=8)

    fig, ax = plt.subplots(figsize=(8, 6), constrained_layout=True)

    vmin = np.nanpercentile(Z, 5)
    vmax = np.nanpercentile(Z, 95)
    levels_smooth = np.linspace(vmin, vmax, 100)

    cf = ax.contourf(X_smooth, Y_smooth, Z_smooth, levels=levels_smooth, cmap="YlGnBu", extend="both")
    cf.set_edgecolor("face")
    
    cbar = fig.colorbar(cf, ax=ax)
    cbar.locator = ticker.MaxNLocator(integer=True)
    cbar.update_ticks()

    line_levels = np.linspace(vmin, vmax, 8)
    c_lines = ax.contour(X_smooth, Y_smooth, Z_smooth, levels=line_levels, colors='black', linewidths=1, alpha=0.5)
    ax.clabel(c_lines, inline=True, inline_spacing=5, fontsize=10, fmt='%.1f')

    ax.set_title(f"2D tail metric across wee and Rf: {value_col} (k={target_k:.7f})")
    ax.set_xlabel(x_label)
    ax.set_ylabel("Rf")

    if save:
        suffix = "_wc" if show_wee_in_wc else ""
        out = os.path.join(plot_dir, f"{value_col}_vs_wee_rf_2d_k_{target_k:.7f}{suffix}.svg")
        fig.savefig(out, dpi=150, bbox_inches="tight")
        print(f"Saved: {out}")

    return fig, df, pivot


def plot_bof_vs_wee_rf_2d(
    params,
    target_k,
    Rf_list,
    wee_list,
    raw_data_dir="output_net/raw_npz/2d/",
    plot_dir="output_net/plots/2d/",
    n_jobs=-1,
    metric="delta_aic",
    force_reanalyze=False,
    save=True,
    save_diagnostic_plot=False,
    show_wee_in_wc=True,
    smooth_zoom=8,
    dr_light_cut=0.88, 
):
    plt.close("all")

    df = _collect_or_build_summary_2d_fixed_k(
        params=params, target_k=target_k, Rf_list=Rf_list, wee_list=wee_list,
        raw_data_dir=raw_data_dir, plot_dir=plot_dir, n_jobs=n_jobs,
        force_reanalyze=force_reanalyze, save_diagnostic_plot=save_diagnostic_plot,
    )

    req = ["Rf", "wee", "aic_U_gamma", "aic_U_gauss", "aic_dr_t2d", "aic_dr_gauss", "aic_dx_t", "aic_dx_gauss", "aic_dy_t", "aic_dy_gauss"]
    missing_cols = [c for c in req if c not in df.columns]
    if missing_cols: raise ValueError("The summary dataframe is missing required columns: " + ", ".join(missing_cols))

    df = df.dropna(subset=req).copy()
    if df.empty: raise ValueError("No valid 2D rows available for best-of-fit plotting.")

    df["delta_aic_U"] = df["aic_U_gamma"] - df["aic_U_gauss"]
    df["delta_aic_dr"] = df["aic_dr_t2d"] - df["aic_dr_gauss"]
    df["delta_aic_dx"] = df["aic_dx_t"] - df["aic_dx_gauss"]
    df["delta_aic_dy"] = df["aic_dy_t"] - df["aic_dy_gauss"]

    df["winner_U"] = (df["delta_aic_U"] < 0).astype(float)
    df["winner_dr"] = (df["delta_aic_dr"] < 0).astype(float)
    df["winner_dx"] = (df["delta_aic_dx"] < 0).astype(float)
    df["winner_dy"] = (df["delta_aic_dy"] < 0).astype(float)

    if metric not in ["delta_aic", "winner"]: raise ValueError("metric must be either 'delta_aic' or 'winner'.")

    value_cols = {
        "U": "delta_aic_U" if metric == "delta_aic" else "winner_U",
        "dr": "delta_aic_dr" if metric == "delta_aic" else "winner_dr",
        "dx": "delta_aic_dx" if metric == "delta_aic" else "winner_dx",
        "dy": "delta_aic_dy" if metric == "delta_aic" else "winner_dy",
    }

    pivots = {}
    for key, col in value_cols.items():
        pivot = df.pivot_table(index="Rf", columns="wee", values=col, aggfunc="mean").sort_index()
        pivots[key] = _fill_pivot_nans_2d(pivot)

    y_vals = pivots["U"].index.values.astype(float)
    x_vals_raw = pivots["U"].columns.values.astype(float)

    if show_wee_in_wc:
        sim_temp = CANNSimulator2D(params)
        wc = float(sim_temp.params["critical_weight"])
        x_vals = x_vals_raw / wc
        x_label = r"$w_{ee}\;(\times w_c)$"
    else:
        x_vals = x_vals_raw
        x_label = r"$w_{ee}$"

    smooth = {}
    for key, pivot in pivots.items():
        Z = pivot.values.astype(float)
        X_smooth, Y_smooth, Z_smooth = _smooth_grid_zoom(x_vals, y_vals, Z, zoom=smooth_zoom, order=3)
        smooth[key] = Z_smooth

    fig, axes = plt.subplots(2, 2, figsize=(18, 12), constrained_layout=True)

    plot_map = [("U", axes[0, 0]), ("dr", axes[0, 1]), ("dx", axes[1, 0]), ("dy", axes[1, 1])]

    if metric == "delta_aic":
        titles = {
            "U": r"U fit: $\Delta$AIC = AIC(Gamma) - AIC(Gaussian)",
            "dr": r"$dr$ fit: $\Delta$AIC = AIC(Radial Student-t) - AIC(Gaussian)",
            "dx": r"$dx$ fit: $\Delta$AIC = AIC(Student-t) - AIC(Gaussian)",
            "dy": r"$dy$ fit: $\Delta$AIC = AIC(Student-t) - AIC(Gaussian)",
        }
    else:
        titles = {
            "U": r"U winner: 1 = Gamma better",
            "dr": r"$dr$ winner: 1 = Radial Student-t better",
            "dx": r"$dx$ winner: 1 = Student-t better",
            "dy": r"$dy$ winner: 1 = Student-t better",
        }

    dr_cmap = _truncated_blues_r_cmap(light_cut=dr_light_cut)

    for key, ax in plot_map:
        ax.set_facecolor("#eeeeee")
        Z_raw = pivots[key].values.astype(float)
        Z_smooth = smooth[key].astype(float)

        if metric == "delta_aic":
            if key == "dr":
                levels, norm, ticks, line_levels, vmin, vmax = _dr_blues_raw_delta_scale(Z_raw)
                cf = ax.contourf(X_smooth, Y_smooth, Z_smooth, levels=levels, norm=norm, cmap=dr_cmap, extend="both")
                _set_contourf_edgecolor_face(cf)

                cbar = fig.colorbar(cf, ax=ax, ticks=ticks)
                cbar.ax.yaxis.set_major_formatter(ticker.FuncFormatter(_format_bof_tick))
                cbar.set_label(r"$\Delta$AIC = AIC(Radial Student-t) - AIC(Gaussian)", fontsize=10)

                _safe_contour_labels(ax=ax, X=X_smooth, Y=Y_smooth, Z=Z_smooth, levels=line_levels, fmt_func=_format_bof_tick, linewidths=0.8, alpha=0.45, fontsize=9)
            else:
                levels, norm, ticks, line_levels, vmin, vmax = _zero_centered_diverging_scale(Z_raw)
                cf = ax.contourf(X_smooth, Y_smooth, Z_smooth, levels=levels, norm=norm, cmap=_soft_red_blue_cmap(), extend="both")
                _set_contourf_edgecolor_face(cf)

                cbar = fig.colorbar(cf, ax=ax, ticks=ticks)
                cbar.ax.yaxis.set_major_formatter(ticker.FuncFormatter(_format_bof_tick))

                _safe_contour_labels(ax=ax, X=X_smooth, Y=Y_smooth, Z=Z_smooth, levels=line_levels, fmt_func=_format_bof_tick, linewidths=0.8, alpha=0.45, fontsize=20)

                if np.nanmin(Z_raw) <= 0.0 <= np.nanmax(Z_raw):
                    try:
                        cl0 = ax.contour(X_smooth, Y_smooth, Z_smooth, levels=[0.0], colors="black", linewidths=1.8, alpha=0.9)
                        ax.clabel(cl0, inline=True, inline_spacing=5, fontsize=20, fmt="0")
                    except Exception: pass
        else:
            levels = np.linspace(0.0, 1.0, 21)
            ticks = np.linspace(0.0, 1.0, 6)

            cf = ax.contourf(X_smooth, Y_smooth, Z_smooth, levels=levels, cmap="Blues", extend="neither")
            _set_contourf_edgecolor_face(cf)

            cbar = fig.colorbar(cf, ax=ax, ticks=ticks)
            cbar.ax.yaxis.set_major_formatter(ticker.FuncFormatter(_format_bof_tick))

            line_levels = np.linspace(0.0, 1.0, 6)
            _safe_contour_labels(ax=ax, X=X_smooth, Y=Y_smooth, Z=Z_smooth, levels=line_levels, fmt_func=lambda x, pos=None: f"{x:.1f}", linewidths=0.8, alpha=0.45, fontsize=9)

        ax.set_title(titles[key], fontsize=14)
        ax.set_xlabel(x_label, fontsize=12)
        ax.set_ylabel(r"$R_f$", fontsize=12)

    fig_title = f"2D best-of-fit across $w_{{ee}}$ and $R_f$ (k={target_k:.7f})" if metric == "delta_aic" else f"2D fit-winner map across $w_{{ee}}$ and $R_f$ (k={target_k:.7f})"
    fig.suptitle(fig_title, fontsize=18)

    if save:
        os.makedirs(plot_dir, exist_ok=True)
        suffix = "_wc" if show_wee_in_wc else ""
        out = os.path.join(plot_dir, f"bof_vs_wee_rf_2d_{metric}_k_{target_k:.7f}{suffix}.svg")
        fig.savefig(out, dpi=150, bbox_inches="tight")
        print(f"Saved: {out}")

    return fig, df, pivots


# ============================================================
# SIMULATION PROCESS WORKERS
# ============================================================

def simulate_plot_and_save_task_2d(args):
    params, Rf, wee, wee_norm, k, raw_data_dir, plot_dir = args

    filename = f"raw_data_2d_Rf_{Rf:.6f}_k_{k:.7f}_wee_{wee:.6f}.npz"
    save_path = os.path.join(raw_data_dir, filename)

    if os.path.exists(save_path):
        return save_path

    try:
        simulator = CANNSimulator2D(params)
        results = simulator.run_simulation(Rf=Rf, Wee=wee, k=k)

        Ue = np.asarray(results["E_bump_height"])
        z_list = np.asarray(results["E_stim"])
        T = len(Ue)
        time_array = np.arange(T) * params["time_step"]

        delta_z_mag = np.asarray(results["delta_z"])
        if delta_z_mag.ndim != 1:
            raise ValueError("results['delta_z'] must be a 1D jump-radius array.")
        if len(delta_z_mag) == T - 1:
            delta_z_mag = np.insert(delta_z_mag, 0, np.nan)
        elif len(delta_z_mag) != T:
            raise ValueError(f"delta_z length {len(delta_z_mag)} does not match T={T} or T-1.")

        delta_z_2d = results.get("delta_z_2d", None)
        if delta_z_2d is not None:
            delta_z_2d = np.asarray(delta_z_2d)
            if delta_z_2d.ndim != 2 or delta_z_2d.shape[1] != 2:
                raise ValueError("results['delta_z_2d'] must have shape (T,2) or (T-1,2).")
            if delta_z_2d.shape[0] == T - 1:
                delta_z_2d = np.vstack([np.array([[np.nan, np.nan]]), delta_z_2d])
            elif delta_z_2d.shape[0] != T:
                raise ValueError(f"delta_z_2d length {delta_z_2d.shape[0]} does not match T={T} or T-1.")

        generate_and_save_heatmaps(
            time=time_array,
            Oc=results.get("Oc"),
            params=params,
            Rf=Rf,
            wee=wee,
            k=k,
            output_dir=plot_dir,
        )

        save_dict = {
            "time": time_array,
            "z_list": z_list,
            "Ue_list": Ue,
            "delta_z_mag": delta_z_mag,
            "Rf": np.array(Rf),
            "k": np.array(k),
            "wee": np.array(wee),
        }
        if delta_z_2d is not None:
            save_dict["delta_z_2d"] = delta_z_2d

        np.savez_compressed(save_path, **save_dict)
        return save_path

    except Exception as e:
        print(f"Simulation Error for Rf={Rf:.6f}, k={k:.7f}: {e}")
        return None


def generate_and_save_heatmaps(time, Oc, params, Rf, wee, k, output_dir, win_bins=1):
    if Oc is None or len(Oc) == 0:
        return
        
    time_s = np.asarray(time)[:len(Oc)]
    time_dec = time_s[win_bins - 1:]
    
    fig_hm = plot_heatmaps(time_s, time_dec, Oc, win_bins, Rf, wee, k)

    os.makedirs(output_dir, exist_ok=True)
    fig_hm_path = os.path.join(output_dir, f"heatmaps_Rf_{Rf:.6f}_k_{k:.7f}_wee_{wee:.6f}.svg")
    fig_hm.savefig(fig_hm_path, dpi=150, bbox_inches="tight")
    
    plt.close(fig_hm)


def generate_simulation_data(params, target_wee, Rf_list, k ,n_jobs=-1, raw_data_dir='output_net/raw_npz/2d', plot_dir='output_net/plots/2d'):
    os.makedirs(raw_data_dir, exist_ok=True)
    os.makedirs(plot_dir, exist_ok=True)
    
    critical_weight = params.get('critical_weight', 1.0) 
    wee_norm = target_wee / critical_weight
    
    max_workers = mp.cpu_count() if n_jobs == -1 else n_jobs
    tasks = [(params, Rf, target_wee, wee_norm,k, raw_data_dir, plot_dir) for Rf in Rf_list]
    saved_paths = []
    
    print(f"🚀 Starting {len(tasks)} simulations across {max_workers} CPU cores...")
    
    with concurrent.futures.ProcessPoolExecutor(max_workers=max_workers) as executor:
        futures = [executor.submit(simulate_plot_and_save_task_2d, task) for task in tasks]
        for future in tqdm(concurrent.futures.as_completed(futures), total=len(futures), desc="Simulating"):
            try:
                res = future.result() 
                if res is not None:
                    saved_paths.append(res)
            except Exception as e:
                print(f"\n A worker process crashed during simulation: {e}")
                
    return saved_paths


def run_offline_analysis(data_paths, params, target_wee, k, n_jobs=-1, plot_dir="output_net/plots/2d/"):
    if not data_paths:
        print("No data paths provided for analysis.")
        return

    os.makedirs(plot_dir, exist_ok=True)
    max_workers = mp.cpu_count() if n_jobs == -1 else n_jobs

    tasks = [(path, params, plot_dir) for path in data_paths]
    results = []

    print(f" Analyzing {len(tasks)} datasets across {max_workers} CPU cores...")

    with concurrent.futures.ProcessPoolExecutor(max_workers=max_workers) as executor:
        futures = [executor.submit(analyze_saved_data_task_2d, task) for task in tasks]
        for future in tqdm(concurrent.futures.as_completed(futures), total=len(futures), desc="Analyzing 2D"):
            try:
                res = future.result()
                if res is not None:
                    results.append(res)
            except Exception as e:
                print(f"\nA worker process crashed during analysis: {e}")

    if results:
        summary_df = pd.DataFrame(results).sort_values(by=["Rf", "k"])
        csv_path = os.path.join(plot_dir, f"summary_2d_fixed_wee_{target_wee:.6f}_{k:.7f}.csv")
        summary_df.to_csv(csv_path, index=False)
        print(f"Summary CSV successfully saved to {csv_path}")
    else:
        print(" No valid results were returned to save into a CSV.")


def _simulate_for_k_2d(args):
    k_val, params, Rf, wee = args
    sim = CANNSimulator2D(params)
    res = sim.run_simulation(Rf=Rf, Wee=wee, k=k_val)
    
    return {
        'k_val': k_val,
        'E_bump_height': res['E_bump_height'],
        'Oc': res.get('Oc')
    }


# ============================================================
# MAIN EXECUTION
# ============================================================

if __name__ == "__main__":
    params = {
        'time_constant_exc': 1.0,
        'position_max': 180.0,
        'position_min': -180.0,
        'gaussian_width_exc': 40.0,
        'gaussian_width_ES': 20.0,
        'num_neurons': 180,
        'simulation_time':  500.0,
        'time_step': 0.1, 
        'recording_start': 50,
        'Fano_factor': 0.5,
        'normalization_k': 0.00005, 
        'inhibitory_gain': 10,
        'input_position': [0, 20],
        'feedforward_scale': 0.604679,
        't_steady': 50,
        'initial_mean_eq': 0,
        'initial_var_eq': 30,
        'initial_scale_eq': 1e-1
    }
    # Just comment out the other parts if you want a specific figure.
    simulator = CANNSimulator2D(params)
    target_wee = 0.4 * simulator.params['critical_weight']
    sim = CANNSimulator2D(params=params)
    wc = simulator.params['critical_weight']
    weelist = [0.1*wc, 0.2*wc, 0.3*wc, 0.4*wc, 0.5*wc, 0.6*wc, 0.7*wc, 0.8*wc, 0.9*wc, 1.0*wc] 
   
    Rf_list = np.linspace(0.01, 0.1, 5).tolist() + np.linspace(0.2, 1.0, 5).tolist() + np.linspace(2, 5.0, 5).tolist()
    Rf_list_diag = [0.01]

    # Figure 4EG & Appendix Figure 4A: example of bump height, tail slope and ccdf for one wee and one Rf with fitting 
    weelist = [target_wee] 
  
    df_tail = _collect_or_build_summary_2d_fixed_k(
        params=params,
        target_k=params['normalization_k'],
        Rf_list=Rf_list_diag,
        wee_list=weelist,
        raw_data_dir='output_net/raw_npz/2d/',
        plot_dir='output_net/plots/2d/',
        n_jobs=2,
        force_reanalyze=True,
        save_diagnostic_plot=True,
        save_summary_csv=False,
    )
    
    # Appendix Figure: 2D heatmaps of BOF metrics across wee and Rf
    plot_bof_vs_wee_rf_2d(
        params,
        target_k=0.00005,
        Rf_list=Rf_list,
        wee_list=weelist,
        n_jobs=2, 
        force_reanalyze=False,
        raw_data_dir="output_net/raw_npz/2d",
        plot_dir="output_net/plots/2d",
        metric="delta_aic",
        save=True,
        save_diagnostic_plot=False,
        show_wee_in_wc=True,
    )
    
    plot_tail_slope_vs_wee_rf_2d(
        params,
        target_k=0.00005,
        Rf_list=Rf_list,
        wee_list=weelist,
        n_jobs=2,
        force_reanalyze=False,
        raw_data_dir="output_net/raw_npz/2d",
        plot_dir="output_net/plots/2d", 
        save=True,
        save_diagnostic_plot=False,
        show_wee_in_wc=True,
    )
    
    sim = CANNSimulator2D(params=params)
    
    # Figure 4 B: Mean readout across centers
    n_points_per_axis = 10
    coords = np.linspace(-40, 40, n_points_per_axis)
    X, Y = np.meshgrid(coords, coords)
    evenly_spaced_centers = np.column_stack([X.ravel(), Y.ravel()])

    measure_readout_mean_2d(
        simulator=sim,
        centers_list=evenly_spaced_centers,
        n_trials=2,
        n_workers=4,
        save_plot=True,
    )
    
    target_wee = sim.params['critical_weight'] * 0.4
    
    # Figure 4 A: Precision
    calibrate_ff_scale_2d(sim, Wee=target_wee)
```
