## Supplementary Codes for "Scale-mixture Langevin sampling in the subspaces of recurrent cortical circuit dynamics": cann_2d.py

```
import os
import sys
import copy
import concurrent.futures
import multiprocessing as mp
from collections import defaultdict

import numpy as np
from tqdm import tqdm

import matplotlib.pyplot as plt
import matplotlib.animation as animation
import matplotlib.colors as mcolors
import seaborn as sns
from matplotlib.collections import LineCollection


class CANNSimulator2D:
    def __init__(self, params=None):
        default_params = {
            'time_constant_exc': 1.0,
            'position_max': 180.0,
            'position_min': -180.0,
            'gaussian_width_exc': 40.0,
            'gaussian_width_ES': 20.0,
            'Fano_factor': 0.5,
            'num_neurons': 40,
            'simulation_time': 100.0,
            'time_step': 0.1,
            'normalization_k': 0.0005,
            'inhibitory_gain': 10,
            'feedforward_scale': 0.604679,
            'recording_start': 20.0,
        }
        self.params = {**default_params, **(params or {})}
        self._compute_parameters()
        self.initialize_network()

    def _compute_parameters(self):
        p = self.params
        p['time_constant_som'] = p['time_constant_exc']
        p['gaussian_width_SE'] = np.sqrt(p['gaussian_width_exc'] ** 2 - p['gaussian_width_ES'] ** 2)
        p['gaussian_width_SS'] = np.sqrt((p['gaussian_width_exc'] ** 2 + p['gaussian_width_ES'] ** 2) / 2)
        p['position_range'] = p['position_max'] - p['position_min']
        p['neuron_density'] = (p['num_neurons'] ** 2) / (p['position_range'] ** 2)
        p['PrefStim'] = np.linspace(p['position_min'], p['position_max'], p['num_neurons'], endpoint=False)
        
        k = p['normalization_k']
        rho = p['neuron_density']
        a = p['gaussian_width_exc']
        numerator = 32 * np.pi * (a**2) * k
        p['critical_weight'] = np.sqrt(numerator / rho)
        
        p['recurrent_weight_e2e'] = 0.5 * p['critical_weight']
        p['recurrent_weight_e2i'] = 0.5 * p['critical_weight']
        p['feedforward_intensity_scale'] = p['critical_weight'] / (4 * np.pi * a**2 * k)

    def initialize_network(self):
        p = self.params
        xrange = p['position_range']
        PrefStim = p['PrefStim']
        
        X, Y = np.meshgrid(PrefStim, PrefStim)
        dist_x = np.angle(np.exp(1j * (X - PrefStim[0]) * (2 * np.pi / xrange))) * xrange / (2 * np.pi)
        dist_y = np.angle(np.exp(1j * (Y - PrefStim[0]) * (2 * np.pi / xrange))) * xrange / (2 * np.pi)
        dist_sq = dist_x**2 + dist_y**2
        
        def create_kernel(width):
            ker = np.exp(-dist_sq / (2 * width**2)) / (2 * np.pi * width**2)
            return np.fft.fft2(ker)

        self.WE = create_kernel(p['gaussian_width_exc'])
        self.WES = create_kernel(p['gaussian_width_ES'])
        self.WSE = create_kernel(p['gaussian_width_SE'])

    def _calculate_stimulus_2d(self, activity):
        if isinstance(activity, list):
            activity = np.array(activity)
        
        steps = activity.shape[0]
        decoded_path = np.zeros((steps, 2))
        
        pref = self.params['PrefStim']
        range_rad = 2 * np.pi / self.params['position_range']
        X, Y = np.meshgrid(pref, pref)
        exp_ix = np.exp(1j * X * range_rad)
        exp_iy = np.exp(1j * Y * range_rad)
        
        for t in range(steps):
            frame = activity[t]
            z_x = np.sum(frame * exp_ix)
            z_y = np.sum(frame * exp_iy)
            
            pos_x = np.angle(z_x) / range_rad if np.abs(z_x) > 1e-9 else 0
            pos_y = np.angle(z_y) / range_rad if np.abs(z_y) > 1e-9 else 0
            decoded_path[t] = [pos_x, pos_y]
            
        return decoded_path

    def run_simulation(
        self,
        Wee,
        Rf=None,
        k=None,
        show_progress=True,
        cue_on=0.0,
        cue_off=None,
        save_Oc=True,       
        record_dt=0.1,   
    ):
        p = self.params

        N = p['num_neurons']
        T = p['simulation_time']
        dt = p['time_step']
        steps = int(T / dt)
        a = p['gaussian_width_exc']
        rho = p['neuron_density']

        F = p["Fano_factor"]
        tauE = p['time_constant_exc']

        ff_scale = p['feedforward_scale']
        k_eff = p['normalization_k'] if k is None else k

        U = np.zeros((N, N))
        Oc = np.zeros((N, N))

        history = {'time': [], 'input_pos': [], 'cue_amp': [], 'E_stim': []}
        if save_Oc:
            history['Oc'] = []
            history['E_bump_height'] = []

        PrefStim = p['PrefStim']
        X_grid, Y_grid = np.meshgrid(PrefStim, PrefStim)
        xrange = p['position_range']
        dist_factor = 2 * np.pi / xrange
        dx_grid = xrange / N

        if cue_off is None:
            cue_off = T

        range_rad = 2 * np.pi / p['position_range']
        exp_ix = np.exp(1j * X_grid * range_rad)
        exp_iy = np.exp(1j * Y_grid * range_rad)

        save_interval = max(1, int(record_dt / dt))

        if show_progress:
            print(f"Starting simulation: {steps} steps, Rf={Rf if Rf is not None else 'Auto'}")
            iterator = tqdm(range(steps))
        else:
            iterator = range(steps)

        for i in iterator:
            t = i * dt
            mu_x, mu_y = p.get('input_position', [0, 0])

            dx = np.angle(np.exp(1j * (X_grid - mu_x) * dist_factor)) * xrange / (2 * np.pi)
            dy = np.angle(np.exp(1j * (Y_grid - mu_y) * dist_factor)) * xrange / (2 * np.pi)
            d2 = dx**2 + dy**2

            base_scale = Rf if Rf is not None else p['feedforward_intensity_scale']
            cue_amp = base_scale if (cue_on <= t < cue_off) else 0.0

            Iext = cue_amp * np.exp(-d2 / (4 * a**2))

            fO = np.fft.fft2(Oc)
            Rec_EE = np.real(np.fft.ifft2(self.WE * fO))

            drift = (-U + Wee * Rec_EE + ff_scale * Iext) * (dt / tauE)
            diffusion = np.sqrt(F * np.clip(U, 0.0, None) * (dt / tauE)) * np.random.normal(0, 1, (N, N))

            U += drift + diffusion
            U = np.maximum(U, 0)

            sq_U = U**2
            denom = 1 + k_eff * rho * np.sum(sq_U) * dx_grid**2
            Oc = sq_U / denom

            if i % save_interval == 0:
                z_x = np.sum(Oc * exp_ix)
                z_y = np.sum(Oc * exp_iy)
                pos_x = np.angle(z_x) / range_rad if np.abs(z_x) > 1e-9 else 0
                pos_y = np.angle(z_y) / range_rad if np.abs(z_y) > 1e-9 else 0

                history['E_stim'].append([pos_x, pos_y])
                history['time'].append(t)
                history['input_pos'].append((mu_x, mu_y))
                history['cue_amp'].append(cue_amp)

                if save_Oc:
                    history['Oc'].append(Oc.copy())
                    history['E_bump_height'].append(np.sum(Oc) * dx_grid**2 / (2 * np.pi * a**2))

        if save_Oc:
            history['delta_z_2d'], history['delta_z'] = self._calculate_stimulus_change_2d(history['Oc'])
            
        history['params'] = copy.deepcopy(self.params)

        return history

    def _calculate_stimulus_change_2d(self, activity):
        if isinstance(activity, list):
            activity = np.array(activity)
            
        steps = activity.shape[0]
        pref = self.params['PrefStim']
        range_rad = 2 * np.pi / self.params['position_range']
        
        X, Y = np.meshgrid(pref, pref)
        exp_ix = np.exp(1j * X * range_rad)
        exp_iy = np.exp(1j * Y * range_rad)
        
        e_x = np.sum(activity * exp_ix, axis=(1, 2))
        e_y = np.sum(activity * exp_iy, axis=(1, 2))
        
        def normalize_complex(e):
            magnitudes = np.abs(e)
            normalized_e = np.zeros_like(e, dtype=complex)
            non_zero_mask = magnitudes > 0
            normalized_e[non_zero_mask] = e[non_zero_mask] / magnitudes[non_zero_mask]
            return normalized_e

        z_x = normalize_complex(e_x)
        z_y = normalize_complex(e_y)
        
        delta_zx_complex = z_x[1:] * np.conj(z_x[:-1])
        delta_zy_complex = z_y[1:] * np.conj(z_y[:-1])
        
        delta_x = np.angle(delta_zx_complex) / range_rad
        delta_y = np.angle(delta_zy_complex) / range_rad
        
        delta_z_2d = np.column_stack((delta_x, delta_y))
        delta_z = np.linalg.norm(delta_z_2d, axis=1)
        
        return delta_z_2d, delta_z

    def _wrap_periodic_2d(self, x):
        L = self.params['position_range']
        return (np.asarray(x) + 0.5 * L) % L - 0.5 * L

    def get_bump_position_2d(self, Rf, Wee=None, k=None):
        if Wee is None:
            Wee = self.params['recurrent_weight_e2e']

        result = self.run_simulation(
            Wee=Wee,
            Rf=Rf,
            k=k,
            show_progress=False,
        )
        return np.asarray(result['E_stim'])

    def _trial_precision_from_path_2d(self, path, mode="axis_mean", eps=1e-12):
        t_record = self.params['recording_start']
        dt = self.params['time_step']
        start_idx = int(t_record / dt)

        path = np.asarray(path)[start_idx:]
        target = np.asarray(self.params['input_position'], dtype=float)

        disp = self._wrap_periodic_2d(path - target[None, :])

        var_x = np.var(disp[:, 0])
        var_y = np.var(disp[:, 1])

        var_x = max(var_x, eps)
        var_y = max(var_y, eps)

        if mode == "axis_mean":
            return 0.5 * (1.0 / var_x + 1.0 / var_y)
        elif mode == "trace":
            return 2.0 / (var_x + var_y)
        else:
            raise ValueError(f"Unknown mode: {mode}")


def _wrap_periodic_2d_static(x, position_range):
    return (np.asarray(x) + 0.5 * position_range) % position_range - 0.5 * position_range


def _trial_precision_from_path_2d_static(
    path,
    params,
    time_array, 
    mode="axis_mean",
    eps=1e-12,
    min_samples=10,
):
    t_record = params["recording_start"]
    path = np.asarray(path, dtype=float)
    time_array = np.asarray(time_array, dtype=float)

    if path.ndim != 2 or path.shape[1] != 2:
        return np.nan

    mask = time_array >= t_record
    path = path[mask]

    finite_mask = np.isfinite(path).all(axis=1)
    path = path[finite_mask]

    if len(path) < min_samples:
        return np.nan

    target = np.asarray(params["input_position"], dtype=float).reshape(2,)
    disp = _wrap_periodic_2d_static(
        path - target[None, :],
        params["position_range"],
    )

    if not np.isfinite(disp).all():
        return np.nan

    cov = np.cov(disp.T, ddof=1)
    if cov.shape != (2, 2) or not np.isfinite(cov).all():
        return np.nan

    var_x = max(float(cov[0, 0]), eps)
    var_y = max(float(cov[1, 1]), eps)

    if mode == "axis_mean":
        return 0.5 * (1.0 / var_x + 1.0 / var_y)
    elif mode == "trace":
        return 2.0 / (var_x + var_y)
    elif mode == "full":
        cov_reg = cov + eps * np.eye(2)
        try:
            prec = np.linalg.inv(cov_reg)
        except np.linalg.LinAlgError:
            return np.nan
        return 0.5 * float(np.trace(prec))
    else:
        raise ValueError(f"Unknown mode: {mode}")


def _calibration_worker_2d(task):
    rf_idx = task["rf_idx"]
    Rf = task["Rf"]
    params = copy.deepcopy(task["params"])
    Wee = task["Wee"]
    k = task["k"]
    precision_mode = task["precision_mode"]

    sim = CANNSimulator2D(params=params)

    results = sim.run_simulation(
        Wee=Wee,
        Rf=Rf,
        k=k,
        show_progress=False,
        save_Oc=False, 
    )

    path = np.asarray(results["E_stim"], dtype=float)
    time_array = np.asarray(results["time"], dtype=float)
    
    precision = _trial_precision_from_path_2d_static(
        path,
        sim.params,
        time_array=time_array, 
        mode=precision_mode,
    )
    return rf_idx, precision


def _center_worker_2d(task):
    """
    Top-level worker function executed by ProcessPoolExecutor.
    Runs a single simulation trial for a specific input center.
    """
    center_idx = task["center_idx"]
    params = task["params"]
    
    sim = CANNSimulator2D(params)
    res = sim.run_simulation(
        Wee=params.get('recurrent_weight_e2e', 0.5 * sim.params['critical_weight']),
        Rf=params.get('Rf', None),
        show_progress=False, 
        save_Oc=False
    )
    
    path = np.asarray(res["E_stim"], dtype=float)
    t_record = params.get("recording_start", 20.0)
    dt = params.get("time_step", 0.1)
    start_idx = int(t_record / dt)
    
    if len(path) > start_idx:
        readout = np.nanmean(path[start_idx:], axis=0)
    else:
        readout = np.array([np.nan, np.nan])
    
    return center_idx, readout


def _run_center_sweep_2d(
    params,
    centers_list,
    n_trials,
    max_workers=None,
    desc="Simulating centers",
):
    tasks = []
    for center_idx, center in enumerate(centers_list):
        for _ in range(n_trials):
            trial_params = copy.deepcopy(params)
            trial_params["input_position"] = center 
            
            tasks.append({
                "center_idx": center_idx,
                "center": center,
                "params": trial_params,
            })

    readouts_by_idx = defaultdict(list)
    failed_workers = 0

    print(f"Starting {len(tasks)} trials across {max_workers} workers...")

    with concurrent.futures.ProcessPoolExecutor(max_workers=max_workers) as executor:
        futures = [executor.submit(_center_worker_2d, task) for task in tasks]

        for future in tqdm(
            concurrent.futures.as_completed(futures),
            total=len(futures),
            desc=desc,
        ):
            try:
                center_idx, readout = future.result()
                if np.all(np.isfinite(readout)):
                    readouts_by_idx[center_idx].append(np.asarray(readout, dtype=float))
            except Exception as e:
                failed_workers += 1
                print(f"\nA worker process crashed during simulation: {repr(e)}")

    n_centers = len(centers_list)
    dim = 2
    for idx in range(n_centers):
        if len(readouts_by_idx[idx]) > 0:
            dim = len(readouts_by_idx[idx][0])
            break

    mean_readouts = np.full((n_centers, dim), np.nan)
    n_valid_by_idx = np.zeros(n_centers, dtype=int)

    for center_idx in range(n_centers):
        vals = readouts_by_idx.get(center_idx, [])
        n_valid_by_idx[center_idx] = len(vals)

        if len(vals) > 0:
            mean_readouts[center_idx] = np.mean(vals, axis=0)

    return (
        mean_readouts,
        readouts_by_idx,  
        n_valid_by_idx,
        failed_workers,
    )


def _run_precision_sweep_2d(
    params,
    Rf_list,
    n_trials,
    Wee,
    k,
    precision_mode,
    max_workers=None,
    desc="Simulating",
):
    tasks = []
    for rf_idx, Rf in enumerate(Rf_list):
        for _ in range(n_trials):
            tasks.append({
                "rf_idx": rf_idx,
                "Rf": float(Rf),
                "params": copy.deepcopy(params),
                "Wee": float(Wee),
                "k": None if k is None else float(k),
                "precision_mode": precision_mode,
            })

    precision_by_idx = defaultdict(list)
    failed_workers = 0

    print(f"Starting {len(tasks)} calibration trials across {max_workers} workers...")

    with concurrent.futures.ProcessPoolExecutor(max_workers=max_workers) as executor:
        futures = [executor.submit(_calibration_worker_2d, task) for task in tasks]

        for future in tqdm(
            concurrent.futures.as_completed(futures),
            total=len(futures),
            desc=desc,
        ):
            try:
                rf_idx, precision = future.result()
                if np.isfinite(precision):
                    precision_by_idx[rf_idx].append(float(precision))
            except Exception as e:
                failed_workers += 1
                print(f"\nA worker process crashed during simulation: {repr(e)}")

    mean_precision = []
    std_precision = []
    n_valid_by_idx = []

    for rf_idx in range(len(Rf_list)):
        vals = precision_by_idx.get(rf_idx, [])
        n_valid_by_idx.append(len(vals))

        if len(vals) == 0:
            mean_precision.append(np.nan)
            std_precision.append(np.nan)
        elif len(vals) == 1:
            mean_precision.append(vals[0])
            std_precision.append(0.0)
        else:
            mean_precision.append(float(np.mean(vals)))
            std_precision.append(float(np.std(vals, ddof=1)))

    return (
        np.asarray(mean_precision, dtype=float),
        np.asarray(std_precision, dtype=float),
        precision_by_idx,
        np.asarray(n_valid_by_idx, dtype=int),
        failed_workers,
    )


def _fit_slope_through_origin(x, y):
    x = np.asarray(x, dtype=float)
    y = np.asarray(y, dtype=float)

    mask = np.isfinite(x) & np.isfinite(y)
    x = x[mask]
    y = y[mask]

    if len(x) < 2:
        return np.nan

    denom = np.dot(x, x)
    if denom <= 0:
        return np.nan

    return float(np.dot(x, y) / denom)


def calibrate_ff_scale_2d(
    simulator,
    Rf_list=np.linspace(0.5, 10, 10),
    Wee=0.4,
    n_trials=20,
    k=None,
    precision_mode="axis_mean",
    theoretical_slope=None,
    n_workers=None,
    save_plot=True,
):
    rho = simulator.params["neuron_density"]

    if theoretical_slope is None:
        theoretical_slope = 2.0 * np.pi * rho

    if k is None:
        k = simulator.params["normalization_k"]

    if n_workers is None:
        n_workers = min(10, max(1, os.cpu_count() // 4))

    old_ff = float(simulator.params["feedforward_scale"])
    Rf_list = np.asarray(Rf_list, dtype=float)

    base_params = copy.deepcopy(simulator.params)

    (
        mean_precision,
        std_precision,
        _,
        n_valid_by_idx,
        failed_workers,
    ) = _run_precision_sweep_2d(
        params=base_params,
        Rf_list=Rf_list,
        n_trials=n_trials,
        Wee=Wee,
        k=k,
        precision_mode=precision_mode,
        max_workers=n_workers,
        desc=f"Running {n_trials} trials per Rf (2D)",
    )

    valid_mask = np.isfinite(mean_precision)

    if valid_mask.sum() < 2:
        raise RuntimeError(
            "Not enough successful calibration trials to fit slope.\n"
            f"valid counts per Rf: {n_valid_by_idx.tolist()}\n"
            f"mean_precision: {mean_precision.tolist()}\n"
            f"failed worker futures: {failed_workers}"
        )

    slope = _fit_slope_through_origin(Rf_list[valid_mask], mean_precision[valid_mask])

    if slope <= 0 or not np.isfinite(slope):
        raise RuntimeError(
            f"Invalid fitted slope: {slope}\n"
            f"Rf(valid)={Rf_list[valid_mask].tolist()}\n"
            f"precision(valid)={mean_precision[valid_mask].tolist()}"
        )

    new_ff = old_ff * (theoretical_slope / slope)
    simulator.params["feedforward_scale"] = float(new_ff)

    updated_params = copy.deepcopy(simulator.params)

    (
        mean_precision_new,
        std_precision_new,
        _,
        n_valid_by_idx_new,
        failed_workers_new,
    ) = _run_precision_sweep_2d(
        params=updated_params,
        Rf_list=Rf_list,
        n_trials=n_trials,
        Wee=Wee,
        k=k,
        precision_mode=precision_mode,
        max_workers=n_workers,
        desc="Recalculating precision with updated ff_scale (2D)",
    )

    slope_new = _fit_slope_through_origin(Rf_list, mean_precision_new)

    if save_plot:
        sim_data = {"Rf_list": Rf_list, "mean_precision_new": mean_precision_new, "std_precision_new": std_precision_new,
                    "theoretical_slope": theoretical_slope, "slope_new": slope_new, "n_trials": n_trials, "precision_mode": precision_mode}
        sim_data_filename = f"precision_calibration_2d_ff{new_ff:.5f}_{precision_mode}.npz"
        np.savez(sim_data_filename, **sim_data)
        print(f"Saved precision calibration data to {sim_data_filename}")

        plt.figure(figsize=(10, 6))
        plt.plot(Rf_list, theoretical_slope * Rf_list, label="Theoretical prediction")

        valid_plot = np.isfinite(mean_precision_new)
        plt.errorbar(
            Rf_list[valid_plot],
            mean_precision_new[valid_plot],
            yerr=std_precision_new[valid_plot],
            fmt="o",
            capsize=3,
            label=f"Simulation ({n_trials} trials / Rf)",
        )

        plt.xlabel("Feedforward intensity (Rf)")
        plt.ylabel("2D precision")
        plt.title(f"2D precision calibration ({precision_mode})")
        plt.legend()
        plt.grid(False)
        plt.savefig(f"precision_vs_Rf_2d_ff{new_ff:.5f}_{precision_mode}.svg", bbox_inches="tight")
        plt.close()

    print(f"old feedforward_scale   = {old_ff:.6f}")
    print(f"first-pass fitted slope = {slope:.6f}")
    print(f"target slope            = {theoretical_slope:.6f}")
    print(f"new feedforward_scale   = {new_ff:.6f}")
    print(f"second-pass slope       = {slope_new:.6f}")
    print(f"valid counts per Rf     = {n_valid_by_idx_new.tolist()}")
    print(f"failed worker futures   = {failed_workers_new}")

    return new_ff


def plot_2d_cann_results(results, simulator):
    times = np.array(results['time'])
    decoded_path = np.array(results['E_stim'])
    
    recording_start = simulator.params.get('recording_start', 50.0)
    start_mask = times >= recording_start
    
    times = times[start_mask]
    decoded_path = decoded_path[start_mask]
    
    if len(times) == 0:
        print("Error: No data left after burn-in slicing.")
        return

    plt.figure(figsize=(10, 6)) 
    ax = plt.gca()

    mean_pos = np.mean(decoded_path, axis=0)
    cov_matrix = np.cov(decoded_path, rowvar=False)

    x_min, x_max = np.min(decoded_path[:, 0]), np.max(decoded_path[:, 0])
    y_min, y_max = np.min(decoded_path[:, 1]), np.max(decoded_path[:, 1])
    
    pad_x, pad_y = (x_max - x_min) * 1.2, (y_max - y_min) * 1.2
    grid_res = 250
    X, Y = np.meshgrid(np.linspace(x_min - pad_x, x_max + pad_x, grid_res),
                       np.linspace(y_min - pad_y, y_max + pad_y, grid_res))
    
    pos = np.empty(X.shape + (2,))
    pos[:, :, 0] = X
    pos[:, :, 1] = Y

    diff = pos - mean_pos
    inv_cov = np.linalg.inv(cov_matrix)
    Z = np.exp(-0.5 * np.sum(np.dot(diff, inv_cov) * diff, axis=2))
    Z = Z / np.max(Z) * 100
    
    heatmap = ax.pcolormesh(X, Y, Z, cmap='Blues', alpha=0.95, shading='gouraud', zorder=1, rasterized=True)
    
    cbar_heat = plt.colorbar(heatmap, ax=ax, pad=0.02)
    cbar_heat.set_label('Posterior Probability (%)', rotation=270, labelpad=15)

    tau = simulator.params.get('time_constant_exc', 1.0) 
    time_window = 10 * tau 
    t_end = times[-1]
    tail_mask = times >= (t_end - time_window)
    
    times_tail = times[tail_mask]
    path_tail = decoded_path[tail_mask]

    if len(times_tail) < 2:
        print("Warning: Not enough data points in the last 10 tau to draw a line.")
        return

    points = np.array([path_tail[:, 0], path_tail[:, 1]]).T.reshape(-1, 1, 2)
    segments = np.concatenate([points[:-1], points[1:]], axis=1)

    norm_val = plt.Normalize(times_tail.min(), times_tail.max())
    
    lc = LineCollection(segments, cmap='viridis', norm=norm_val, zorder=2)
    lc.set_array(times_tail[:-1])
    lc.set_linewidth(2.0) 
    lc.set_alpha(0.85) 
    
    line = ax.add_collection(lc)
    ax.autoscale() 

    cbar_line = plt.colorbar(line, ax=ax, pad=0.02)
    cbar_line.set_label('Time (ms)', rotation=270, labelpad=15)
        
    plt.title(r'2D Bump Trajectory (Last $10\tau$) & Posterior Heatmap')
    plt.xlabel('Position X')
    plt.ylabel('Position Y')
    plt.axis('equal') 
    
    cx, cy = mean_pos[0], mean_pos[1]
    std_x = np.sqrt(cov_matrix[0, 0])
    std_y = np.sqrt(cov_matrix[1, 1])
    
    radius_x = max(abs(x_max - cx), abs(cx - x_min), 3 * std_x)
    radius_y = max(abs(y_max - cy), abs(cy - y_min), 3 * std_y)
    
    plot_radius = max(radius_x, radius_y) * 1.05
    
    plt.xlim(cx - plot_radius, cx + plot_radius)
    plt.ylim(cy - plot_radius, cy + plot_radius)
    
    plt.tight_layout()
    plt.savefig('trajectory_heatmap_10tau.svg')
    plt.close()


def measure_readout_mean_2d(
    simulator,
    centers_list,
    n_trials=20,
    n_workers=None,
    save_plot=True,
):
    if n_workers is None:
        n_workers = min(10, max(1, os.cpu_count() // 4))

    ff_scale = float(simulator.params["feedforward_scale"])
    centers_list = np.asarray(centers_list, dtype=float)

    base_params = copy.deepcopy(simulator.params)

    sweep_results = _run_center_sweep_2d(
        params=base_params,
        centers_list=centers_list,
        n_trials=n_trials,
        max_workers=n_workers,
        desc=f"Running {n_trials} trials per center (2D)",
    )
    
    mean_readouts = sweep_results[0]
    n_valid_by_idx = sweep_results[-2] 
    failed_workers = sweep_results[-1]

    if save_plot:
        sim_data = {
            "centers_list": centers_list,
            "mean_readouts": mean_readouts,
            "n_trials": n_trials,
            "ff_scale": ff_scale
        }
        
        sim_data_filename = f"mean_across_centers_2d_ff{ff_scale:.5f}.npz"
        np.savez(sim_data_filename, **sim_data)
        print(f"Saved mean data to {sim_data_filename}")

        plt.figure(figsize=(10, 6))
        
        x_axis = centers_list if centers_list.ndim == 1 else np.linalg.norm(centers_list, axis=1)
        y_axis = mean_readouts if mean_readouts.ndim == 1 else np.linalg.norm(mean_readouts, axis=1)

        valid_plot = np.isfinite(y_axis)
        
        plt.plot(
            x_axis[valid_plot],
            y_axis[valid_plot],
            marker="o",
            linestyle="-",
            label=f"Mean Readout ({n_trials} trials/center)",
        )

        plt.plot(
            x_axis[valid_plot], 
            x_axis[valid_plot], 
            linestyle="--", 
            color="gray", 
            label="Ideal (Readout = Center)"
        )

        plt.xlabel("Input Center Magnitude / Position")
        plt.ylabel("Mean Readout Magnitude / Position")
        plt.title(f"2D Mean Readout vs. Input Center (ff_scale={ff_scale:.3f})")
        plt.legend()
        plt.grid(True, linestyle="--", alpha=0.6)
        
        plt.savefig(
            f"mean_vs_center_2d_ff{ff_scale:.5f}.svg",
            bbox_inches="tight",
        )
        plt.close()

    print(f"feedforward_scale used  = {ff_scale:.6f}")
    print(f"valid counts per center = {np.array(n_valid_by_idx).tolist()}")
    if failed_workers:
        print(f"failed worker futures   = {failed_workers}")

    return mean_readouts


# ============================================================
# MAIN EXECUTION
# ============================================================

if __name__ == "__main__":
    params = {
        'time_constant_exc': 1.0,
        'position_max': 180.0,
        'position_min': -180.0,
        'gaussian_width_exc': 40.0,
        'gaussian_width_ES': 20.0,
        'num_neurons': 180,
        'simulation_time':  500.0,
        'time_step': 0.1,
        'recording_start': 50.0,
        'Fano_factor': 0.5,
        'normalization_k': 0.00005,
        'inhibitory_gain': 10,
        'input_position': [0, 20],
        'feedforward_scale': 0.604679,
        't_steady': 50.0,
        'initial_mean_eq': 0,
        'initial_var_eq': 30,
        'initial_scale_eq': 1e-1
    }

    sim = CANNSimulator2D(params=params)
    
    # Figure 4 B: Mean readout across centers
    n_points_per_axis = 10
    coords = np.linspace(-40, 40, n_points_per_axis)
    X, Y = np.meshgrid(coords, coords)
    evenly_spaced_centers = np.column_stack([X.ravel(), Y.ravel()])

    measure_readout_mean_2d(
        simulator=sim,
        centers_list=evenly_spaced_centers,
        n_trials=2,
        n_workers=4,
        save_plot=True,
    )
    
    target_wee = sim.params['critical_weight'] * 0.4
    
    # Figure 4 A: Precision
    calibrate_ff_scale_2d(sim, Wee=target_wee)
```
