## Supplementary Codes for "Scale-mixture Langevin sampling in the subspaces of recurrent cortical circuit dynamics": README.md

## Long tail sampling with CANN Simulation

This repository provides Python scripts for simulating and analyzing Continuous Attractor Neural Networks (CANNs) in both one-dimensional (1D) and two-dimensional (2D) configurations. 

---

## Directory Structure

```bash
.
├── 1d/
│   ├── 1d.py                 # Main script for 1D CANN simulation
│   └──cann_stimulator.py    # CANN class definition
├── 2d/
│   ├── 2d.py                 # Main script for 2D CANN simulation
│   └── cannND.py             # cannND class definition (coupled 2D model)
└── README.md                 # This file (example)
```

---

## Quick Start 🚀

For a streamlined execution, it is highly recommended to run the main simulation scripts for each dimension. These scripts are designed with sequential execution of specific code blocks in mind for optimal performance and time-saving.

**For 1D Simulation:**
Navigate to the `1d` directory and execute the lines within `1d.py` sequentially as indicated in the file.

**For 2D Simulation:**
Navigate to the `2d` directory and execute the lines within `2d.py` sequentially as indicated in the file.


**Note:** Detailed instructions for running specific parts of the simulations are commented within `1d.py` and `2d.py`.

---

## Manual Workflow for 1D & 2D CANN Simulation

If you prefer to understand or run specific components individually:

1.  **1D CANN Simulation & Analysis**
    * Navigate to the `1d/` directory.
    * **Run Simulation:** Execute the simulation parts of `1d.py`.
        * This utilizes `cann_stimulator.py` for the network model.

2.  **2D CANN Simulation & Analysis**
    * Navigate to the `2d/` directory.
    * **Run Simulation:** Execute the simulation parts of `2d.py`.
        * This utilizes `cannND.py` for the 2D network model.


## Requirements

- The scipt is run in python 3.11. Normally as long as python>=3.8, the script is good to run.
=======
##
